# Comprehensive architecture of the bacterial RNA interactome

**DOI:** 10.1101/2025.09.11.675593

**Authors:** Gaurav Dugar

## Abstract

RNA-based regulation pervades bacteria, but obtaining a transcriptome-wide map of native RNA contacts has been technically challenging. Here TRIC-seq (Total RNA Interaction Capture) is introduced as an in situ, genetics-free proximity-ligation approach that preserves cellular context and resolves both intramolecular (structure) and intermolecular (regulatory) RNA contacts at high resolution and specificity. In *Escherichia coli*, TRIC-seq captures thousands of unique interactions, recovering known small RNA (sRNA) regulons and revealing non-canonical contacts among rRNAs, tRNAs and mRNAs, including a widespread interaction between the 3′ extension of 16S rRNA and 5’UTRs of stress-response mRNAs. Unsupervised analysis exposes a highly modular interactome organized by sRNA hubs. Beyond *E. coli*, TRIC-seq maps interactomes in diverse bacteria, enabling de novo discovery of novel sRNAs along with their targets. Finally, TRIC-seq uncovers a large, functionally coherent cohort of stress-related mRNAs that co-aggregate and are depleted for ribosome contacts.

## Introduction

RNA molecules are not merely passive messengers but key regulators of gene expression that orchestrate complex cellular responses. In bacteria, this is exemplified by sRNAs that modulate translation and mRNA stability, often with the help of the RNA chaperones like Hfq^1^. Beyond these canonical interactions, a growing body of evidence points to a richer regulatory RNA world, involving RNA–RNA contacts among diverse RNA classes.

In eukaryotes, stress can trigger assembly of stress granules—dynamic, non-membranous aggregates of non-translating mRNAs and RNA-binding proteins (RBPs) formed by phase separation^2,3^. Although bacteria lack organelles, they exhibit analogous behaviours, including formation of bacterial ribonucleoprotein bodies (BR-bodies)^4,5^ and condensation of RNA polymerase into foci^6^. Moreover, highly multiplexed spatial transcriptomics now shows that many bacterial transcripts are subcellularly localised^7,8^, suggesting that co-localised RNAs could form localised regulatory networks.

Despite this context, mapping native RNA-RNA contacts in prokaryotes remains difficult. Several high-throughput proximity-ligation approaches - PARIS, RIC-seq, and KARR-seq were developed for mammalian cells and combine crosslinking with proximity ligation to recover RNA duplexes transcriptome-wide, enabling RNA structure mapping at scale^9–12^. However, these methods predominantly detect intermolecular interactions among abundant RNAs. In bacteria, RNA crowding is at least an order of magnitude higher than mammalian cells: thousands of mRNAs plus tens of thousands of rRNAs/tRNAs are confined to a ∼1 fL cell volume^7,13,14^, which increases stochastic encounters and makes it harder to discriminate true biological contacts from artifactual ligations.

Pull-down based strategies like RIL-seq, iRIL-seq, CLASH and MAPS have successfully delineated sRNA-mRNA interactions associated with specific RBPs (e.g., Hfq) or with individual bait RNAs in bacteria^15–18^. Yet, these approaches capture the interactome fraction bound to the chosen factor, require genetic tractability, and are limited by the non-conserved nature of many RBPs and sRNAs, hindering de novo sRNA regulon discovery in non-model microbes. Moreover, tertiary structures of full-length bacterial mRNAs are not available as current transcriptome-wide methods report secondary structures^19^. Therefore, a comprehensive method is needed to capture the RNA interactome while simultaneously resolving tertiary structures of RNAs across the microbial domain.

TRIC-seq (Total RNA Interaction Capture) is presented here as an in situ method that maps native RNA-RNA interactions and structures across bacterial transcriptomes. The protocol uses a three-step fixation consisting of methanol, formaldehyde, and the amine reactive crosslinker dithiobis succinimidyl propionate (DSP). This crosslinking strategy is adapted from the high-resolution chromosome capture (DNA-DNA) method Micro-C-XL^20^ in eukaryotes and is further reinforced by recent *E. coli* Micro-C maps^21^, which achieved ultra-high resolution chromosome maps by adding an amine-reactive crosslinker after formaldehyde fixation. After fixation, proximal RNA fragments are ligated in-cell before lysis, which minimizes post lysis artifacts and preserves native spatial context.

Building on the feasibility of complex enzymology in permeabilized bacteria (e.g., split-pool scRNA-seq^22^), TRIC-seq performs an in situ workflow: MNase fragmentation to generate random RNA fragments, followed by 3′-end pCp-biotin labelling and proximity ligation essentially as done in RIC-seq^11^ to produce chimeras that report both intramolecular (structure) and intermolecular (regulatory) contacts (**Fig. 1a**). By coupling in-cell ligation chemistry with a null model that accounts for interaction frequency of each RNA, TRIC-seq resolves specific trans-interactions for the majority of expressed RNAs, thereby delivering a comprehensive, transcriptome-wide RNA interactome in bacteria.

**Figure 1.**
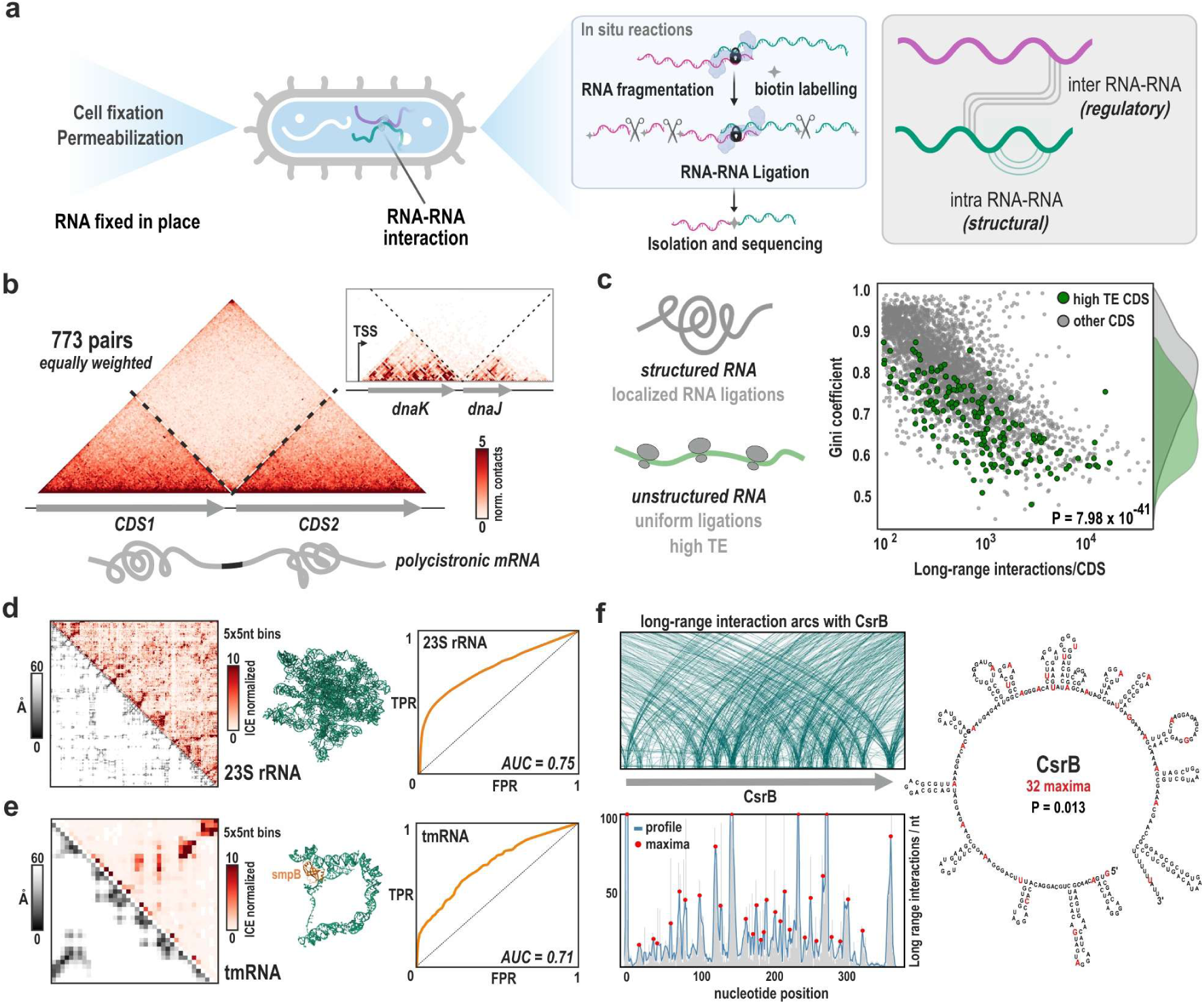
TRIC-seq workflow and validation of structural mapping. **a**, Overview of the TRIC-seq pipeline. Cellular RNAs are fixed in situ to preserve their native interactions and localisation. Following permeabilization, a series of enzymatic reactions, including RNA fragmentation, 3’-end biotin labelling, and proximity ligation, are performed within the intact cell. The resulting chimeric RNAs, which represent both intramolecular (structural) and intermolecular (regulatory) contacts, are then isolated for paired- end sequencing. **b,** Metaplot of TRIC-seq contacts across 773 consecutive CDS pairs from polycistronic transcripts. The plot reveals that individual CDSs fold into discrete structural domains with limited interaction across the intergenic region (dashed lines), a principle exemplified by the inset showing the contact map for the *dnaK-dnaJ* operon. **c,** A schematic illustrates that unstructured RNAs, often with high translation efficiency (TE), have more uniform ligations. The scatter plot shows that the top 200 genes with highest TE^24^ (green dots) have significantly lower Gini coefficients, indicating more open structure consistent with active translation. **d,** Benchmark on 23S rRNA. Left: ICE-normalized contact map (5x5 nt bins) in comparison to its complex tertiary structure. Right: ROC curve comparing TRIC-seq contact strength against nucleotide proximity in the known crystal structure, yielding an AUC of 0.75. **e,** Same analysis performed on tmRNA, which also shows a high concordance between the TRIC-seq contact map and the known structure. **f,** Left: IGV snapshot and quantification of long-range interaction arcs originating from the CsrB locus. Right: The predicted secondary structure of CsrB RNA with the maxima of long-range interaction highlighted in red. A one-tailed hypergeometric test confirms that the maxima are significantly enriched (P = 0.013) in accessible regions.

## Results

### TRIC-seq resolves native RNA structures

TRIC-seq was performed in several replicates on *E. coli* cultures grown to stationary phase, a condition of high regulatory activity where most sRNAs are known to be highly active^23^. After filtering and mapping, approximately 15 million unique chimeric reads (chimeras) were recovered in total. Each chimera represents a unique RNA-RNA proximity ligation event, providing a deep view of the RNA interactome (**Supplementary Table 1**). To evaluate the dataset, comparisons were performed to assess alignment with established transcriptome-wide methods like ribosome profiling and DMS-seq in *E. coli*^24^.

Chimeric reads were visualized using contact heat maps to infer local transcriptome structure. A meta-analysis of 773 consecutive CDS (coding sequence) pairs within polycistronic mRNA transcripts revealed a general principle of bacterial mRNA architecture: individual CDSs fold into discrete, semi-independent structural domains. This is exemplified by the *dnaK–dnaJ* heat-shock operon, where a sharp discontinuity in interaction frequency is observed, indicating structural insulation between the two CDSs (**Fig. 1b, Supplementary Fig. 1a**). Interactions were sharply confined within operon boundaries, with contacts between neighbouring transcripts being rare (**Supplementary Fig. 1b**). This modular, ORF-centric organisation is consistent with recent findings that such structures are a key determinant of translation efficiency (TE)^24^. To explore this link functionally, a Gini coefficient was calculated for the long-range intramolecular contacts of each CDS. The reasoning was that an unfolded, actively translating mRNA would present more uniformly distributed single-stranded regions accessible for ligation during TRIC-seq protocol, resulting in a low Gini coefficient. Indeed, a strong inverse correlation was found between the Gini coefficient and TE (**Fig. 1c, Supplementary Fig. 2**), using a set of high-TE genes previously defined using ribosome profiling^24,25^. This establishes that TRIC-seq provides a quantitative, transcriptome-wide readout of an RNA’s biophysical state and its engagement with the translational machinery.

The Hi-C-like contact maps generated by TRIC-seq also report on spatial proximities, providing direct information on the average tertiary structure of RNA molecules. This capability was validated by examining the contact map of several housekeeping RNAs (hkRNAs). TRIC-seq recapitulated the elaborate structures of ribosomal RNAs (**Fig. 1d; Supplementary Fig. 3)** and resolved the compact fold of transfer-messenger RNA, essential for ribosome rescue (**Fig. 1e**). This matches results from PARIS, RIC-seq and KARR-seq for mammalian ribosomes^12^, demonstrating that TRIC-seq can accurately map the average tertiary folds of RNAs in their native state. Apart from known structures, contact maps of individual mRNAs revealed a wide diversity of folding patterns, characterized by discrete interaction *islands* and *deserts* (**Supplementary Fig. 4**). This organisation is reminiscent of the topological domains and A/B compartments that define eukaryotic chromosome architecture^26^, suggesting that bacterial RNAs also adhere to principles of higher-order structural organisation.

Beyond recapitulating the overall fold, the ligation frequency data can be used to identify specific structural features. By profiling the per-nucleotide frequency of long-range intermolecular ligation events, which serves as a proxy for local accessibility, it is possible to probe accessible (unpaired and stem termini) regions of highly structured RNAs like the sRNA CsrB which harbours dozens of stem loops (**Fig. 1f**) and hkRNAs like ribosomal RNAs, RNase P RNA and 6S RNA (**Supplementary Figs. 5, 6, 7**). As intermolecular ligation events requires base exposure and close spatial proximity, the TRIC-seq reports accessibility in the native three-dimensional context, while DMS-seq primarily reports base-pairing state^27^.

### *trans*-interactome reveals sRNA regulatory hubs

With structural mapping validated, TRIC-seq was next applied to uncover genome-wide trans-acting interactions. GcvB, one of the best characterised bacterial sRNAs^28–31^, was first used first to illustrate TRIC-seq’s readout. GcvB associated chimeras were visualised as arc lines in IGV (**Fig. 2a**): arcs radiating from GcvB locus aligned with several of its known targets. Guided by this locus-level view, transcriptome-wide trans-acting interactions were called using a configuration-model framework that preserves each RNA’s interaction frequency (degree) and generates a null from one million randomized interactomes. For each RNA pair, an enrichment odds ratio (*O^f^*) and an FDR-corrected P value were computed against this null, separating specific contacts from stochastic proximity **(Supplementary Fig. 8a)**. Measurements were highly reproducible across biological replicates: interaction counts between RNA pairs showed strong concordance **(Supplementary Fig. 8b)**, and odds ratios for shared pairs correlated robustly between replicates **(Supplementary Fig. 8c)**. Consistent with sRNA-mediated regulatory control in bacteria, sRNA-5′UTR pairs were among the most enriched pairings **(Supplementary Fig. 8d,e)**.

**Figure 2.**
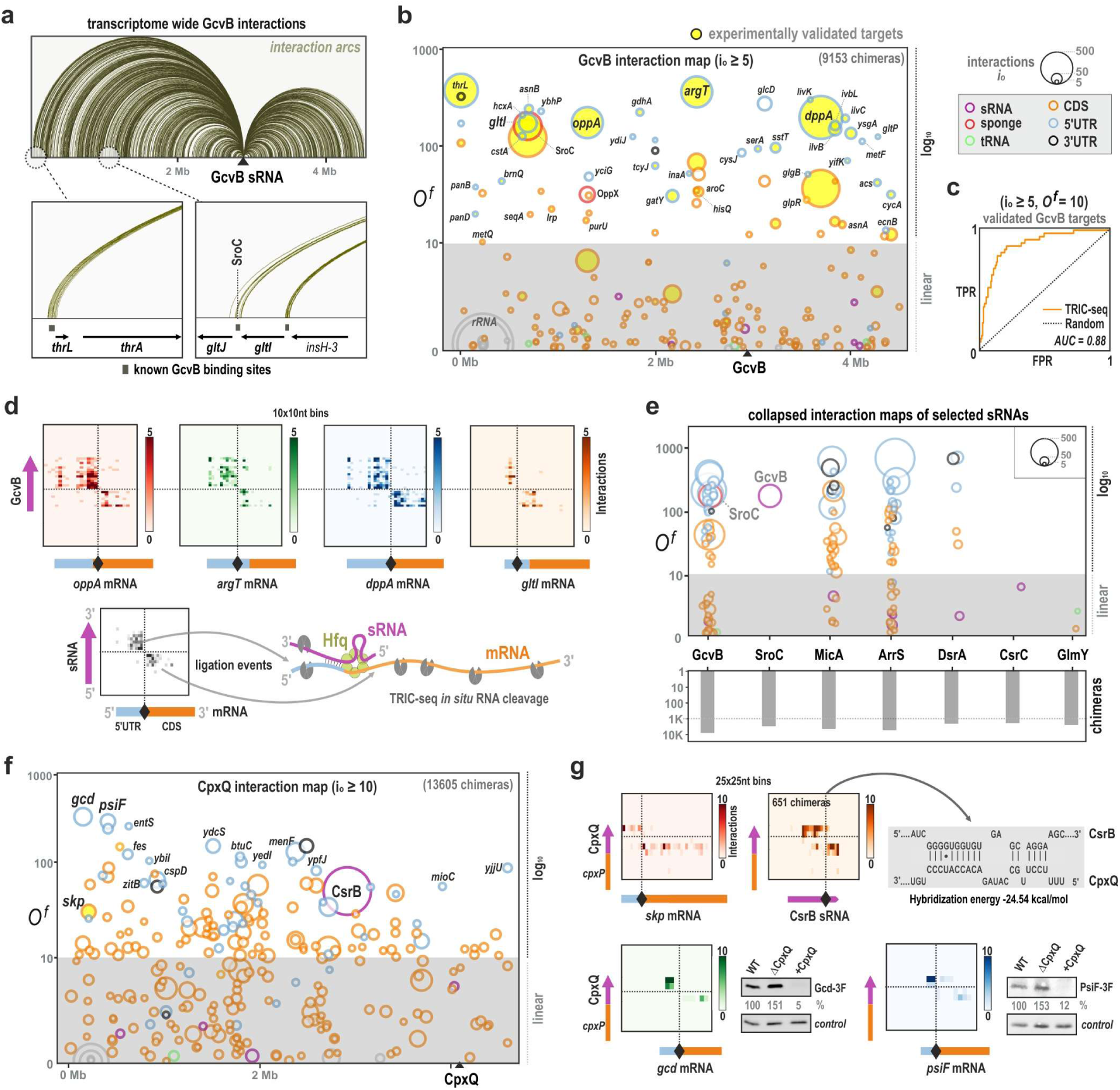
TRIC-seq delineates sRNA regulons and discovers novel interactions. **a**, Transcriptome-wide arc diagram of GcvB interactions from one *E. coli* replicate. Each arc denotes a unique long-range interaction from the GcvB locus. Insets, zooms over validated targets (*thrL*, *gltI*) and the sponge SroC; black bars indicate known GcvB binding sites. **b,** Global interaction map of the GcvB (9153 chimeras) showing partners with interaction count i_o_ ≥ 5. x-axis, genomic coordinate; y-axis, odds ratio (*O^f^*) against the degree-preserving configuration-model null. Circle area ∝ supporting chimeric reads (i_o_); edge colour encodes RNA feature class (sRNA, 5′UTR, CDS, 3′UTR, tRNA, sponge; legend in panel). Literature-validated targets are highlighted in yellow^30^. **c,** ROC curve distinguishing the 56 validated GcvB targets from other transcripts using *O^f^*as the score (AUC = 0.88; dashed line, random). **d,** Inter-RNA heat maps (10 × 10-nt bins) localise GcvB binding on four targets—*oppA*, *argT*, *dppA*, *gltI*. Focused diagonal signals mark the base-pairing interface (schematic below). **e,** Collapsed interaction maps (csMaps) for representative sRNAs. Hub sRNAs (e.g., GcvB, MicA, ArrS) show numerous high- *O^f^*partners, whereas protein-sequestering sRNAs (CsrC, GlmY) lack significant RNA targets. Circle size ∝ i_o_; scale inset at top right. **f,** Global interaction map for CpxQ (io ≥ 10; 13605 chimeras) reveals an extensive regulon and a specific interaction with the major regulatory RNA CsrB. **g,** Top left, inter-RNA heat map confirming the known CpxQ–*skp* interaction. Top right, CpxQ–CsrB heat map with predicted duplex at the binding site using IntaRNA^74^. Bottom, immunoblots: Δ*cpxQ* increases Gcd and PsiF protein levels (3xFLAG tagged), whereas CpxQ overexpression decreases them, consistent with direct CpxQ-mediated repression.

Global interaction map for GcvB summarized targets by genomic coordinate (x-axis) and *O^f^* (y-axis), with point size proportional to supporting chimeric events (io) and colour denoting RNA features (**Fig. 2b**). Fragmentation during TRIC-seq library construction provided sufficient resolution to assign contacts to specific transcript regions (5’UTR, CDS, or 3’UTR). Across two replicates, TRIC-seq produced highly overlapping sets of GcvB targets **(Supplementary Fig. 9a)**.

For benchmarking trans regulatory interactions, the 56 literature-validated GcvB targets were treated as positives. Using *O^f^* as the ranking statistic and restricting to GcvB interacting targets with io ≥ 5, ROC analysis yielded AUC = 0.88 and an optimal threshold of *O^f^* = 6.95 (**Fig. 2c**). For reporting, a conservative criterion of *O^f^* ≥ 10 together with io ≥ 5 was adopted; **Supplementary Fig. 9b shows** how specificity and precision remain high across io thresholds under this *O^f^* cut-off. Under these settings, 54 GcvB partners were identified and 31/56 validated targets were recovered. The resulting GcvB interactome shows strong concordance with Hfq pull-down based RIL-seq^23^ and CLASH^16^ while substantially expanding the known regulon (**Supplementary Fig. 10**).

Individual RNA-RNA interactions were visualized using inter-RNA heat maps, which plots the interactions along the sequence of one RNA against another. For a canonical sRNA-mRNA interaction, this results in a focused signal at the diagonal around the base-pairing region, as seen for interaction of GcvB with *oppA*, *argT*, *dppA* and *gltI* mRNAs at 10 bp resolution (**Fig. 2d**). Intra–inter composite maps further contextualized trans interactions together with their individual RNA structures (**Supplementary Fig. 11**). This is exemplified by the interaction between RyfD sRNA and *lpp* mRNA encoding the lipoprotein Lpp (**Supplementary Fig. 12**). While TRIC-seq revealed RyfD binding to the 5’UTR of *lpp*, the intra-inter map reveals ligation events between RyfD and the *lpp* 3’UTR owing to the mRNA’s tertiary structure, which brings the two ends of the mRNA into close spatial proximity.

To summarize the complete set of partners for any given RNA, a collapsed interaction map (csMap) was used, which flattens genomic positions onto a single axis and enables side-by-side comparison of target profiles across multiple RNAs. csMaps contrasted multi-target sRNAs (GcvB, MicA, ArrS) with protein-sequestering RNAs (CsrC^32^, GlmY^33^) that lacked RNA targets despite ample chimeric reads, underscoring methodological specificity rather than coverage limitations (**Fig. 2e; Supplementary Fig. 13)**. Like GcvB, other multitarget sRNAs (e.g. MgrR, DsrA, RprA, RyhB, CpxQ) also showed strong overlap with RIL-seq-defined targets using the same thresholds^23^ **(Supplementary Fig. 13,14)**.

The platform’s power for discovery was highlighted by the analysis of CpxQ sRNA, derived from the 3’UTR of *cpxP* mRNA^34^. The global interaction map for CpxQ revealed several targets related to inner membrane and periplasmic stress as previously observed in the related bacterium *Salmonella enterica*^34^ (**Fig. 2f, Supplementary Fig. 14b**). TRIC-seq uncovered CpxQ’s sole known target in *E. coli*, encoding the periplasmic chaperone, *skp*^35^. Remarkably, CpxQ was found to specifically interact with CsrB^36^, a major regulatory RNA (**Fig. 2f, g, Supplementary Fig. 15**). This finding is of particular interest because CsrB, known exclusively as an antagonist of the post-transcriptional regulatory protein CsrA, is now found to also bind CpxQ sRNA, potentially coordinating the post-transcriptional regulation of both of their large regulons^23,34,37^. Next, two of the newly identified targets, *gcd* and *psiF*, were validated at the protein level. CpxQ deletion led to the upregulation of both Gcd and PsiF proteins, while CpxQ overexpression led to their downregulation, consistent with CpxQ binding near the Shine-Dalgarno (SD) sequence in the inter-RNA heat maps (**Fig. 2g**).

### The RNA interactome is organized by RNA class and function

An analysis of transcriptome-wide long-range interactions revealed that different classes of RNA display unique interaction profiles. csMaps of 5’UTRs show that they are overwhelmingly dominated by contacts with sRNAs, confirming their role as primary regulatory hubs (**Fig. 3a**). CDSs also interact with sRNAs, typically near the SD region; unlike 5’UTRs, CDSs additionally contact tRNAs and hkRNAs, consistent with ongoing translation (**Fig. 3b, Supplementary Fig. 16**).

**Figure 3.**
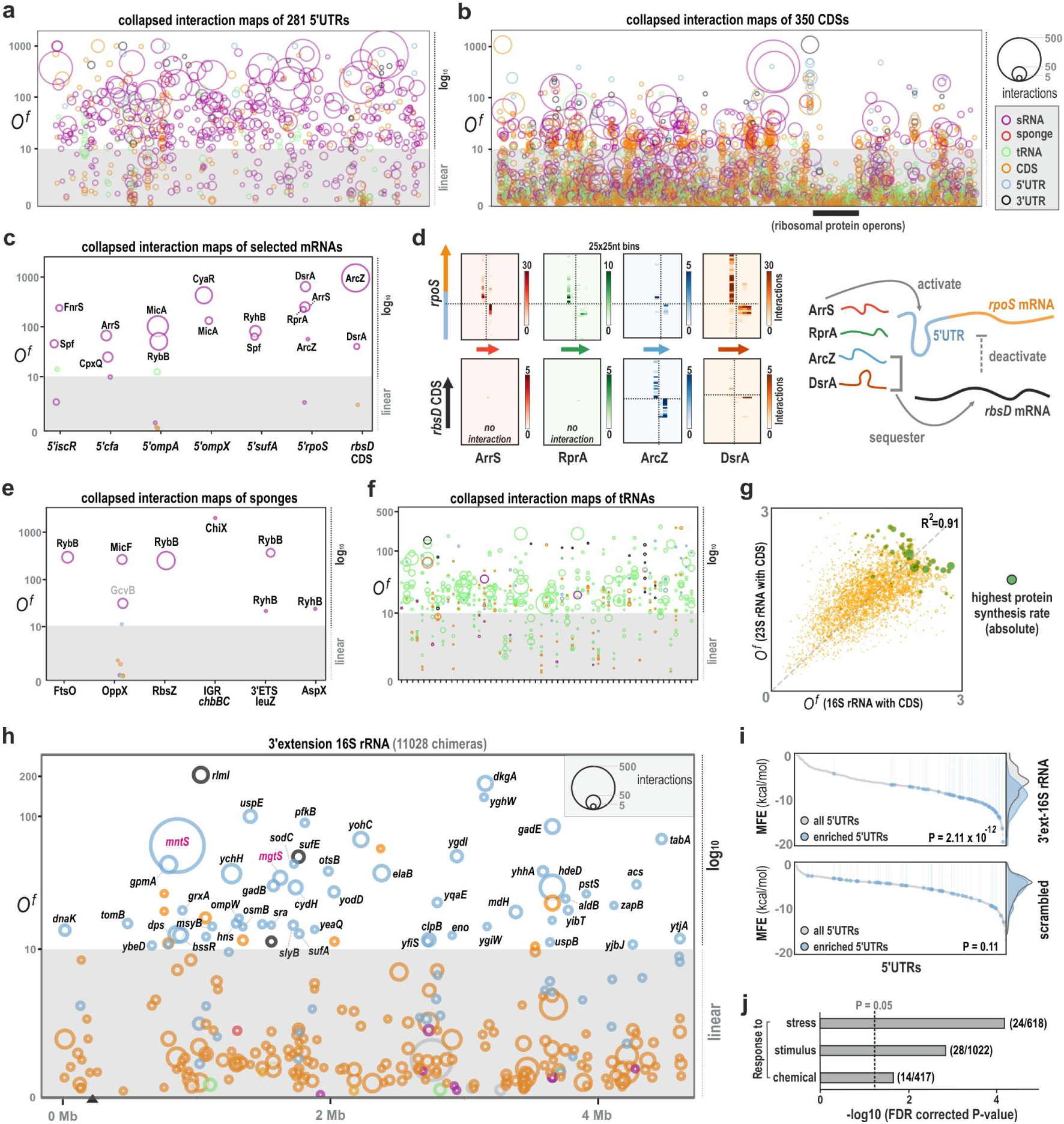
Global interactome organisation reveals RNA-class fingerprints and a 16S-tail tethering mechanism. **a**, Collapsed interaction maps (csMaps) for the 281 most interactive 5’UTRs (genome order). Partners are dominated by sRNAs (magenta); circle area ∝ supporting chimeric reads (i_o_); edge/fill colours denote RNA class (legend inset). **b,** csMaps for the 350 most interactive CDSs (genome order). Black bar marks the genomic region encoding ribosomal protein operons. **c,** csMaps for selected regulatory mRNAs illustrate diverse patterns: sRNA-targeted 5’UTRs of *iscR, cfa, ompA, ompX, sufA, rpoS* and the *rbsD* CDS. **d,** Inter-RNA heat maps (25 × 25-nt bins) resolve the post-transcriptional logic at *rpoS*: activating sRNAs (ArrS, RprA, ArcZ, DsrA) bind 5’UTR of *rpoS* (top row). The *rbsD* CDS sequesters ArcZ/DsrA (bottom row); schematic at right. **e,** csMaps of known RNA sponges show highly cognate, specific interactions with their sRNA targets (e.g., RybB sponges FtsO, 3′ETS-*leuZ*, RbsZ; low background to other RNAs). **f,** csMaps of tRNAs reveal primary contacts with other tRNAs and CDSs, consistent with co-localisation at translating ribosomes. **g,** Odds ratios for CDS contacts with 23S versus 16S rRNAs are tightly correlated (R² = 0.91). CDSs with the highest absolute protein-synthesis rates (green) show the strongest association with both subunits. **h,** Global interaction map for the 3′ end of 16S rRNA (11,028 chimeras) shows widespread, specific contacts with 5’UTRs. **i,** Cumulative distributions of minimum free energy (MFE) for predicted duplexes between 5’UTRs and the 16S 3′ extension. Observed (enriched) 5’UTRs form significantly more stable duplexes than all 5’UTRs (P = 2.11 × 10⁻¹², top); the effect is absent with a scrambled 3′-tail control (P = 0.11, bottom). **j,** Gene Ontology enrichment for mRNAs whose 5’UTRs interact with the 16S 3′ extension. Stress/stimulus-response terms are most enriched (bars show −log₁₀ FDR; counts at right).

Examination of individual loci illustrates this functional diversity (Fig. 3c). The *iscR* 5′UTR is targeted specifically by FnrS and Spf, concordant with overexpression library screens identifying these sRNAs as repressors of IscR translation^38^. Regulatory logic is also resolved at *rpoS*: multiple sRNAs bound distinct activating sites on the *rpoS* 5′UTR, including the new binding partner ArrS sRNA, which occupied the same site as the activating sRNAs RprA, ArcZ, and DsrA^39,40^. Furthermore, TRIC-seq supports a recent report that the overexpression of the *rbsD* CDS sequesters the sRNAs DsrA and ArcZ, leading to reduced RpoS^41^ (**Fig. 3d, Supplementary Fig. 16**).

Unlike sRNAs which often target multiple functionally related RNAs, sponge RNAs like SroC typically bind and sequester a specific sRNA^42,43^ (**Fig. 2e**). The TRIC-seq captured several of these highly cognate interactions with high specificity, consistent with prior observations for SroC^43^ (**Fig. 3e**). TRIC-seq identified the three sponges of RybB sRNA – FtsO^44^ (*ftsI* ORF derived), 3’ETS-leuZ^45^ (tRNA extension) and RbsZ^46^ (*rbsB* 3’UTR derived), along with previously known (*ompA, ompC, ompW, fadL*)^47^ and new (*bamA, bhsA, yhjY, loiP*) outer membrane protein encoding mRNA targets (**Fig. 3e, Supplementary Fig. 17**). TRIC-seq also suggest new potential sponges based on highly specific pairing to individual sRNAs, such as the alanine tRNA fragment which was found to interact with SdsR sRNA (**Supplementary Fig. 17e**).

Apart from regulatory RNAs, TRIC-seq could also resolve the interactome of hkRNAs. As a class, tRNAs interacted predominantly with other tRNAs and CDSs, reflecting shared ribosomal occupancy during active translation (**Fig. 3f**). CDS–rRNA coupling provided an internal control. Across CDSs, *O^f^*values for 16S and 23S were not only highly correlated but nearly identical (*O^f^* with 16S ≈ *O^f^* with 23S), indicating that TRIC-seq predominantly captures contacts with assembled 70S ribosomes during translation (**Fig. 3g; Supplementary Fig. 18)**. Moreover, CDSs with the highest absolute protein-synthesis rates^25^ showed the highest odds ratios with both subunits, quantitatively linking the interactome to translation output. Other hkRNAs, including RnpB and tmRNA, displayed distinct profiles based on their role in tRNA maturation and trans-translation, respectively (**Supplementary Fig. 19)**.

Global profiles also revealed enriched contacts between the 3′ extension of pre-16S rRNA and the 5′UTRs of 49 mRNAs (**Fig. 3h**). During 16S maturation this tail persists only briefly before RNase trimming, during which it can base-pair with mRNA leaders. The enriched 5′UTRs form markedly more stable duplexes with the tail than 5′UTRs genome-wide (P = 2.11 × 10⁻¹²), whereas this difference disappears with a scrambled-tail control (**Fig. 3i**). These observations are consistent with a tethering model in which the pre-16S 3′ extension base-pairs with selected 5′UTRs, positioning them near the small (30S) subunit and potentially facilitating early steps of translation initiation. The associated mRNAs were found to be enriched for stress/stimulus-response Gene Ontology terms (**Fig. 3j**); examples include *mgtS* and *mntS*, which encode small inner-membrane proteins regulating Mg²⁺ and Mn²⁺ homeostasis in *E. coli*^48,49^ (**Supplementary Fig. 20a**).

### Modular architecture of the *E. coli* RNA interactome

After profiling RNA-class fingerprints, the global organisation of the interactome was evaluated using unsupervised clustering of *E. coli* RNA features (5′UTRs, CDSs, 3′UTRs and sRNAs) that had at least one trans interaction with another feature supported by at least 5 chimeric reads. The network was strikingly modular, segregating into 30 clusters (**Fig. 4a**). Features were grouped by similarity of target/partner profiles rather than by direct edges, and major sRNA regulons emerged as discrete modules; for example, targets of GcvB, CyaR, Spf, and RybB resolved into separate clusters (**Fig. 4b**). A systematic enrichment analysis confirmed that targets of nearly every multitarget sRNA concentrated within one or two specific clusters, mapping regulatory inputs directly onto the module structure of the interactome **(Supplementary Fig. 21)**.

**Figure 4.**
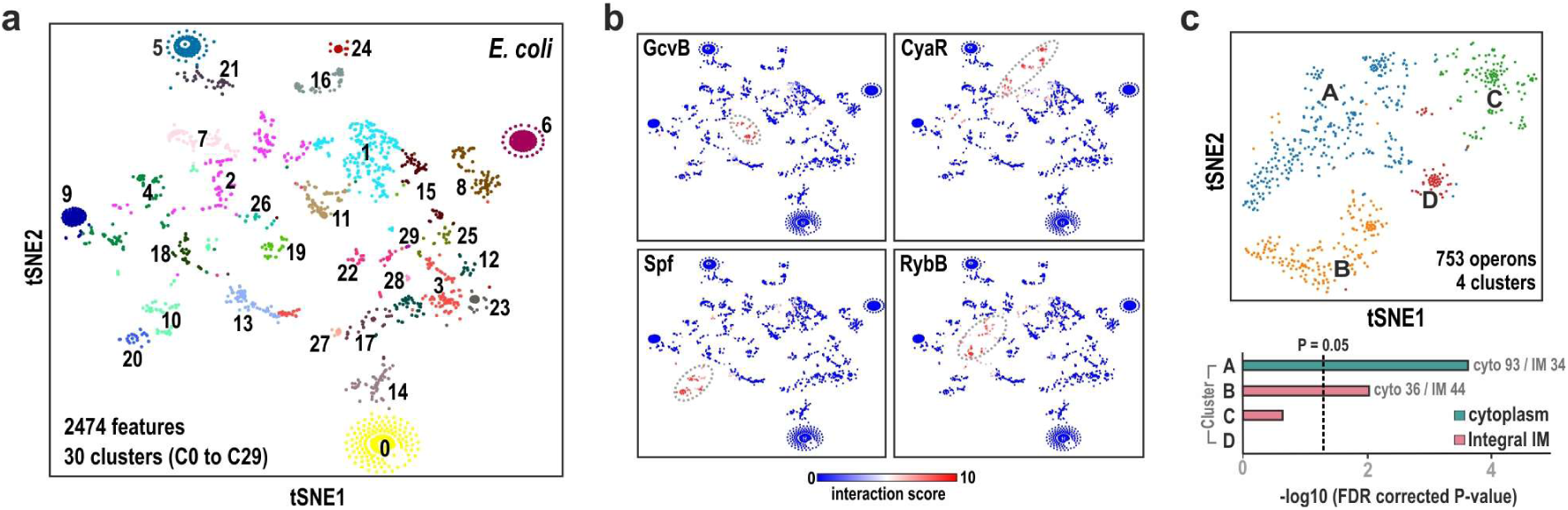
Unsupervised clustering reveals modular RNA interactome structure. **a**, Two-dimensional t-SNE of 2,474 *E. coli* RNA features clustered by shared long-range interaction partners. The interactome segregates into 30 distinct clusters. Each dot is a feature (5′UTR, CDS, 3′UTR and sRNA), coloured by cluster. **b,** The same embedding as in (a) overlaid with normalised interaction scores for four sRNAs (GcvB, CyaR, Spf, RybB). Targets concentrate within specific clusters (dashed ellipses). **c,** Top, t-SNE embedding of 753 protein-coding operons (sRNAs excluded) clustered by their interaction profiles, resolving four communities. Bottom, enrichment analysis (bars, −log₁₀ FDR; dashed line, FDR = 0.05). Cluster A is enriched for operons encoding cytoplasmic proteins only, whereas Cluster B is enriched for operons encoding at least one integral inner-membrane (IM) protein.

Transcripts encoding cytoplasmic and inner membrane proteins are known to occupy distinct subcellular zones in bacteria^7,8^. Thereby, even low odds, proximity-driven interactions may retain special information. When transcripts from entire protein-coding operons were clustered by their inter-transcript interaction profiles, the network segregated into communities enriched for shared cellular destinations (cytoplasm or inner membrane) (**Fig. 4c, Supplementary Fig. 22)**. Together, these patterns indicate that the modular organisation resolved by TRIC-seq reflects both regulatory inputs and underlying spatial organisation within the cell.

### A CDS-centric hub forms a stress-related RNA condensate

A notable exception to sRNA-organized modules was Cluster 13 (C13) (**Fig. 4a**). When the embedding was recomputed using only CDS-CDS interactions, CDSs in sRNA-dependent clusters (e.g., C11, maintained by interactions with the sRNA SdsR) dispersed, whereas C13 remained tightly cohesive (**Fig. 5a; Supplementary Fig. 23)**. csMaps for C13 CDS members showed predominant contacts with other CDSs, overwhelmingly within the same cluster, yielding a dense intra-cluster web in circular interaction views (**Fig. 5b,c; Supplementary Fig. 24)**. These features indicate that direct mRNA–mRNA associations, rather than shared sRNA partners, underlie C13 cohesion. Furthermore, Gene Ontology analysis for C13 mRNAs revealed strong enrichment for stress and temperature-response functions (Fig. 5d).

**Figure 5.**
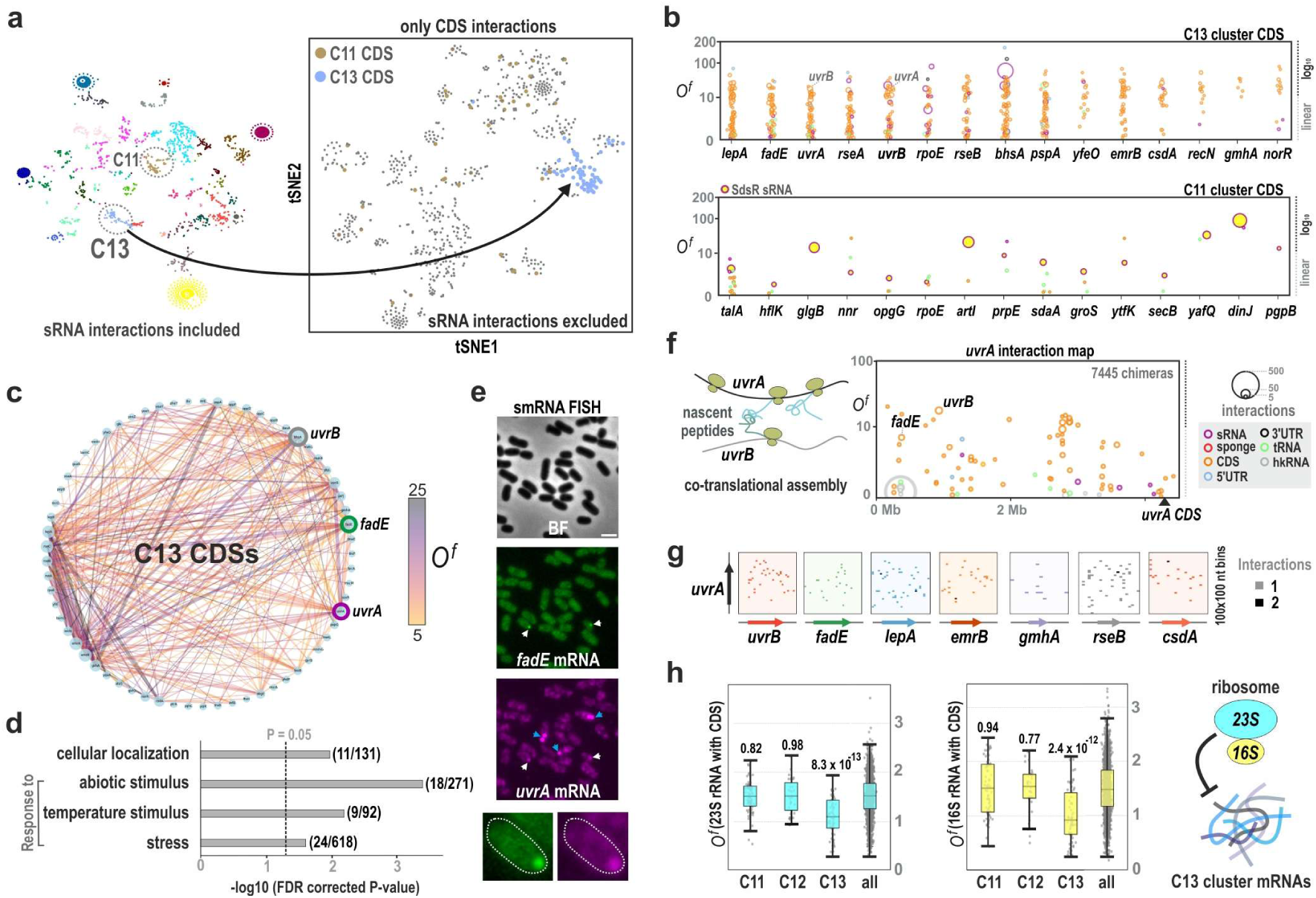
A CDS-centric interactome module forms a ribosome-poor, granule-like mRNA condensate. **a**, t-SNE embeddings highlight members of C11 and C13 clusters. Left, embedding from Fig. 4a includes sRNA interactions. Right, the embedding recomputed using only CDS features: the CDS in sRNA-organized cluster C11 disperses, whereas CDS members in C13 remains cohesive, indicating maintenance by direct mRNA–mRNA contacts. **b,** Collapsed interaction maps (csMaps) for representative CDSs from C13 (top) versus C11 (bottom). C13 CDSs predominantly contact other CDSs (orange), whereas C11 members are organized by shared interactions with the sRNA SdsR. y-axis, odds ratio (*O^f^*); circle area ∝ supporting chimeras (i_o_); colours denote RNA class. **c,** Circular network of C13 CDSs showing dense intra-cluster connectivity. Edge width ∝ supporting chimeric reads; edge colour encodes *O^f^*. Key nodes (e.g., *uvrA*, *uvrB*, *fadE*) are highlighted. **d,** Gene Ontology enrichment for C13 genes. Bars show − log₁₀(FDR); dashed line, FDR = 0.05. Stress- and temperature-response terms are over-represented. **e,** smFISH demonstrates in vivo co-aggregation: *fadE* mRNA (green) and *uvrA* mRNA (magenta) co-localise in cytoplasmic foci (white arrowheads). Self-association of *uvrA* into larger aggregates is indicated by blue arrows. **f,** Global interaction map for *uvrA* (7445 chimeras) shows interaction with C13 members; strong *uvrA–uvrB* contacts are consistent with co-translational complex assembly (schematic, left) that could nucleate the condensate. **g,** Inter-RNA heat maps (100 × 100-nt bins) for *uvrA* paired with C13 members (*uvrB, fadE, lepA, emrB, gmhA, rseB, csdA*). Signals are distributed across coding regions rather than focused at a defined site, consistent with co-aggregation rather than discrete base-pairing. **h,** Boxplots of *O^f^*for CDS contacts with rRNAs. Left, 23S; middle, 16S; right, schematic. C13 shows the lowest ribosome association compared to other clusters (C11 and C12 shown) and all CDSs (two-sided Wilcoxon tests), consistent with ribosome exclusion in the condensate.

In vivo imaging supported a physical basis for this organisation. smFISH detected *uvrA* transcripts in most cells, with multi-mRNA foci in a subset of cells; dual colour smFISH for *uvrA* and *fadE* mRNAs (both C13 members) showed cytoplasmic co-localisation consistent with co-aggregation (**Fig. 5e; Supplementary Fig. 25)**. TRIC-seq contact patterns suggested potential nucleation mechanisms: despite being ∼1 Mb apart on the chromosome, *uvrA–uvrB* exhibited strong interactions (**Fig. 5f**), consistent with co-translational assembly of the UvrAB complex. Inter-RNA heat maps of *uvrA* with seven other C13 members showed diffuse, transcript-wide signals rather than focused interfaces (**Fig. 5g**), consistent with general co-association rather than site-specific duplexes typical of sRNA–mRNA regulatory interaction.

A defining property of this hub was ribosome exclusion. C13 CDSs showed the lowest odds ratios with 16S and 23S rRNAs among clusters (**Fig. 5h**). A case study of *emrA* (C13) revealed an interactome profile dominated by other C13 members and depleted for rRNA contacts, in contrast to the control *ompA*, which was enriched for rRNA interaction and showed minimal CDS–CDS association **(Supplementary Fig. 26)**. Collectively, these observations support a higher-order regulatory principle in which large cohorts of stress-related transcripts are sequestered into protective, translationally poor, granule-like assemblies.

### RNA Interactomes across diverse bacterial species

Most understanding of bacterial riboregulation stems from a handful of model species, leaving much of the microbial RNA interactome unexplored^50^. To assess generality and enable de novo regulon discovery, TRIC-seq was applied to three phylogenetically diverse bacteria.

Firstly, TRIC-seq was performed on *Stutzerimonas stutzeri,* a Gram-negative bacterium harbouring Hfq^51^. Unsupervised clustering of long-range interactions resolved a modular interactome with 28 clusters (**Fig. 6a**). Clustering of partner profiles uncovered candidate sRNAs, including SSnc1, which derives from the 3′ end of PSJM300_04280, encoding a cAMP-dependent protein kinase. The global interaction map for SSnc1 revealed several putative mRNA targets (**Fig. 6b**). Inter-RNA heat maps for 18 of these targets confirmed site-specific binding near their SD regions (**Fig. 6c**). Notably, the *S. stutzeri* interactome also contained a cluster highly enriched for contacts with the 16S rRNA 3′ extension, consistent with the tethering mechanism observed in *E. coli* (**Fig. 6a, Supplementary Fig. 27**).

**Figure 6.**
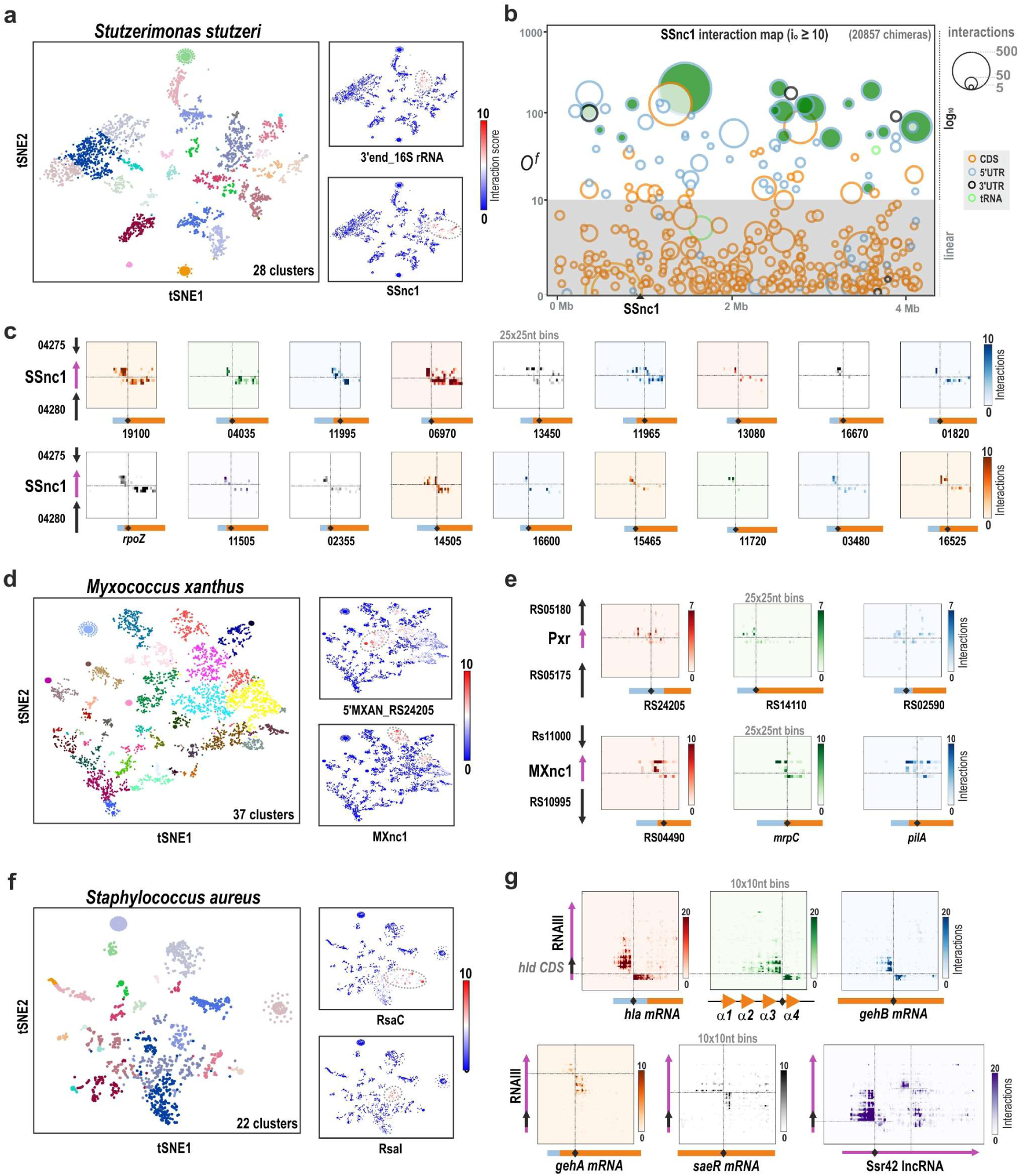
Modular RNA interactome of diverse microbes. **a**, t-SNE embedding of the *Stutzerimonas stutzeri* interactome. Insets show overlays with normalised interaction scores for the 3′ extension of 16S rRNA and the sRNA SSnc1. **b,** Global interaction map for SSnc1 (i_o_ ≥ 10; 20857 chimeras). x-axis, genomic coordinate; y-axis, odds ratio (*O^f^*) versus a configuration-model null. Circle area ∝ chimeric reads; colours denote RNA class. 5′UTR targets highlighted in green are shown in (e). **c,** Inter-RNA heat maps (25 × 25-nt bins) for 18 SSnc1 targets, revealing site-specific binding near the SD region of the 5′UTRs. **d,** t-SNE of the *Myxococcus xanthus* interactome. Insets: overlays with normalised interaction scores for candidate sRNAs originating near MXAN_RS24205 and MXAN_RS10995. **e,** Inter-RNA heat maps (25 × 25-nt bins) for the developmental sRNA Pxr and for a candidate sRNA MXnc1, showing specific binding near the SD sequence of multiple mRNA targets. Pxr is a known sRNA controlling fruiting-body development in *M. xanthus*, but its trans-targets have remained undefined. **f,** t-SNE of the *Staphylococcus aureus* interactome. Insets: overlays with normalised interaction scores for RsaC and RsaI sRNAs. **g,** Inter-RNA heat maps (10 × 10-nt bins) showing RNAIII interactions with *hla* and *psmα4* along with new targets (*gehB*, *gehA*, *saeR*) and the long non-coding RNA Ssr42.

The broad applicability of TRIC-seq is further underscored by its performance in organisms where the canonical RNA chaperone Hfq is not the central player in sRNA-mediated regulation. In *Myxococcus xanthus*, which lacks Hfq^52^, the interactome partitioned into 37 clusters (**Fig. 6d, Supplementary Fig. 28a)**. TRIC-seq delineated trans-targets for the developmental sRNA Pxr^53^, whose trans targets had remained elusive. Furthermore, TRIC-seq identified the candidate sRNA MXnc1 originating from an intergenic region. Inter-RNA heat maps show site-specific pairing of MXnc1 with multiple mRNA targets including *mrpC* and *pilA* (**Fig. 6e).**

In the Gram-positive pathogen *Staphylococcus aureus*, Hfq is not essential for regulatory sRNA-mRNA interactions^54,55^, RNA interactome clustering again revealed a clear modular structure (**Fig. 6f, g, Supplementary Fig. 28b)**. The TRIC-seq data from *S. aureus* allowed for a detailed analysis of its extensively studied long regulatory RNA, RNAIII (514 bp), a major virulence regulator^56^. Canonical RNAIII targets including *hla* (encoding α-hemolysin) and *psmα4* were mapped with precision alongside established regulators (*rot, RpiRc*) and several new targets like *saeR, gehA,* and *gehB*. The identification of *saeR* is noteworthy, as the SaeRS two-component system itself controls the expression of multiple virulence factors including α-hemolysin^58^. *gehA* and *gehB* mRNAs encode secreted lipases known to hydrolyze triglycerides to help the bacterium evade host innate immunity^59^, fit well with RNAIII’s global role in virulence. Interactions with *hla*, *psmα4*, and *gehB* mRNAs converged at the same RNAIII site immediately upstream of the *hld* ORF^56^, whereas the binding to *gehA* and *saeR* mRNAs occurs further downstream (**Fig. 6g**). Furthermore, TRIC-seq captured several sRNA-sRNA interactions, such as RsaI-RsaE and RsaI-RsaG, which were also identified using pull-down based MAPS and CLASH approaches^57,60^ **(Supplementary Fig. 28c).** Strikingly, a prominent interaction was discovered between RNAIII and the long non-coding RNA Ssr42 (1233 nt)^61,62^ (**Fig. 6g**). This interaction is particularly intriguing, as several studies have shown that deletion of *ssr42* leads to reduced expression of RNAIII targets like α-hemolysin and PSMα4^62–64^. The physical interaction captured by TRIC-seq suggests that Ssr42 regulates virulence by directly binding to and modulating the activity of RNAIII.

## Discussion

This work introduces TRIC-seq, a technology that provides an unbiased, in situ map of the bacterial RNA interactome. By capturing both intramolecular tertiary structures and intermolecular interactions simultaneously, it offers an unprecedented view into the organisational principles of the bacterial transcriptome. The findings confirm and extend numerous previous studies on regulatory interactions while revealing novel, system-level properties of RNA networks that pave the way for future investigations.

A new era of bacterial transcriptomics is dawning, driven by technologies that probe gene expression at the single-cell and subcellular level. Single-cell RNA sequencing (scRNA-seq) methods have revealed transcriptional diversity within isogenic bacteria populations^22,65^, answering *what* genes are expressed in individual cells. Complementary imaging-based technologies like bacterial-MERFISH now provide an answer to *where* these transcripts are located^7^. However, a critical dimension has remained missing: a global map of *which* RNAs are physically interacting. Such a map, provided here by TRIC-seq, is essential for connecting gene expression and spatial organisation to the functional regulatory networks that govern cellular behaviour.

Together with recent reports of stress-mediated mRNA aggregation in bacteria^66,67^, this study suggests that the co-association of mRNAs into protective, translationally poor condensates is a conserved strategy for managing the transcriptome during adverse conditions^68,69^. Future work will focus on the formation, regulation, and function of such condensates in bacteria. Another significant finding is the widespread interaction between the 16S rRNA 3’ extension and the 5’UTRs of stress-related mRNAs, suggesting a tethering mechanism that primes the translation of a defined set of mRNAs under specific stress conditions. This observation adds to the growing body of evidence for ribosome heterogeneity and specialized translation in bacteria^70^, where subpopulations of ribosomes translate specific mRNAs in response to environmental cues.

The datasets generated here represent a foundational resource. Transcriptome-wide contact maps provide orthogonal restraints for training and benchmarking AI-driven RNA folding models, addressing a bottleneck in deriving accurate tertiary structures from sequence alone^71^. Perturbational comparisons, such as wild type versus RBP deletions, can separate RNA structures and trans contacts under RBP control and help distinguish structural modulators from interaction chaperones ^72,73^.

The permeabilized-cell enzymatic workflow is compatible with split-pool based bacterial scRNA-seq methods^22^, enabling future integration to map interactomes within transcriptionally defined subpopulations. The genetics-free nature further permits application to non-model microbes and mixed communities, opening the possibility to assess RNA exchange and inter-species regulatory RNA-RNA interactions. In conclusion, TRIC-seq provides a broadly applicable tool to chart the bacterial RNA interactome with high specificity in native context, offering a platform to systematically explore regulatory RNA networks across the microbial world.

## Supporting information

Source data for main figures

---

**Supplementary Figure 1.**
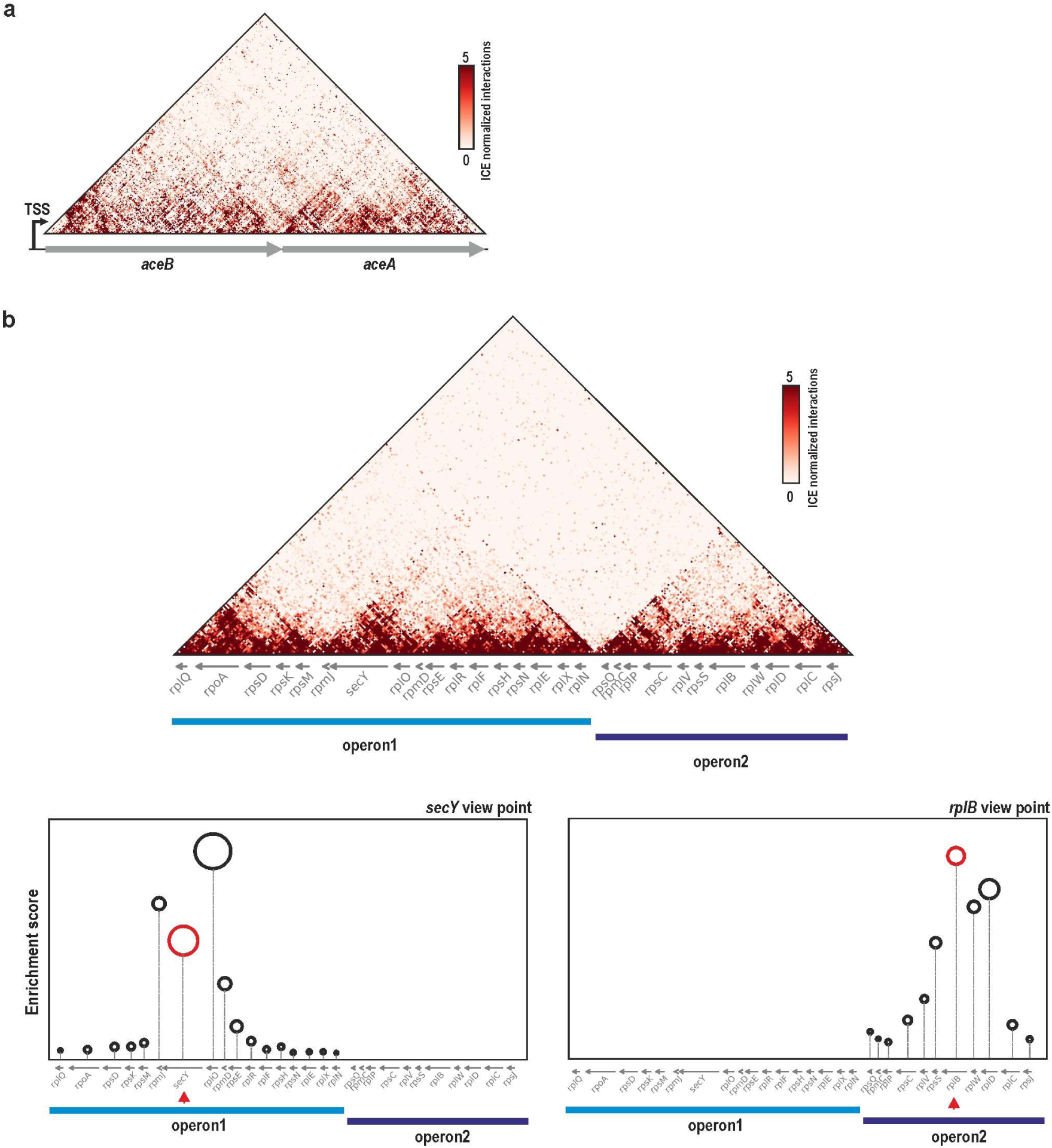
ORF-centric folding within operons and confinement of interactions across operon boundaries. **a**, ICE-normalized contact map of the polycistronic *aceB–aceA* mRNA. Each CDS (*aceB*, *aceA*) forms a semi-independent structural domain (TSS indicated), illustrating the ORF-centric folding principle highlighted in Fig. 1b. **b,** Top, ICE-normalized contact map across two adjacent ribosomal-protein operons (operon1 and operon2); a sharp drop in contacts at the operon junction indicates that interactions are largely confined within operon boundaries. Bottom, enrichment plots for *secY* (operon1) and *rplB* (operon2). Each circle represents the pair between the query gene and a gene in the locus; the y-axis shows the enrichment score (Methods), and circle area is proportional to supporting interactions. Blue and purple bars denote operons. The red circle marks the query gene’s self-interaction; the red triangle marks its genomic position on the x-axis.

**Supplementary Figure 2.**
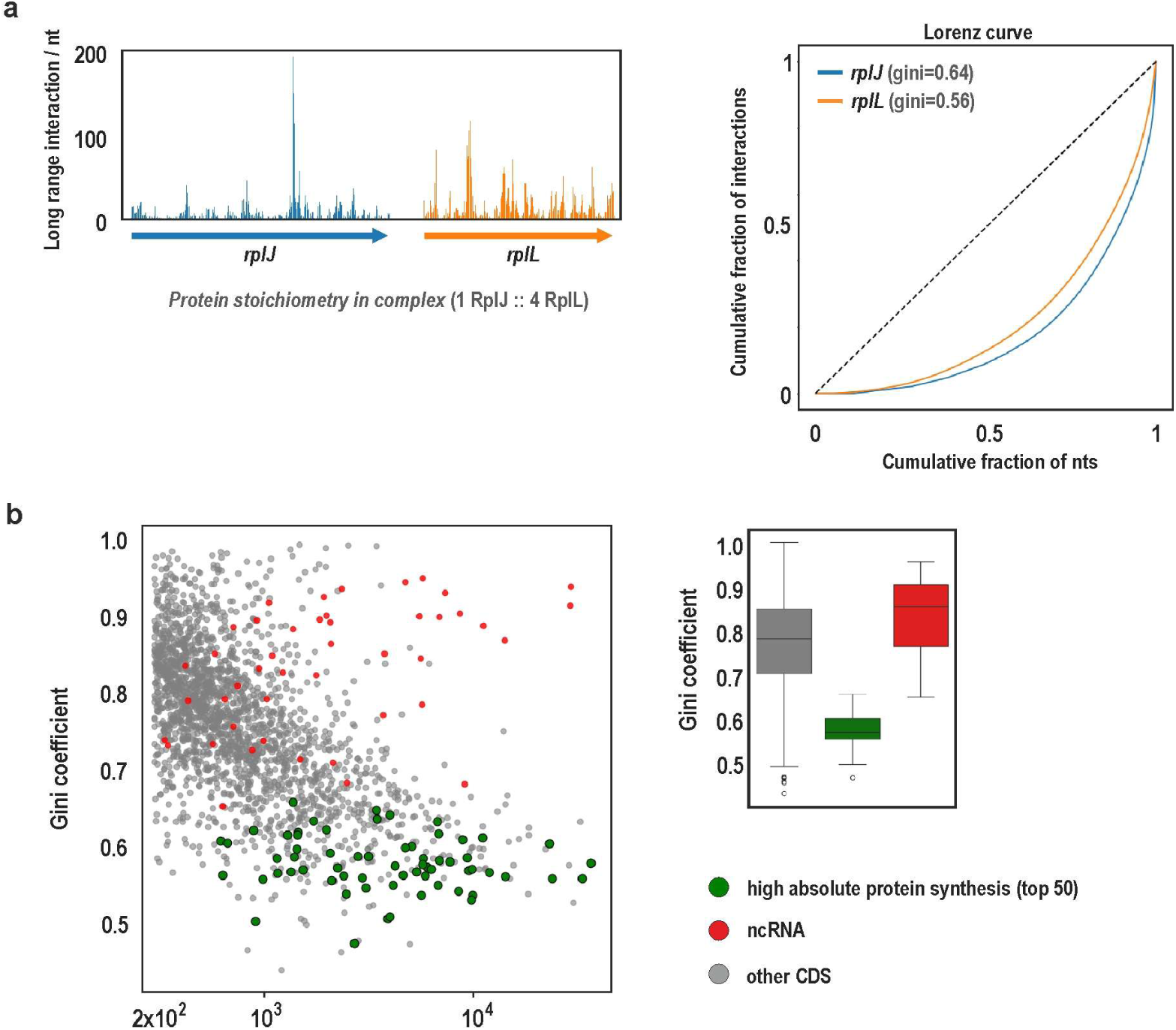
The Gini coefficient of long-range interactions reports RNA structural heterogeneity and translational state. **a**, Ribosomal protein operon, *rplJ*-*rplL*. Left, per-nucleotide profiles of long-range ligations (interactions/nt) show sharper, localised peaks for *rplJ* than for *rplL*. Right, Lorenz curves (cumulative fraction of interactions vs cumulative fraction of nucleotides) and corresponding Gini coefficients quantify this difference: *rplJ* (Gini = 0.64) deviates further from the line of equality than *rplL* (Gini = 0.56), indicating a more uneven, structured contact distribution. **b,** Genome-wide relationship between Gini and RNA class/function. Left, scatter of Gini versus total long-range interactions for all CDSs; highly expressed protein-coding genes with the highest absolute protein synthesis rates^25^ (top 50; green) tend to have low Gini values, whereas structured ncRNAs (red) occupy high-Gini territory. Right, boxplots summarizing Gini for three classes (high-synthesis CDSs, other CDSs, ncRNAs). Lower Gini reflects more uniform contacts consistent with an unfolded, actively translating state; higher Gini reflects localised contacts characteristic of stable RNA structures.

**Supplementary Figure 3.**
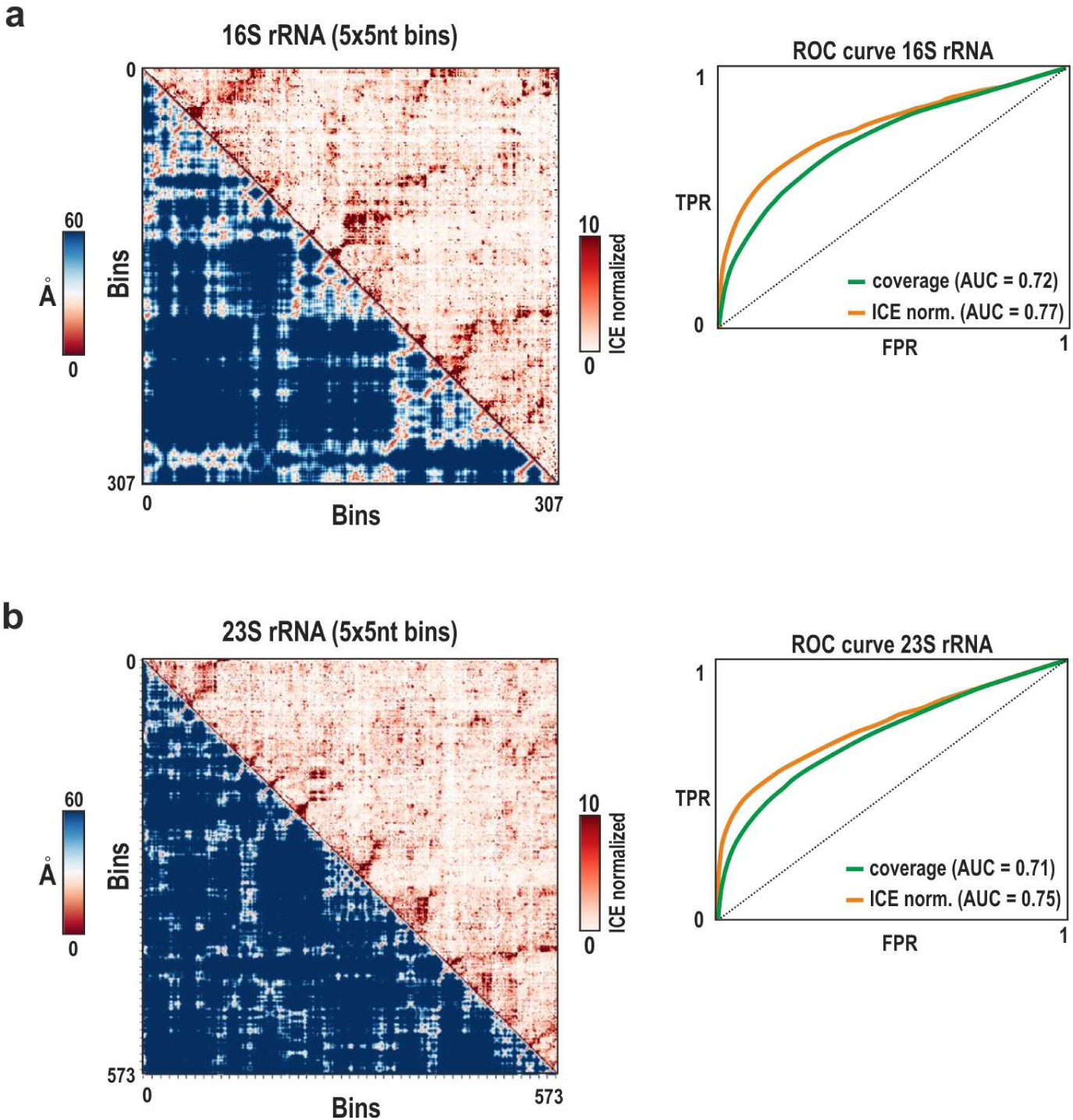
Benchmarking TRIC-seq structural data against known ribosomal RNA 3D structures. **a**, 16S rRNA; **b,** 23S rRNA, both binned at 5 × 5 nt. In each matrix, the lower triangle (blue scale) shows Euclidean distances (Å) between nucleotide bins derived from the crystal structure (PDB: 7ACJ), and the upper triangle (red scale) shows the corresponding ICE-normalized TRIC-seq contact frequencies. Right, ROC curves quantify how well TRIC-seq scores predict proximal nucleotide pairs (distance ≤ 25 Å). ICE normalization (orange) improves predictive power over raw data (green).

**Supplementary Figure 4.**
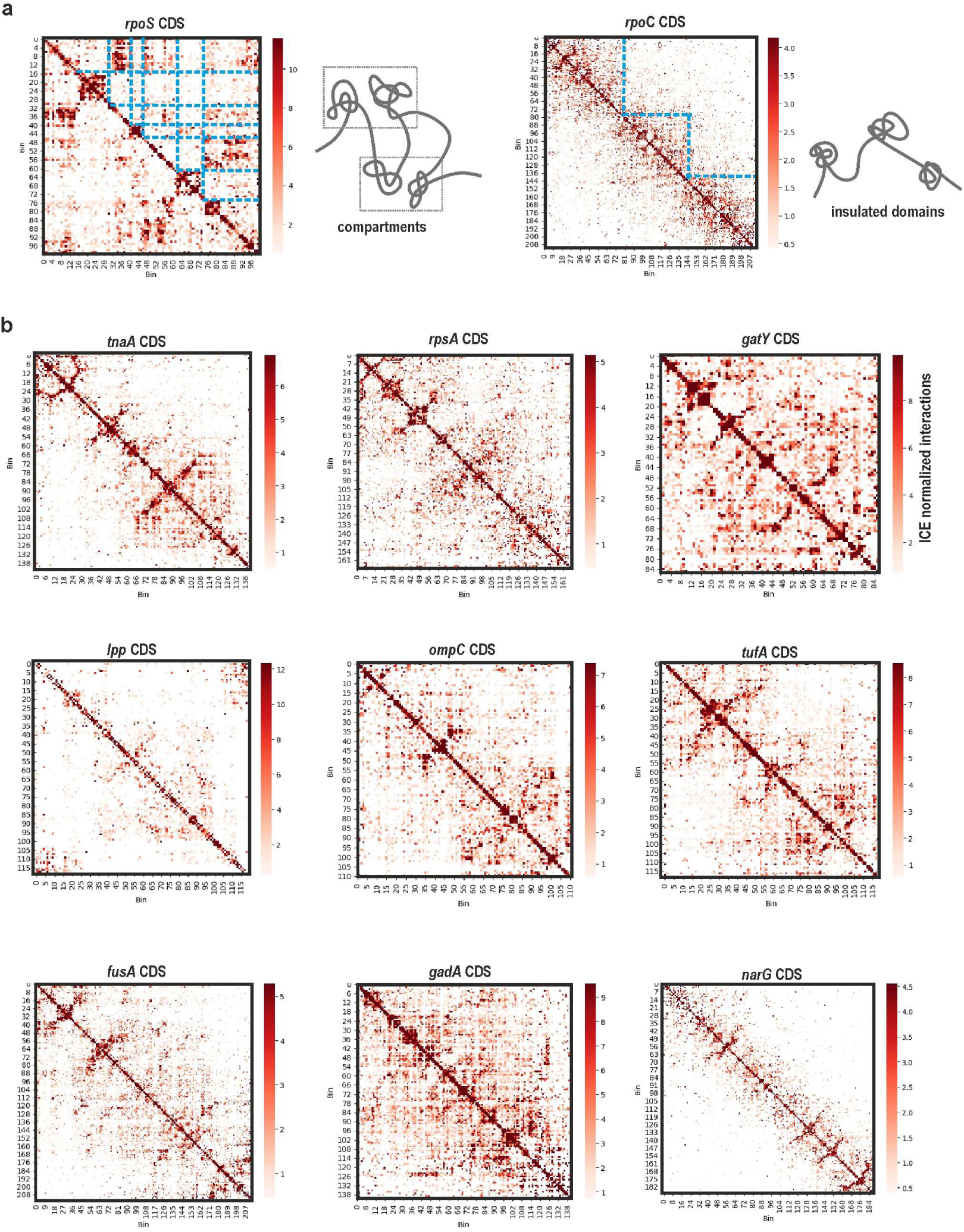
Intramolecular contact maps reveal diverse tertiary organisations of mRNAs. **a**, ICE-normalized contact maps for the *rpoS* and *rpoC* CDSs illustrate two distinct higher-order organisations. *rpoS* shows a compartmentalized architecture with checkerboard patterns, whereas *rpoC* resolves into a series of discrete domains along the diagonal, indicating multiple structurally insulated units. Schematic structure shown next to each map. **b,** Gallery of ICE-normalized maps for nine additional *E. coli* CDSs (*tnaA, rpsA, gatY, lpp, ompC, tufA, fusA, gadA, narG*). Each transcript exhibits a characteristic folding pattern, highlighting the broad structural diversity of bacterial mRNAs.

**Supplementary Figure 5.**
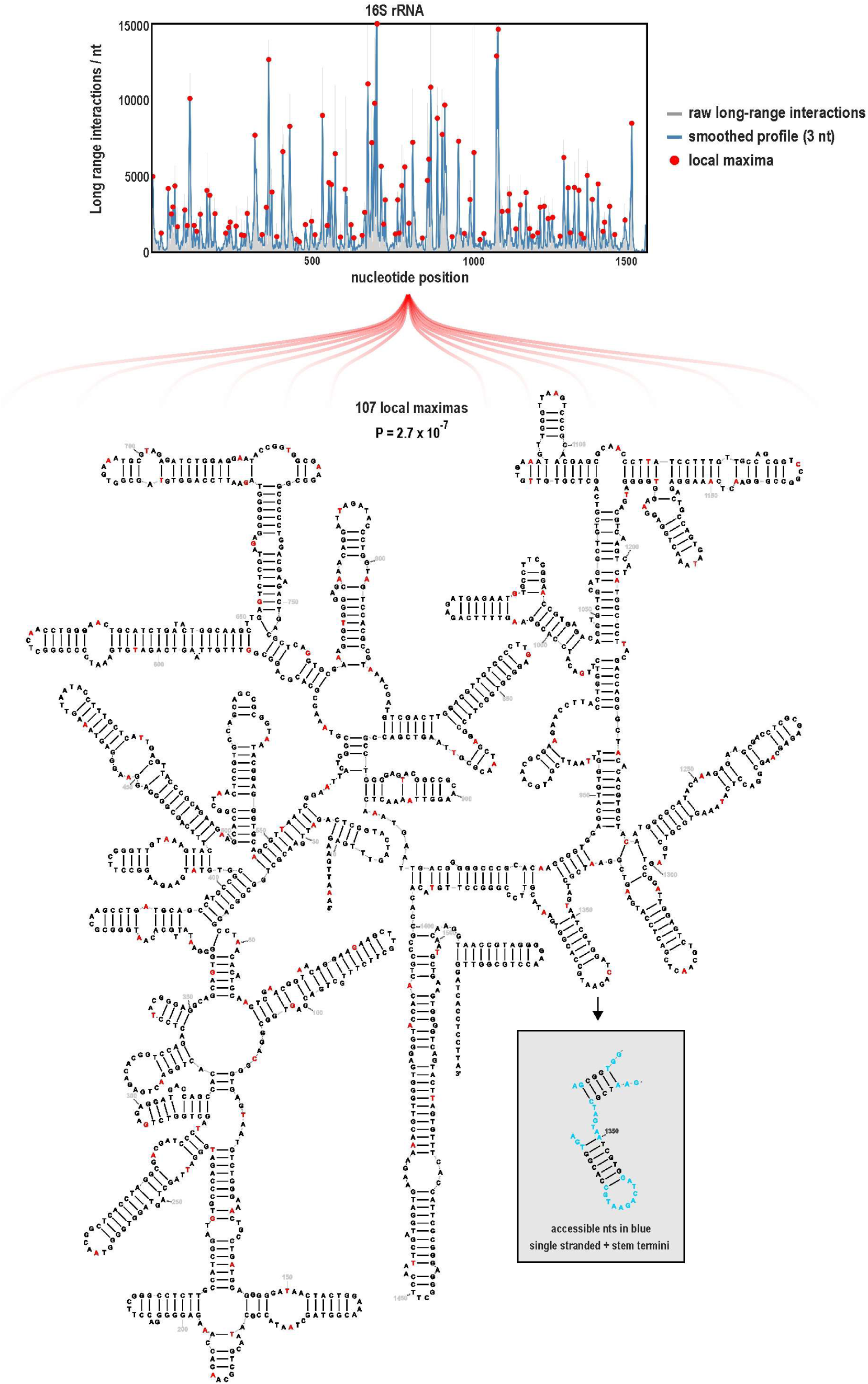
Identification of accessible regions in 16S rRNA. Top, 1D long-range ligation profile along 16S rRNA (grey bars, raw interactions per nucleotide; blue line, 3-nt moving average). Red dots mark local maxima (n = 107), interpreted as sites with highest ligation propensity. Bottom, secondary-structure diagram of 16S with these maxima highlighted in red. T represents uridine (U) in the RNA structure. A one-tailed hypergeometric test shows that peaks are significantly enriched at accessible nucleotides (unpaired bases and stem termini) relative to a uniform random placement (P = 2.7 × 10⁻⁷). Inset, example region with accessible nucleotides indicated in blue.

**Supplementary Figure 6.**
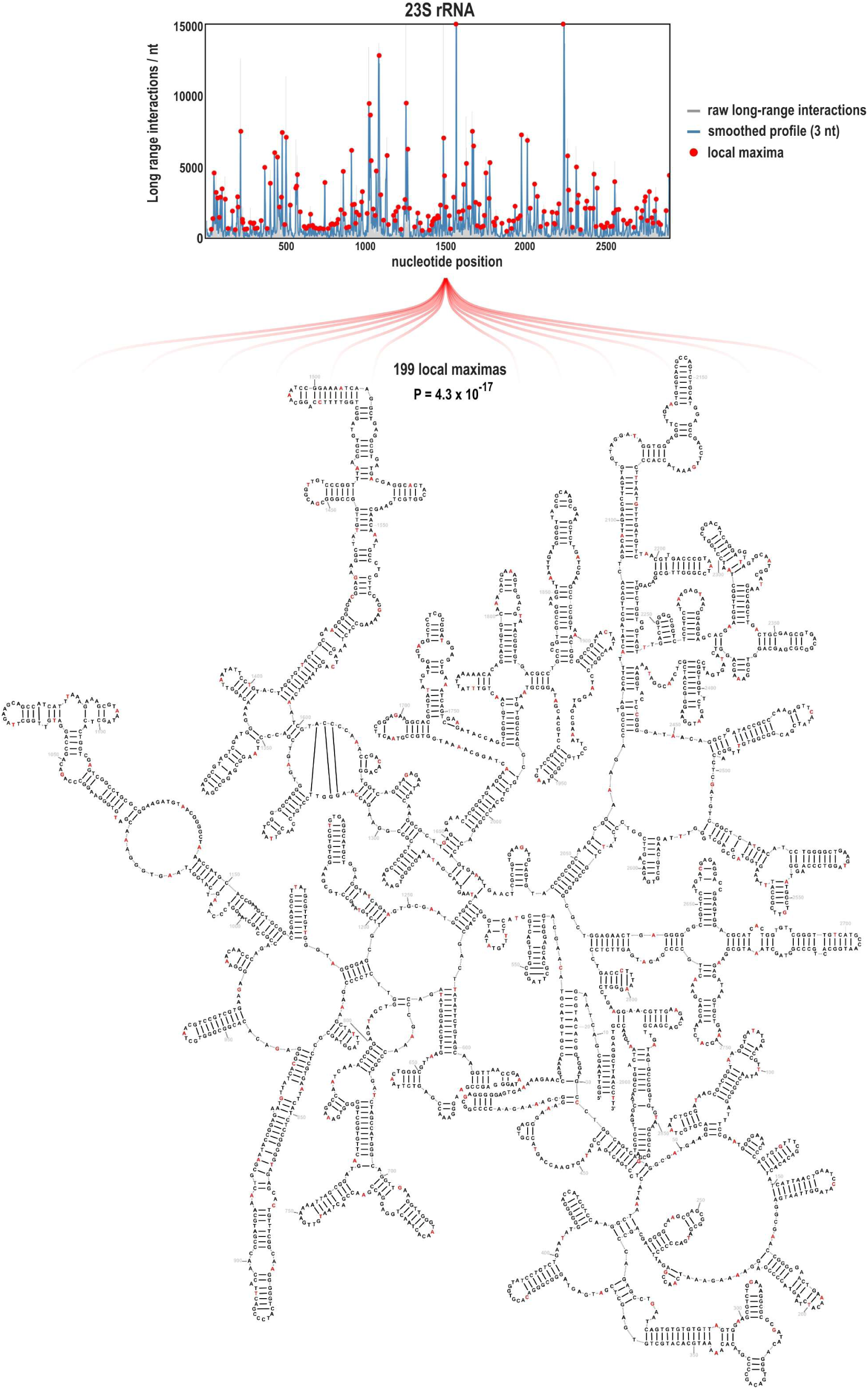
Identification of accessible regions in 23S rRNA. Top, 1D long-range ligation profile along 23S rRNA (grey bars, raw interactions per nucleotide; blue line, 3-nt moving average). Red dots mark local maxima (n = 199), i.e., positions with the highest ligation propensity. Bottom, secondary-structure diagram of 23S with these maxima highlighted in red. T represents uridine (U) in the RNA structure. A one-tailed hypergeometric test shows that peaks are significantly enriched at accessible nucleotides relative to a uniform-random placement (P = 4.3 × 10⁻¹⁷).

**Supplementary Figure 7.**
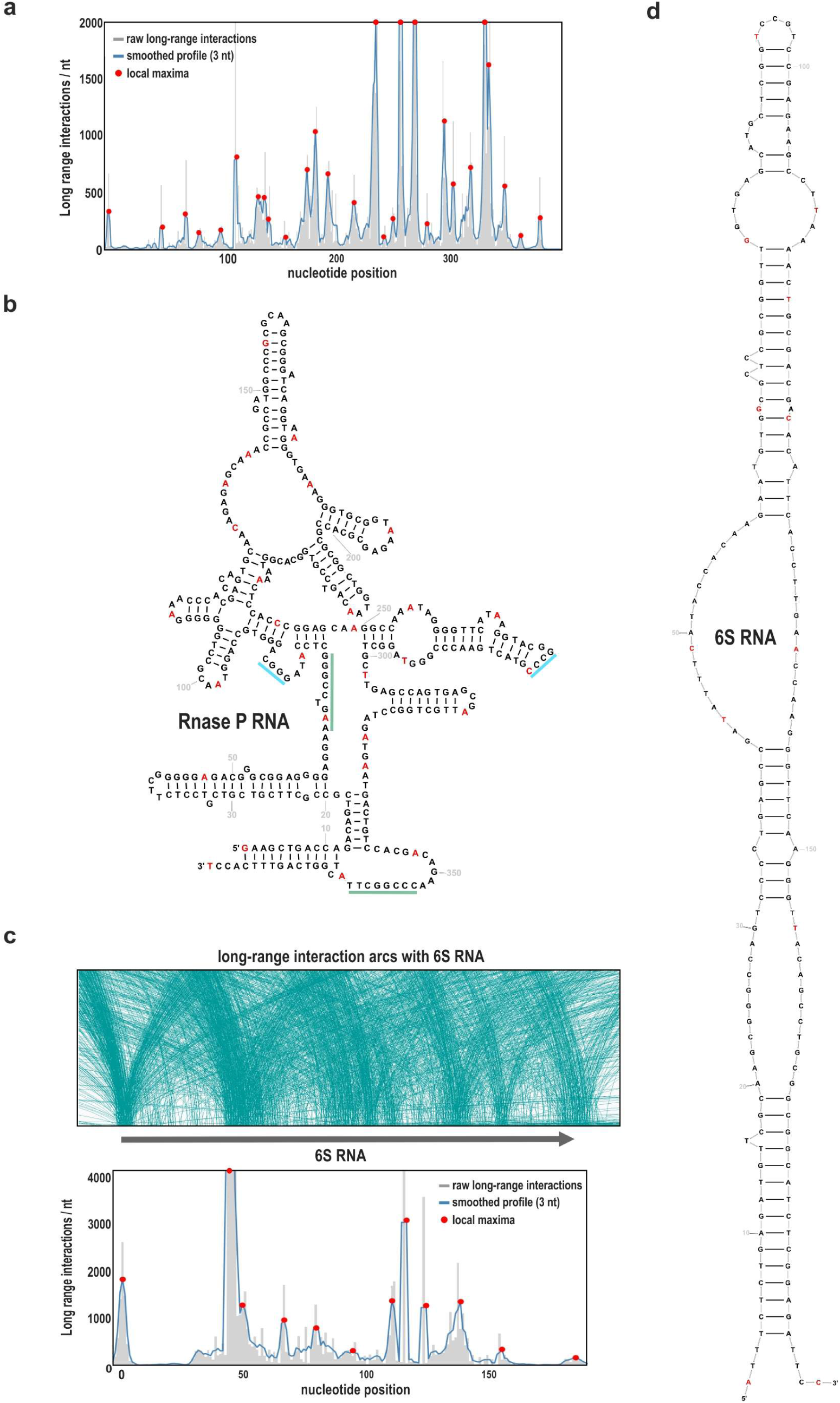
Accessible single-stranded regions in RNase P RNA and 6S RNA. **a**, 1D long-range ligation profile along *E. coli* RNase P RNA (grey bars, raw interactions per nucleotide; blue line, 3-nt moving average). Red dots mark local maxima, interpreted as highly accessible sites. **b,** Secondary-structure model of RNase P RNA with local maxima (from a) in red. Nucleotides that form the pseudoknot are highlighted in blue/cyan. **c,** Top, Integrative Genomics Viewer (IGV) snapshot showing long-range interaction arcs emanating from the 6S RNA (*ssrS*) locus (arcs shown from representative libraries to avoid clutter). Bottom, per-nucleotide long-range ligation profile for 6S RNA (grey bars, raw; blue line, 3-nt moving average) with local maxima in red. **d,** Secondary-structure model of 6S RNA with local maxima highlighted in red. T represents uridine (U) in the RNA structures.

**Supplementary Figure 8.**
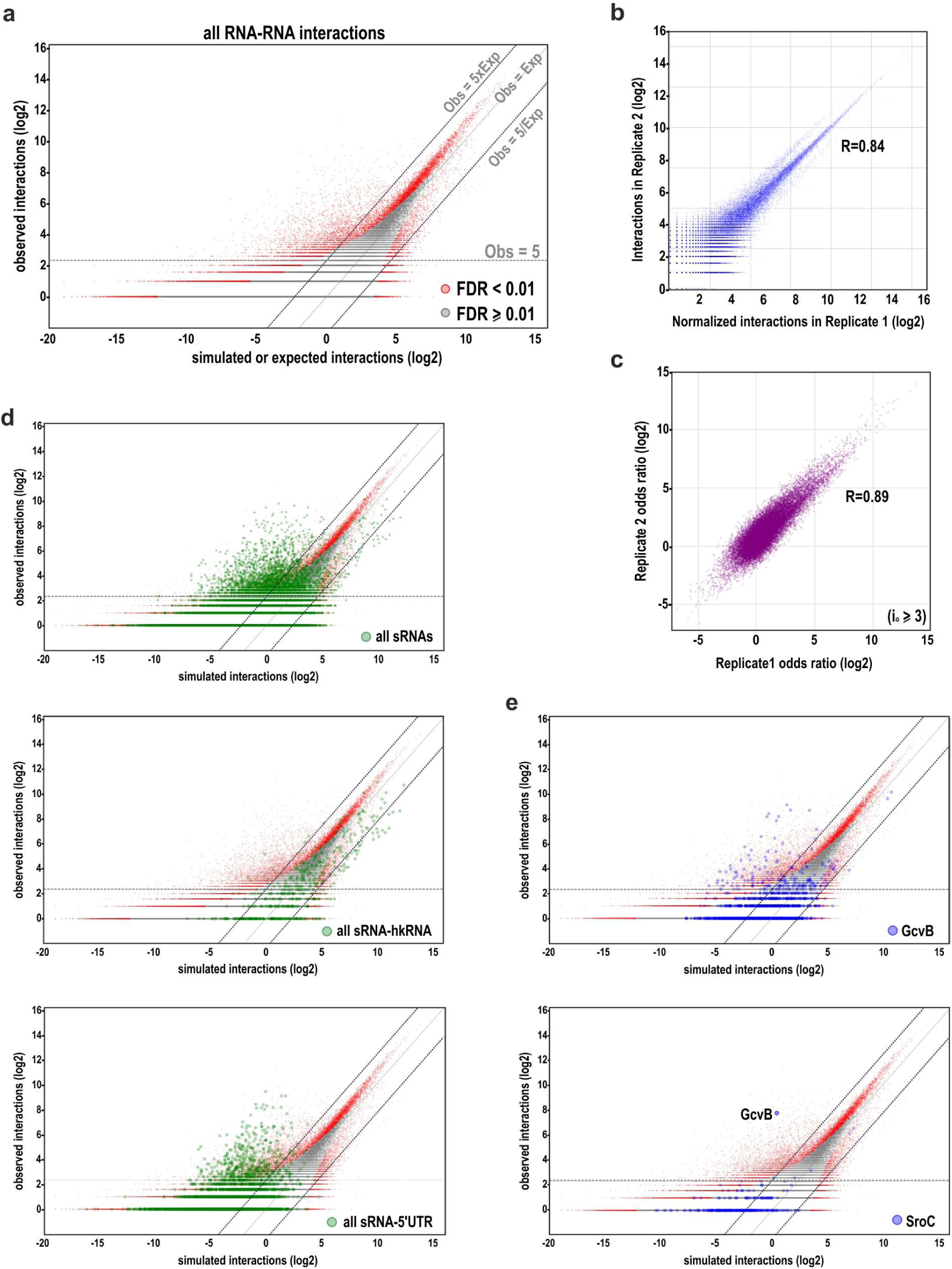
Statistical framework for identifying significant *trans*-interactions. **a**, Observed versus simulated (1 million interactomes using configuration-model) interaction counts for all RNA– RNA pairs (log scale). Points along the diagonal (Obs = Exp) reflect stochastic background. Interactions passing multiple-testing correction (FDR < 0.01, red) are strongly enriched, frequently above the Obs = 5 × Exp line. **b,** Reproducibility of interaction counts between biological replicates (log scale). Replicate 1 yielded 7.96 M chimeras and Replicate 2 yielded 2.74 M chimeras; Replicate 1 was normalized to the total of Replicate 2 for this comparison. Only pairs with ≥1 interaction in both datasets are shown; Pearson R = 0.84. **c,** Reproducibility of odds ratios (Of) between replicates for pairs detected in both. Despite the ∼3× difference in total chimeras, odds ratio (*O^f^*) values are highly consistent (R = 0.89), underscoring the robustness of the enrichment metric to library size. **d,** Same as (a), highlighting subsets involving sRNAs. All sRNA interactions (green) and specifically sRNA– 5′UTR pairs show strong enrichment, consistent with regulatory base-pairing, whereas sRNA–hkRNA pairs are generally de-enriched (most points below the diagonal), as expected for non-interacting combinations. **e,** Same as (a), with pairs involving GcvB (top) or SroC (bottom) highlighted (blue). The selectively enriched partners delineate their regulons, illustrating how the framework resolves individual sRNA networks.

**Supplementary Figure 9.**
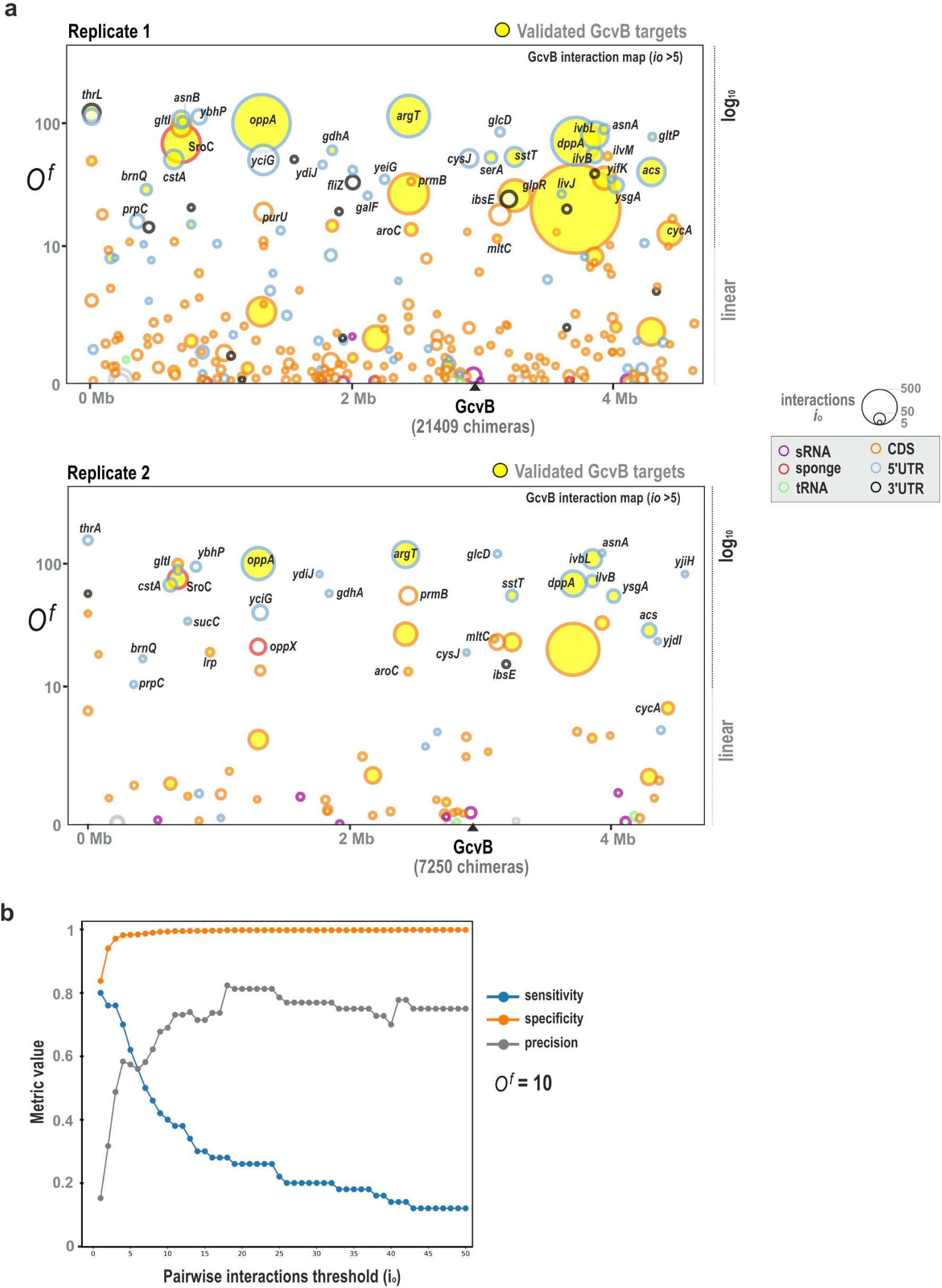
Global interaction maps for GcvB and performance metrics. **a**, Global interaction maps for the sRNA GcvB from two biological replicates (Replicate 1: 21409 chimeras; Replicate 2: 7250 chimeras). Each circle is a partner RNA plotted by genomic coordinate (x) and enrichment odds ratio *O^f^*(y); circle area ∝ raw interaction count i_o_ and edge colour denotes RNA class. All annotated features (5′UTR/CDS/3′UTR) belonging to the 56 literature-validated^30^ GcvB targets are highlighted in yellow. **b,** Performance metrics for GcvB target prediction as a function of the minimum pairwise interaction count threshold (iₒ), with odds ratio cut-off fixed at 10. Curves show sensitivity, specificity and precision computed against the 56 validated targets. Specificity rapidly approaches 1 and precision plateaus at low iₒ, indicating that few chimeric reads are sufficient for high-confidence target calls, whereas sensitivity declines as the threshold increases.

**Supplementary Figure 10.**
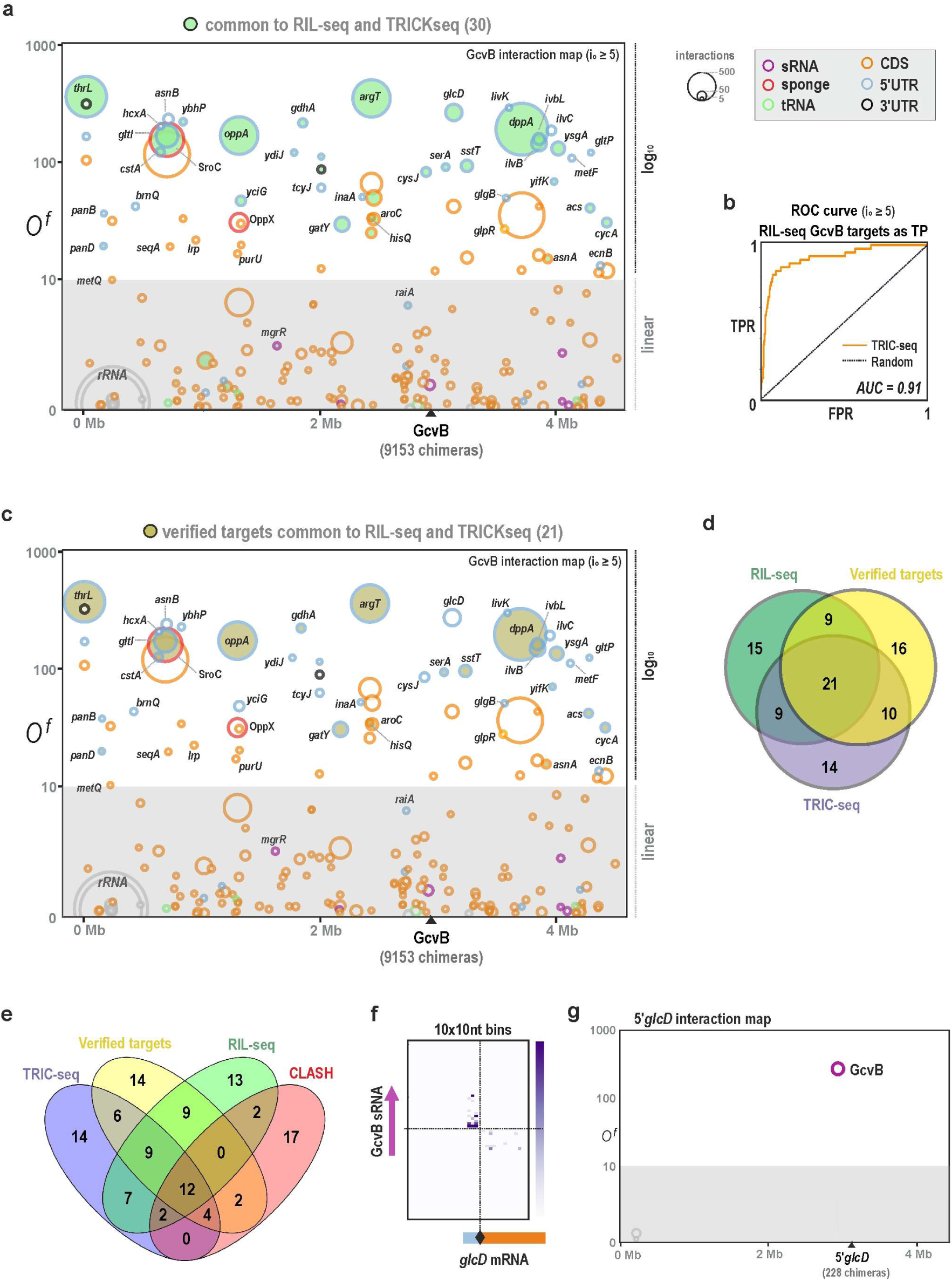
Comparison of TRIC-seq GcvB targets with RIL-seq and CLASH. **a**, A global interaction map of GcvB highlighting the 30 GcvB targets that are common to both the RIL-seq and TRIC-seq datasets in green. **b,** ROC curve using *O^f^* to predict the RIL-seq GcvB target set; AUC = 0.91, indicating strong concordance. **c,** Global interaction map of GcvB highlighting the 21 experimentally validated GcvB targets that were successfully identified by both RIL-seq and TRIC-seq. **d, e,** Venn diagrams comparing the target sets from TRIC-seq, RIL-seq, CLASH, and the set of experimentally validated targets. The diagrams illustrate both the significant overlap between the methods and the unique targets identified by each approach. **f,** An Inter-RNA heat map (10 × 10-nt bins) for the GcvB–*glcD* interaction, pinpointing binding around the Shine–Dalgarno region. **g,** *glcD* 5’UTR global interaction map confirms the highly specific interaction with GcvB.

**Supplementary Figure 11.**
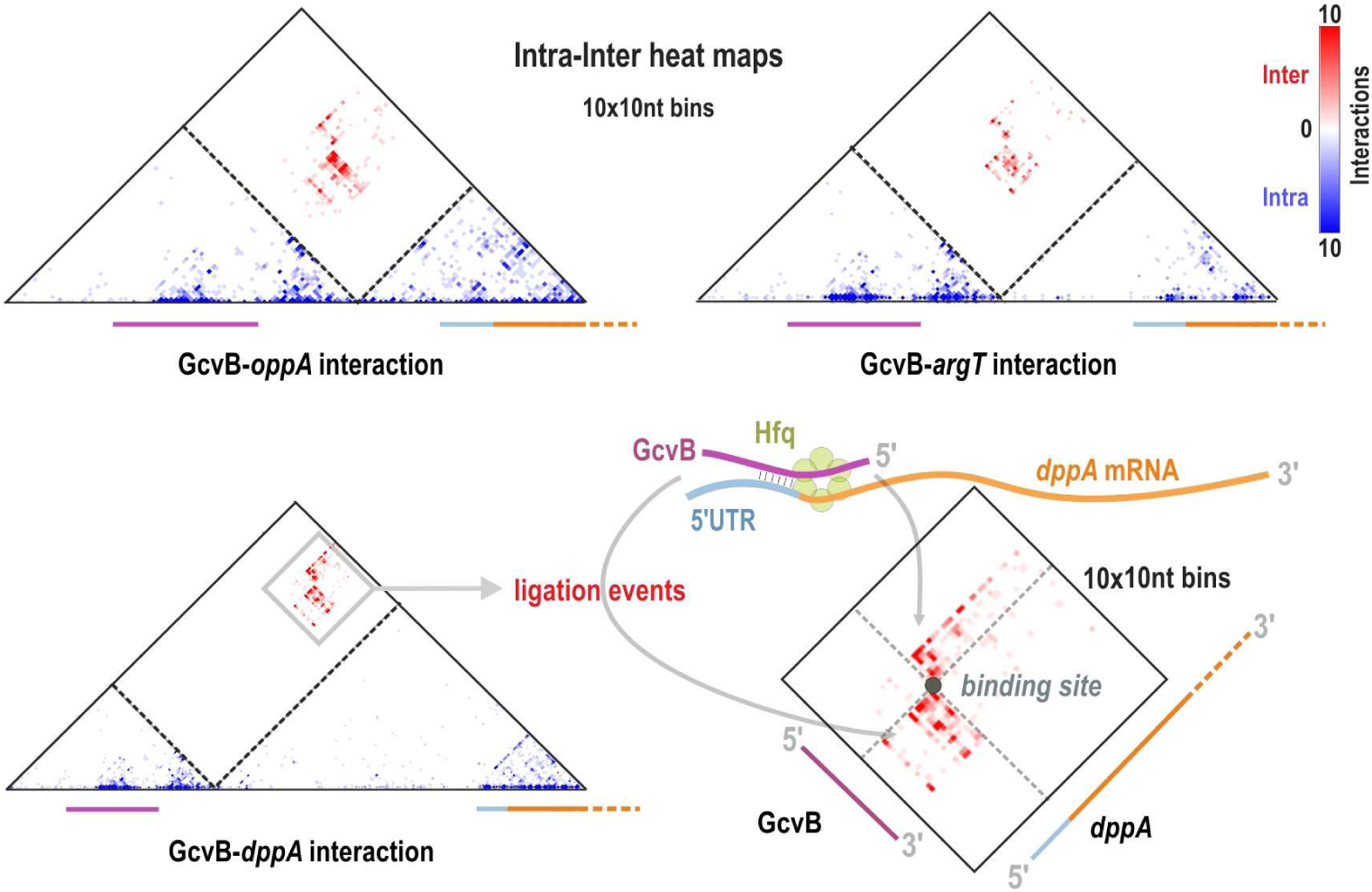
Composite maps resolve simultaneous RNA structure and interaction. Intra–inter contact maps (10 × 10-nt bins) for GcvB with three canonical targets—*oppA*, *argT*, and *dppA*. Each triangular matrix concatenates the two RNAs along the axes. Blue signals report intramolecular contacts within each RNA (structural), whereas red signals mark intermolecular ligations at the base-pairing interface. Schematic of the GcvB–*dppA* interaction illustrating proximity-ligation events captured by TRIC-seq.

**Supplementary Figure 12.**
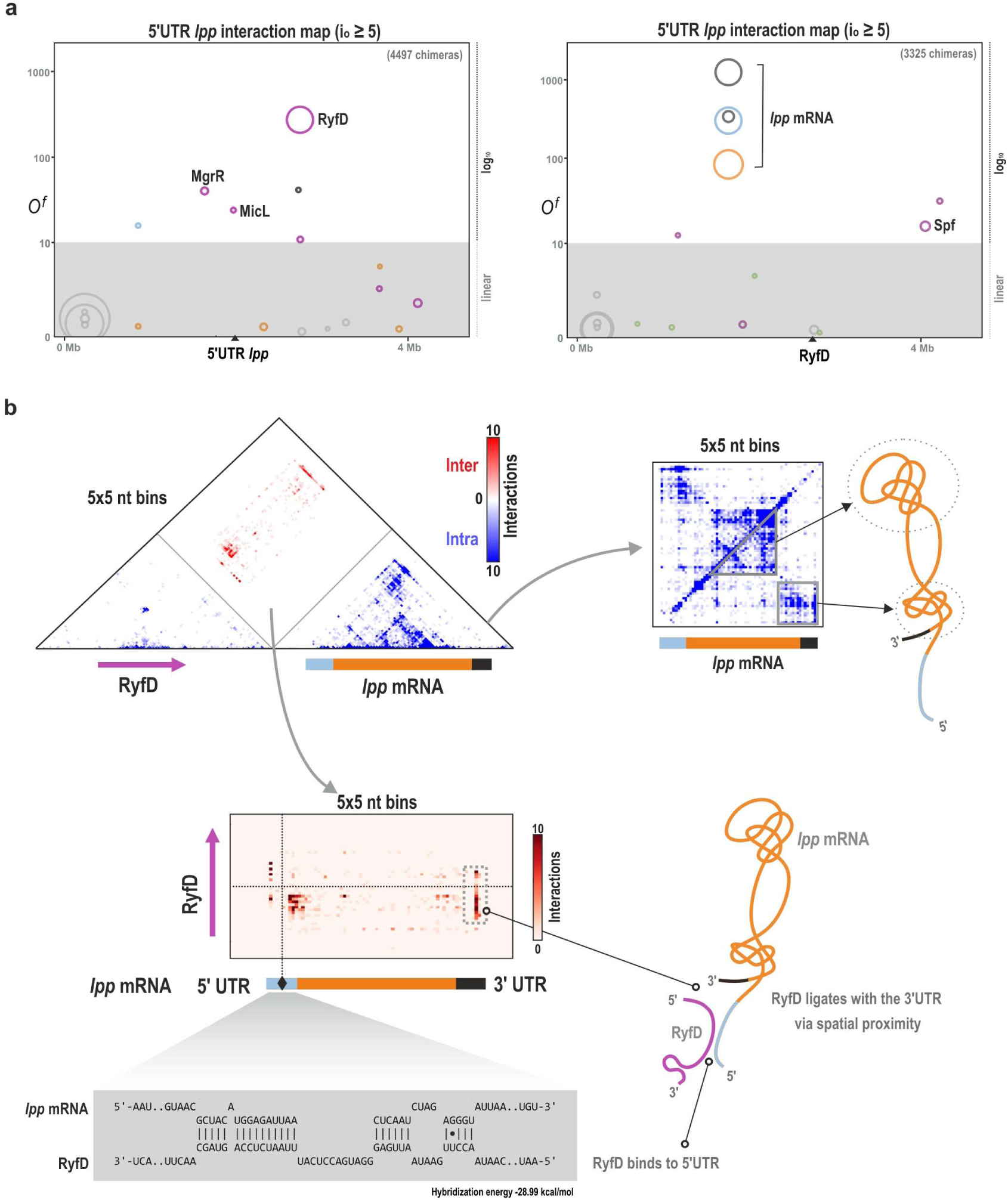
Structure-driven interaction between the RyfD sRNA and *lpp* mRNA. **a**, Global interaction maps for the *lpp* 5’UTR (left; i_o_ ≥ 5; 4497 chimeras) and for RyfD (right; i_o_ ≥ 5; 3325 chimeras). Circles are partners plotted by genomic coordinate (x) and enrichment odds ratio *O^f^*(y); circle area ∝ i_o_ and colour encodes RNA class. Both maps reveal a strong, specific interaction between RyfD and the full-length monocistronic *lpp* transcript (5’UTR, CDS, and 3’UTR). **b,** Detailed analysis of the RyfD–*lpp* pairing. Top left, intra– inter heat map (5 × 5-nt bins) simultaneously showing intramolecular contacts (blue) within each RNA and the intermolecular interface (red). Top right, the *lpp* intramolecular map alone reveals a compact tertiary fold. Bottom, schematic and predicted duplex (IntaRNA) indicate that RyfD base-pairs to the *lpp* 5’UTR; the global 3D fold of *lpp* brings its 3’UTR into spatial proximity with the bound RyfD, explaining additional ligations to the 3’UTR that likely arise from spatial proximity rather than direct base-pairing.

**Supplementary Figure 13.**
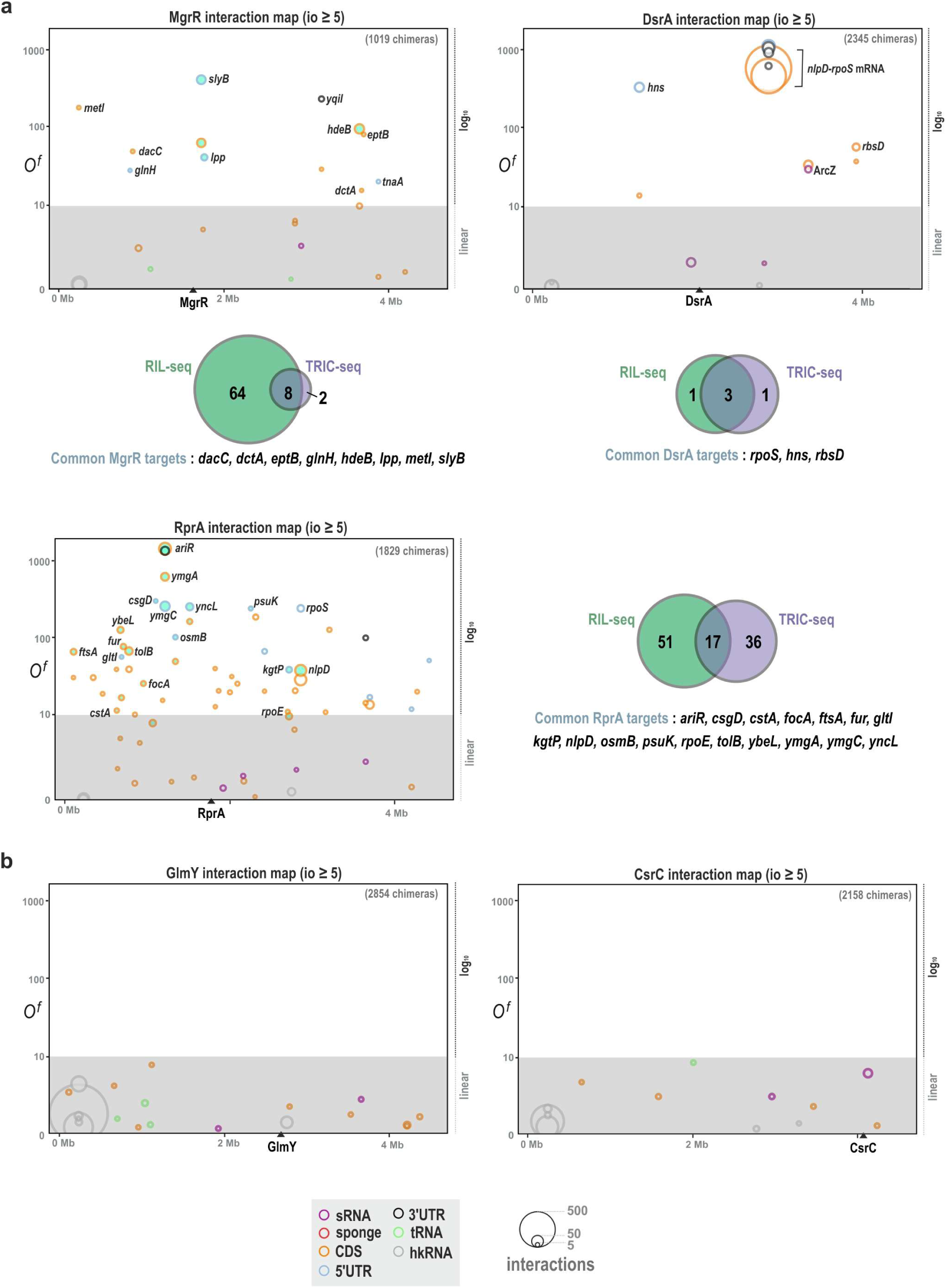
TRIC-seq specifically identifies sRNA targets. **a**, Global interaction maps for MgrR (i_o_ ≥ 5; 1019 chimeras), DsrA (i_o_ > 5; 2345 chimeras), and RprA (i_o_ ≥ 5; 1829 chimeras). Each circle represents a partner RNA plotted by genomic coordinate (x) and enrichment odds ratio **O**f (y; log₁₀); circle area ∝ raw interaction count (i_o_) and edge colour encodes RNA class (legend). Venn diagrams summarize overlap between TRIC-seq calls and RIL-seq targets (bottom right of each panel). For example, common MgrR targets include *dacC, dctA, eptB, glnH, hdeB, lpp, metI, slyB*; common DsrA targets include *rpoS, hns, rbsD*; common RprA targets include *ariR, csgD, cstA, focA, ftsA, fur, gltI, kgtP, nlpD, osmB, psuK, rpoE, tolB, ybeL, ymgA, ymgC, yncL*. In the MgrR and RprA maps, RIL-seq targets are filled green for clarity. **b,** Global interaction maps for protein-sequestering sRNAs GlmY (i_o_ ≥ 5; 2854 chimeras) and CsrC (i_o_ ≥ 5; 2158 chimeras). In contrast to MgrR, DsrA and RprA, these sponges show no significant RNA partners, underscoring TRIC-seq specificity and its ability to distinguish mRNA targeting sRNAs from protein-sequestering sRNAs.

**Supplementary Figure 14.**
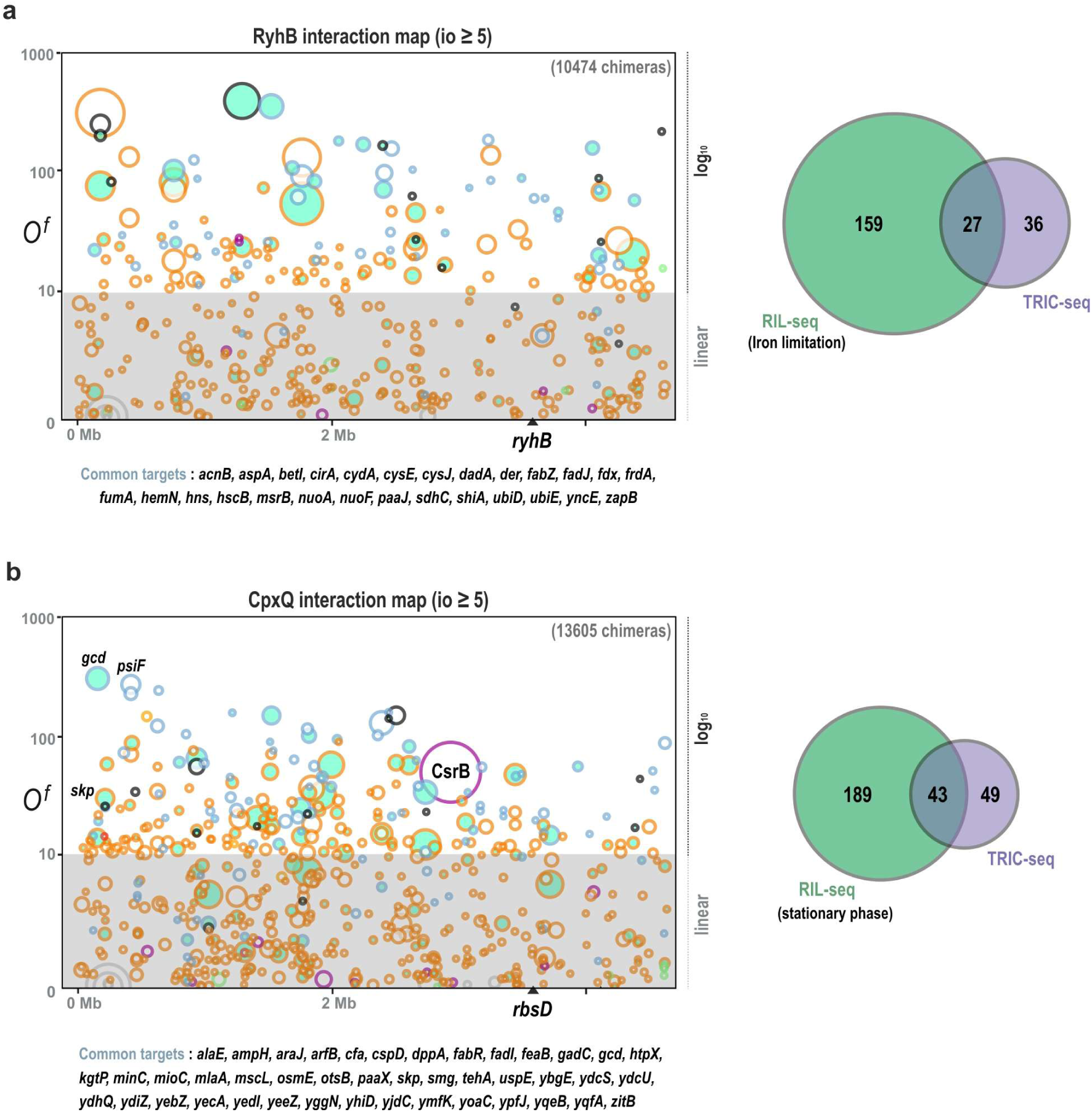
TRIC-seq captures the large regulons of RyhB and CpxQ sRNAs. **a**, Global interaction map for the iron-responsive sRNA RyhB (i_o_ ≥ 10; 10474 chimeras). The regulon comprises dozens of mRNAs involved in iron homeostasis and metabolism. Venn diagram (right) shows substantial overlap with RIL-seq targets. Note: for RyhB, the RIL-seq comparison set is from iron-limitation conditions^23^ (the stationary-phase RIL-seq data used for other sRNAs contained few RyhB interactions). Targets from RIL-seq are filled green on the map. **b,** Global interaction map for CpxQ (i_o_ ≥ 5; 13605 chimeras) highlighting its extensive regulon. The Venn diagram confirms strong concordance with stationary-phase RIL-seq data. RIL-seq targets are filled green.

**Supplementary Figure 15.**
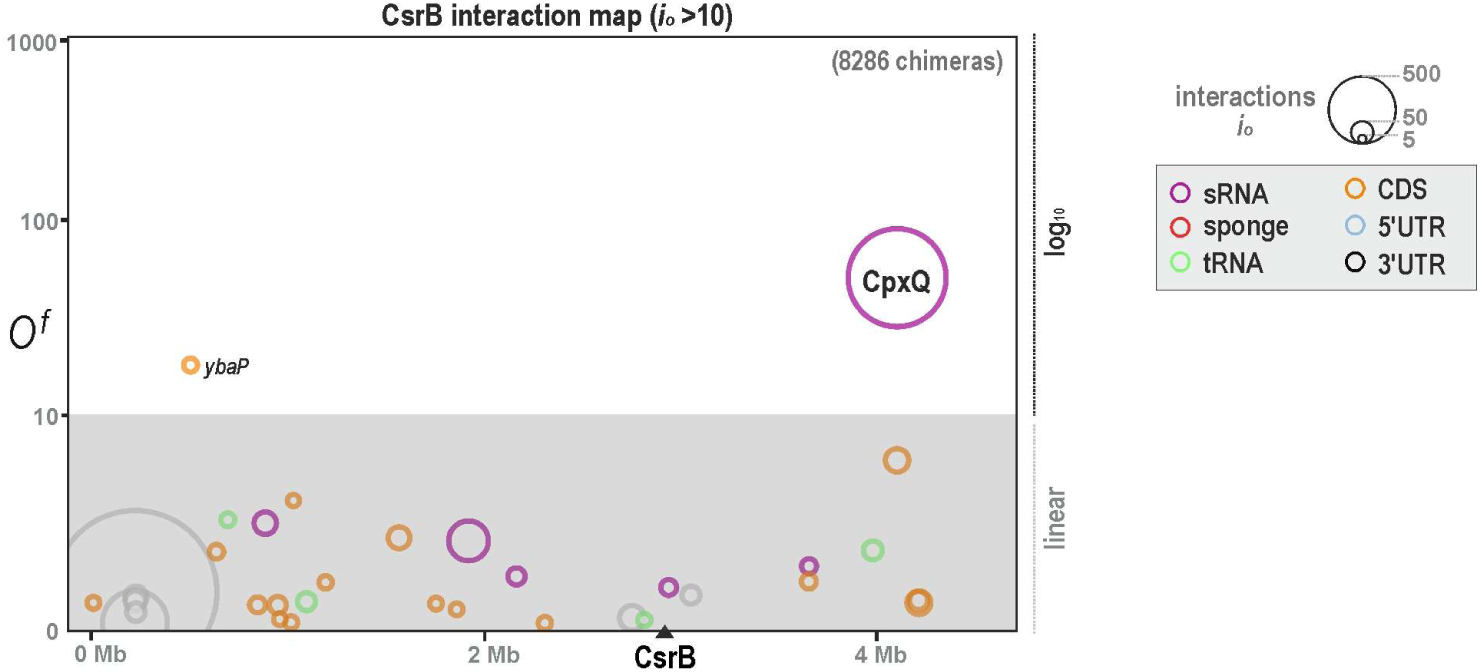
Global interaction map of CsrB reveals a specific sRNA–sRNA contact with CpxQ. Global interaction map for CsrB (i_o_ ≥ 10; 8286 chimeras). Each circle is a partner RNA plotted by genomic coordinate (x) and enrichment odds ratio *O^f^* (y); circle area ∝ io and edge colour encodes RNA class. As expected for a protein-sequestering RNA that primarily sequesters the protein CsrA, CsrB shows an almost complete absence of RNA partners. A single, specific high-odds interaction with the sRNA CpxQ stands out.

**Supplementary Figure 16.**
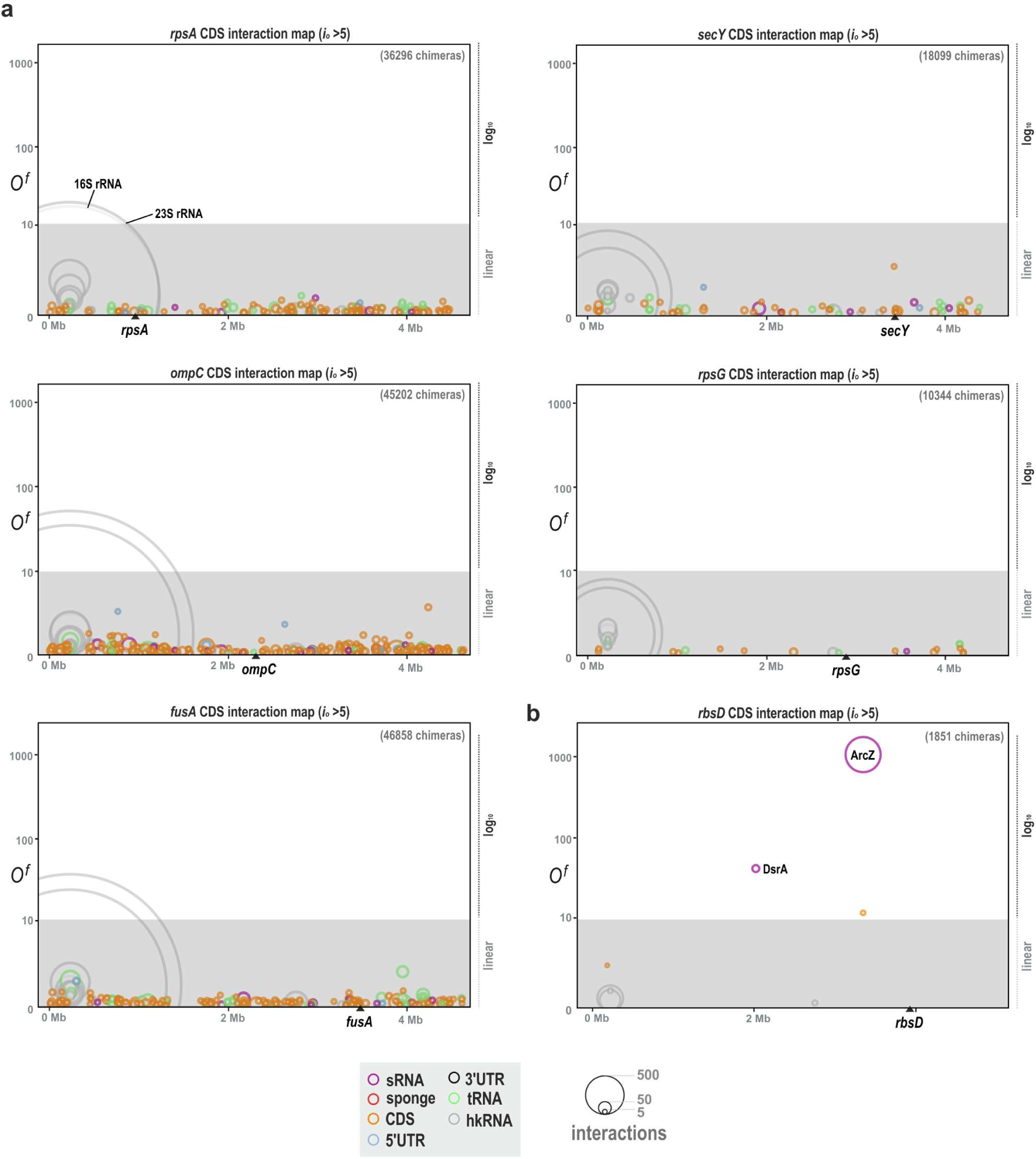
Interaction profiles of selected CDSs. **a**, Global interaction maps (i_o_ ≥ 5) for highly expressed CDSs (*rpsA, secY, ompC, rpsG, fusA*). Circles are partners plotted by genomic coordinate (x) and enrichment odds ratio *O^f^* (y); circle area ∝ io and edge colour encodes RNA class. Profiles are dominated by contacts with hkRNAs (rRNAs, tRNAs) consistent with active translation and show numerous low-odds CDS–CDS contacts expected from nonspecific encounters in a crowded cytoplasm. **b,** Global map for the *rbsD* CDS shows a highly specific profile, with strong, enriched contacts to the sRNAs ArcZ and DsrA. This pattern supports the model that *rbsD* ORF acts as an sRNA sponge, sequestering ArcZ/DsrA and thereby limiting their activation of *rpoS* translation^41^.

**Supplementary Figure 17.**
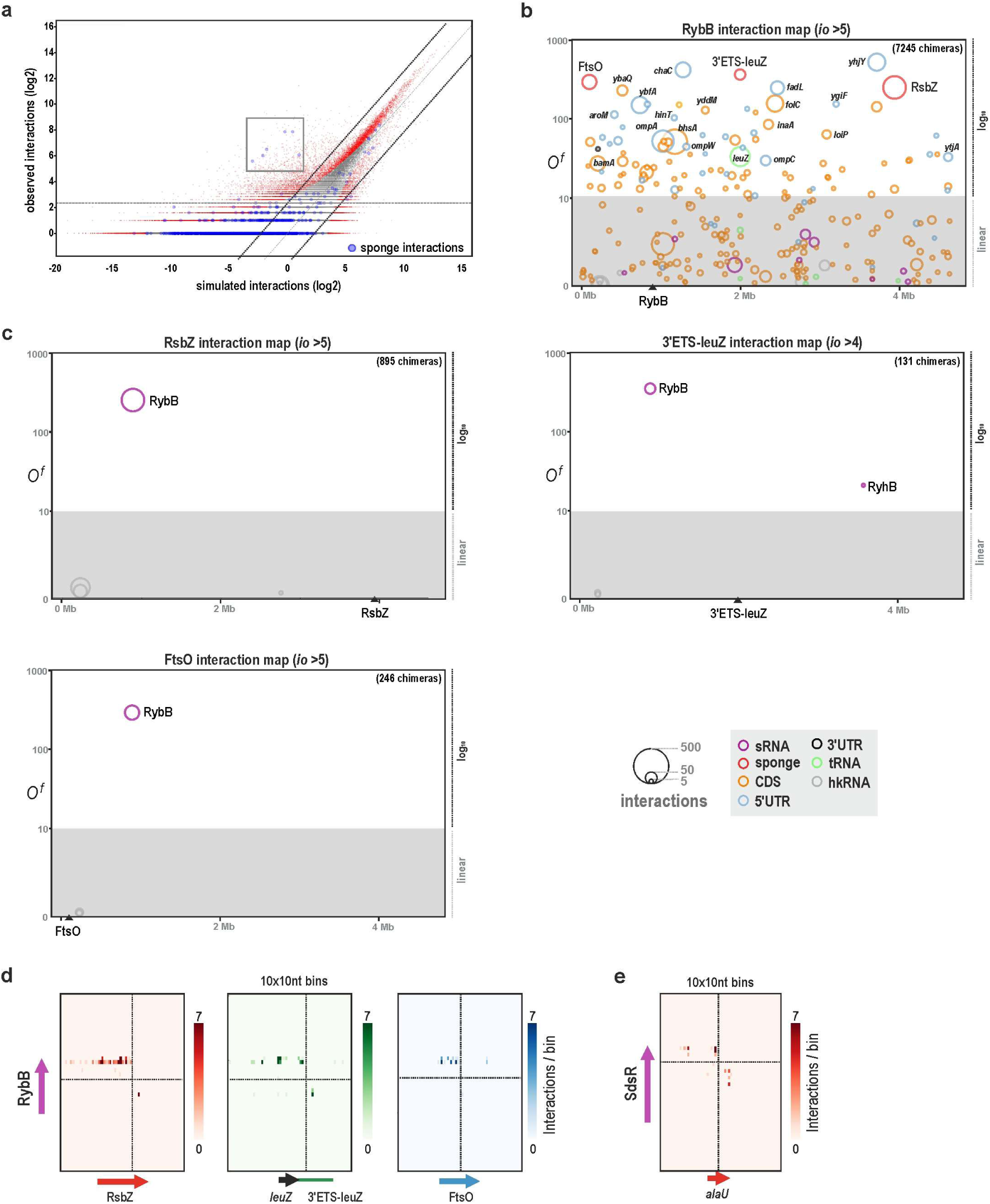
TRIC-seq captures the extreme specificity of RNA sponges. **a**, Observed-versus-simulated (configuration-model) interaction counts for all RNA–RNA pairs (log₂). Known sRNA–sponge pairs (blue) are strong outliers (boxed), falling well above the enrichment lines and among the most statistically significant interactions genome-wide (background significant pairs in red, FDR < 0.01). **b,** Global interaction map for RybB (i_o_ ≥ 5; 7245 chimeras). Circles are partners plotted by genomic coordinate (x) and odds ratio *O^f^* (y); area ∝ i_o_ and colour encodes RNA class. RybB shows specific contacts to its sponges (RsbZ, 3′ETS-leuZ, FtsO) and to multiple outer-membrane protein mRNAs. **c,** Global maps for the three sponges targeting RybB: RsbZ (i_o_ ≥ 5; 895 chimeras), 3′ETS-leuZ (i_o_ ≥ 4; 131 chimeras), and FtsO (i_o_ ≥ 5; 246 chimeras). Each shows an exclusive, high-adds interaction with RybB and minimal off-target contacts, consistent with dedicated sponge function. **d,** Inter-RNA heat maps (10x10 nt bins) resolving the interaction between RybB and its three sponge RNAs: RsbZ, 3’ETS-leuZ, and FtsO. **e,** Inter-RNA heat map revealing a novel, specific interaction between the stress-responsive sRNA SdsR and the *alaU* tRNA.

**Supplementary Figure 18.**
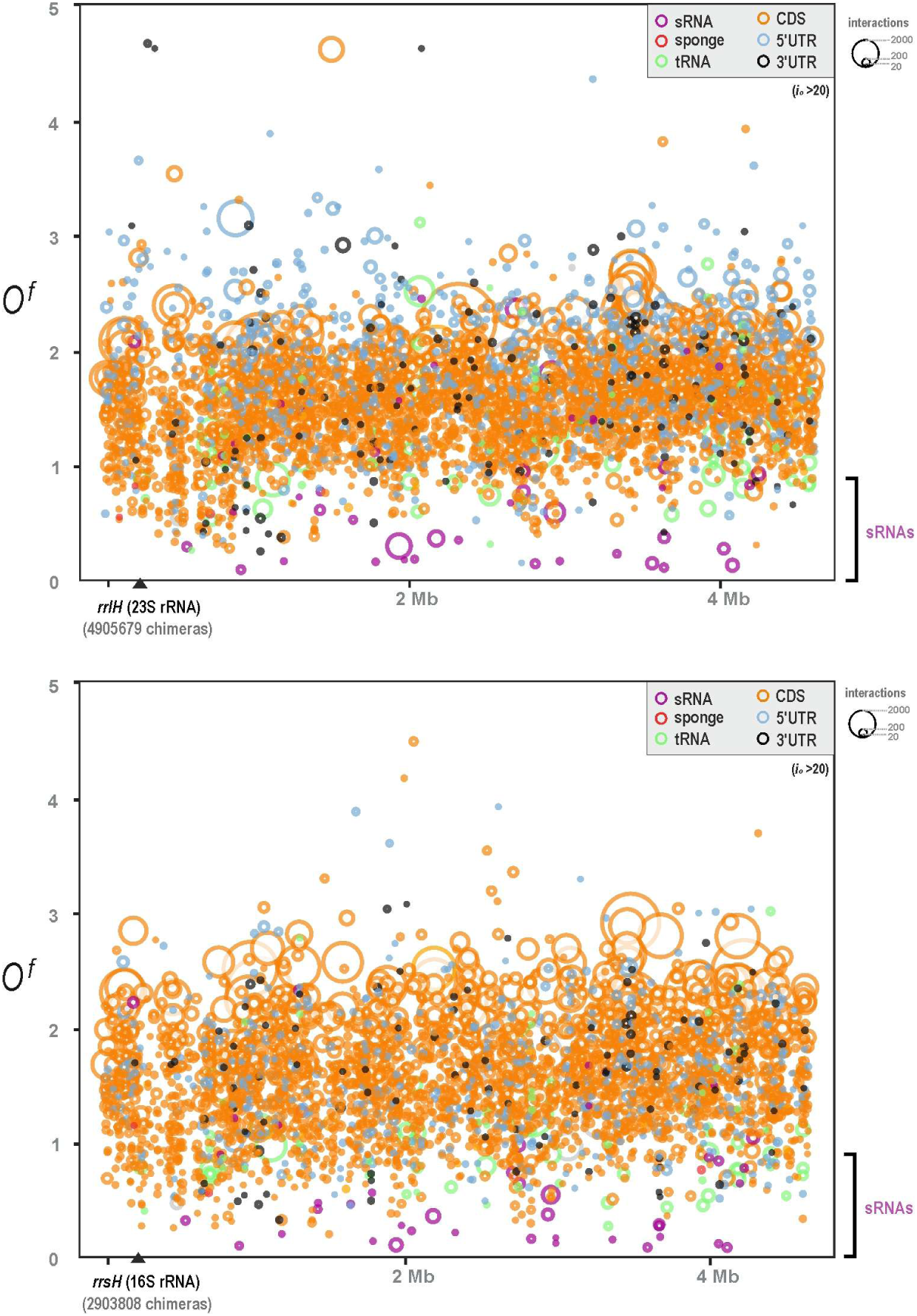
Global interaction maps of ribosomal RNA subunits. Global interaction maps for 23S rRNA (top; i_o_ ≥ 20) and 16S rRNA (bottom; i_o_ ≥ 20). As expected for core translation components, both rRNAs show widespread contacts with CDSs (orange) across the genome, whereas sRNAs (magenta) exhibit the lowest enrichment with both subunits. Note - rRNAs interact with a large fraction of transcriptome during active translation (tRNAs, UTRs, CDSs) and because the null model accounts for node degree, their expected interactions are also high, so *O^f^* tend to be around 1 even when absolute counts are large. In contrast, sRNAs contact a restricted subset of mRNAs; the null expectation is low, allowing specific targets to achieve high *O^f^* values.

**Supplementary Figure 19.**
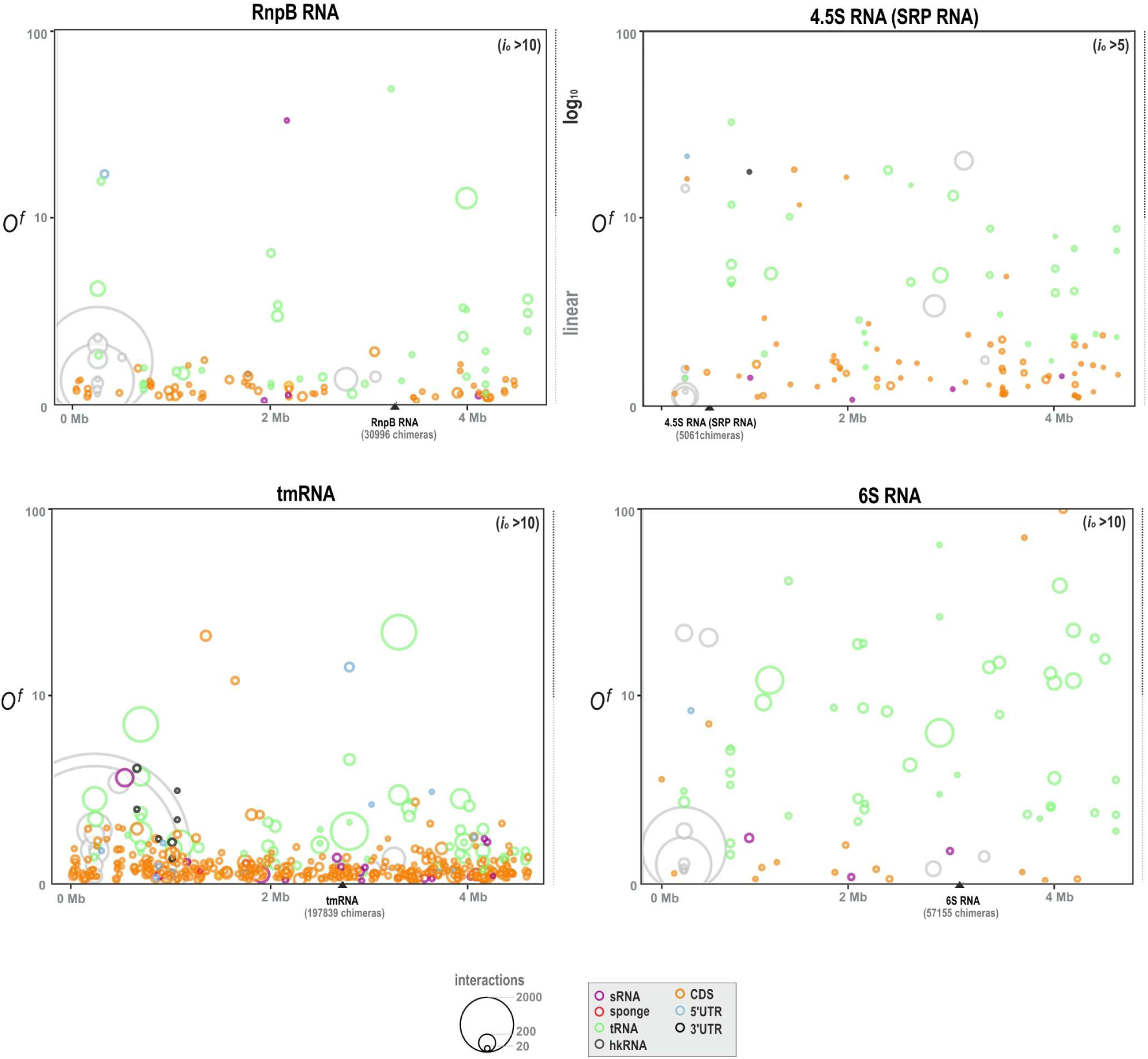
Interaction profiles of core housekeeping RNAs. Global interaction maps for four essential *E. coli* non-coding RNAs—RnpB (RNase P), 4.5S RNA (SRP RNA), tmRNA (SsrA), and 6S RNA (SsrS)—show distinct partner spectra. Each circle denotes a partner RNA plotted by genomic coordinate (x) and enrichment odds ratio *O^f^* (y); circle area ∝ raw interaction count (i_o_) and colours indicate RNA class (legend). RnpB (i_o_ ≥ 10; 30996 chimeras): contacts are strongly enriched for tRNAs, consistent with RNase P’s role in processing precursor tRNAs. 4.5S RNA (i_o_ ≥ 5; 5061 chimeras): partners include both CDSs and tRNAs, aligning with SRP’s co-translational targeting of nascent membrane proteins and their mRNAs. tmRNA (i_o_ ≥ 10; 197839 chimeras): widespread interactions with CDSs and tRNAs reflect its broad role in ribosome rescue and trans-translation. 6S RNA (i_o_ ≥ 10; 57155 chimeras): shows multiple high-odds contacts with tRNAs. This was an unexpected finding, as 6S RNA has not previously been implicated in translation-related processes, warranting further investigation.

**Supplementary Figure 20.**
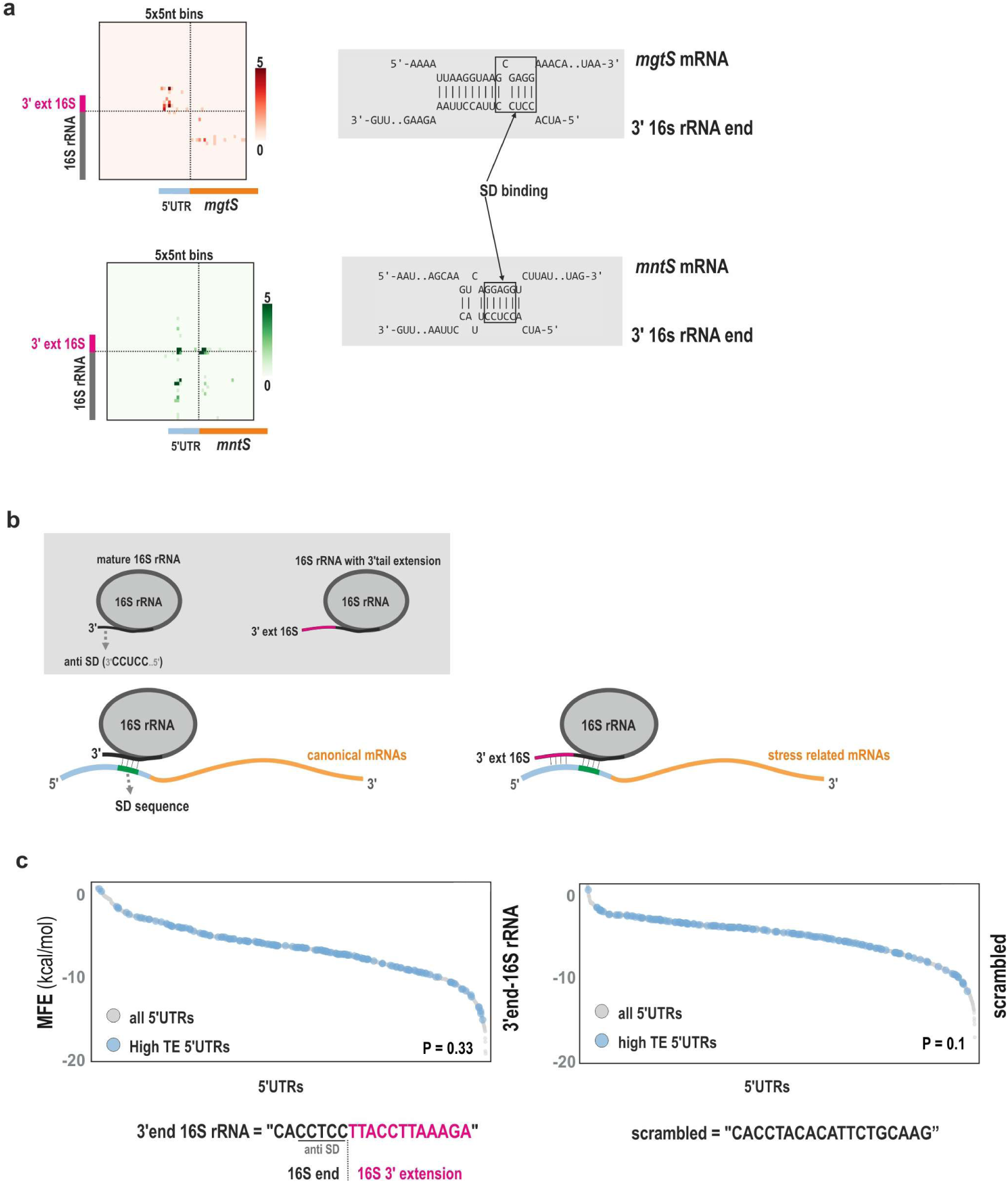
16S rRNA 3′-extension preferentially binds stress-related mRNAs. **a**, Inter-RNA heat maps (5 × 5-nt bins; left) and predicted duplexes (right) showing contacts between the 16S rRNA 3′ extension and the 5’UTRs of *mgtS* and *mntS*, which encode small proteins involved in Mg²⁺ and Mn²⁺ homeostasis, respectively. **b,** Left, canonical initiation: the mature 16S rRNA anti–Shine–Dalgarno (anti-SD) pairs with an mRNA SD. Right, tethering mechanism: the extended 3′ tail of 16S rRNA forms a longer duplex with selected stress-related 5’UTRs, potentially priming them for translation. **c,** Predicted duplex stability (minimum free energy, MFE) between the authentic 3′ end of 16S rRNA (sequence shown below) and all *E. coli* 5’UTRs compared to the subset with high translation efficiency (TE). No significant difference is observed (P = 0.33), indicating the 3′-tail interaction is not biased toward highly translated transcripts. Right panel, scrambled-tail control yields a similarly non-significant difference. Sequences used for predictions (bottom) indicate the anti-SD and the 3′-extension segments.

**Supplementary Figure 21.**
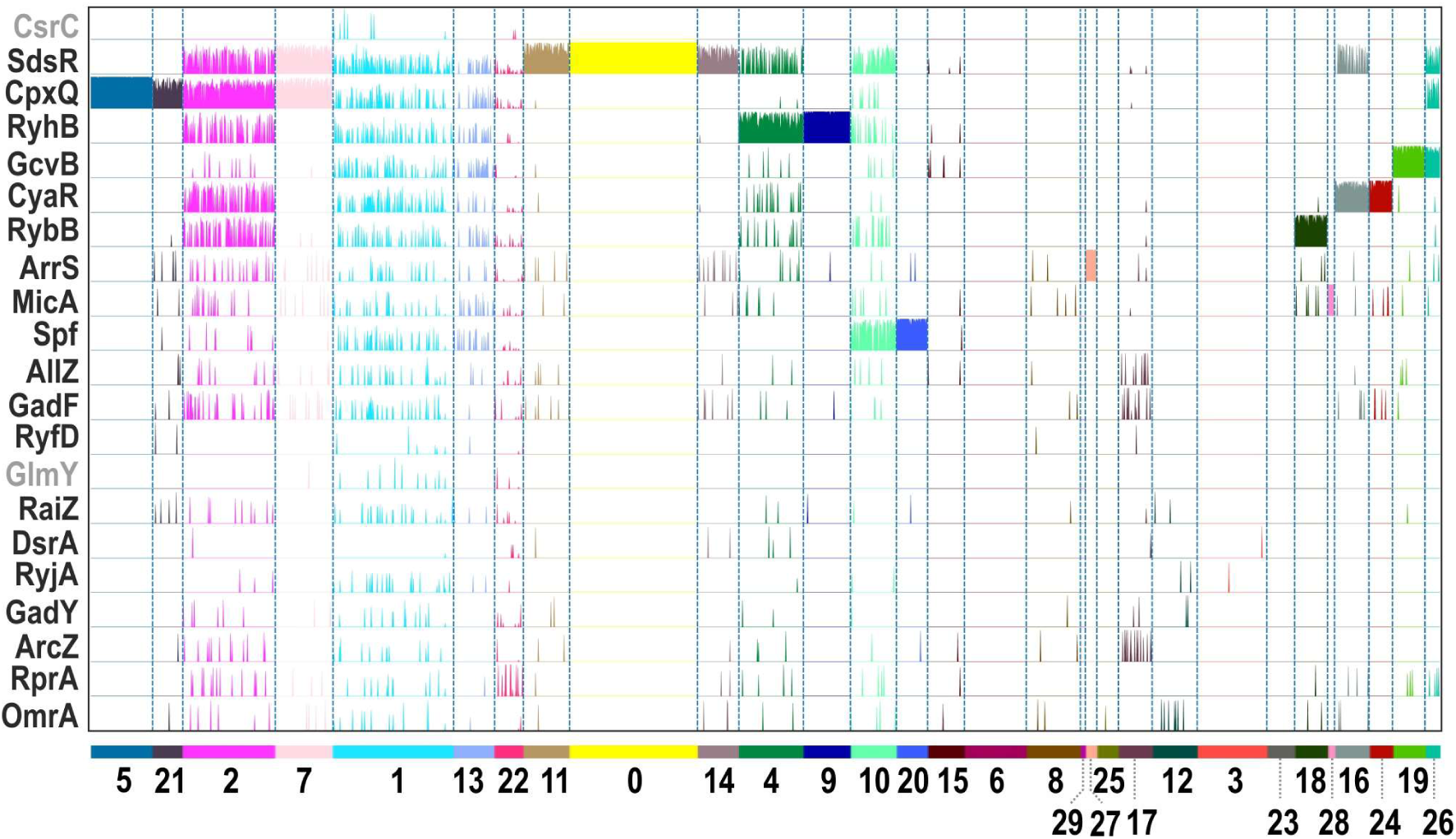
Cluster-by-cluster interaction fingerprints of *E. coli* sRNAs. Heat map summarizing the cluster-level enrichment of 18 major sRNAs across the 30 t-SNE clusters defined in Fig. 4a. Rows are sRNAs; columns are clusters (vertical dashed dividers; colour bar along the bottom labels cluster IDs). For each sRNA–cluster pair, the bar height/intensity reflects the aggregated enrichment of interactions between the sRNA and RNAs in that cluster (see Methods). Distinct, largely non-overlapping vertical patterns reveal hub sRNA–anchored modules (e.g., RyhB with cluster 9, RybB with cluster 18, MicA with cluster 28). Protein-sequestering sRNAs (e.g., CsrC, GlmY) show minimal enrichment across clusters, consistent with their lack of RNA targets. This fingerprinting illustrates the modularity of the bacterial RNA regulatory network.

**Supplementary Figure 22.**
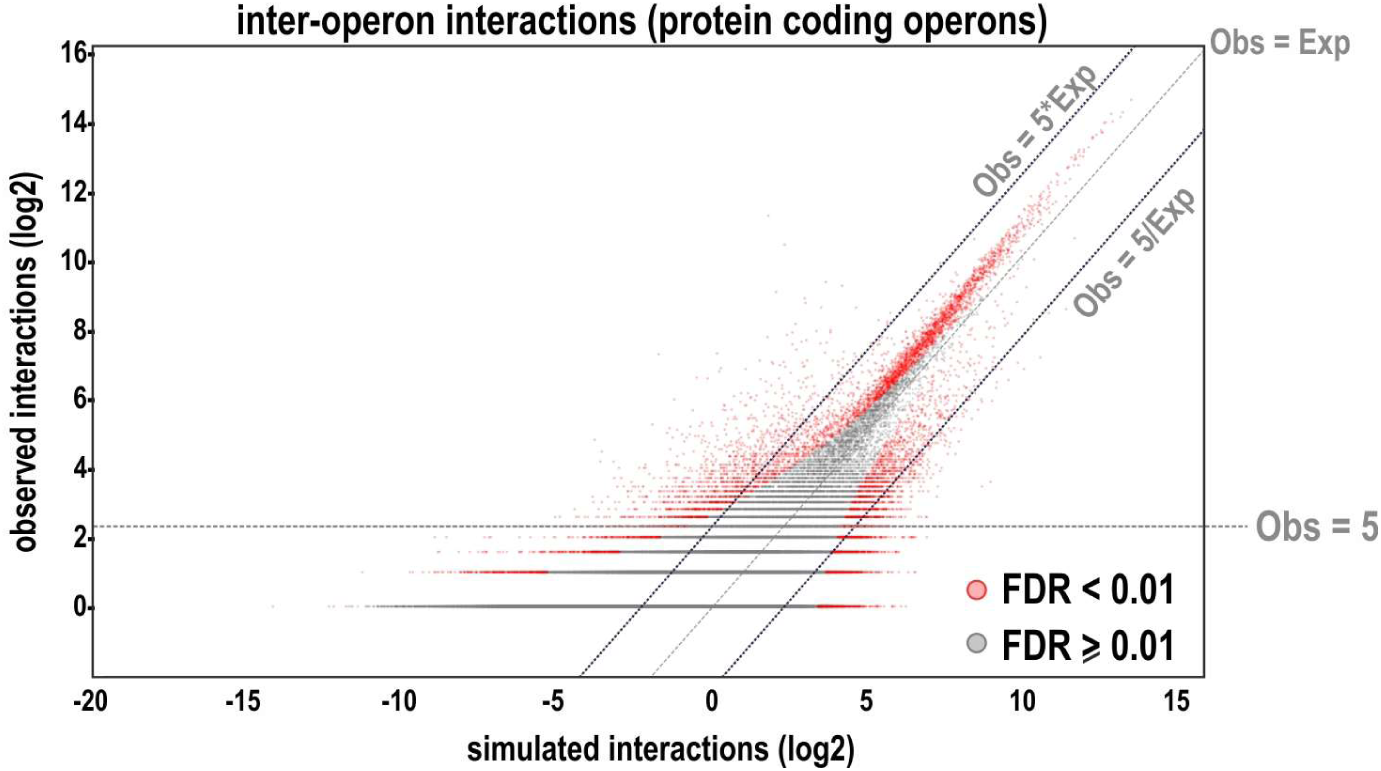
Statistical framework for inter-operonic trans interaction. An observed-vs-simulated plot for interactions between protein-coding operons. 1 million interactomes generated by degree preserving configuration-model was used for simulated data. 753 operonic transcripts from start codon of the first gene to stop codon of the last gene were included and sRNA transcripts were excluded from the analysis.

**Supplementary Figure 23.**
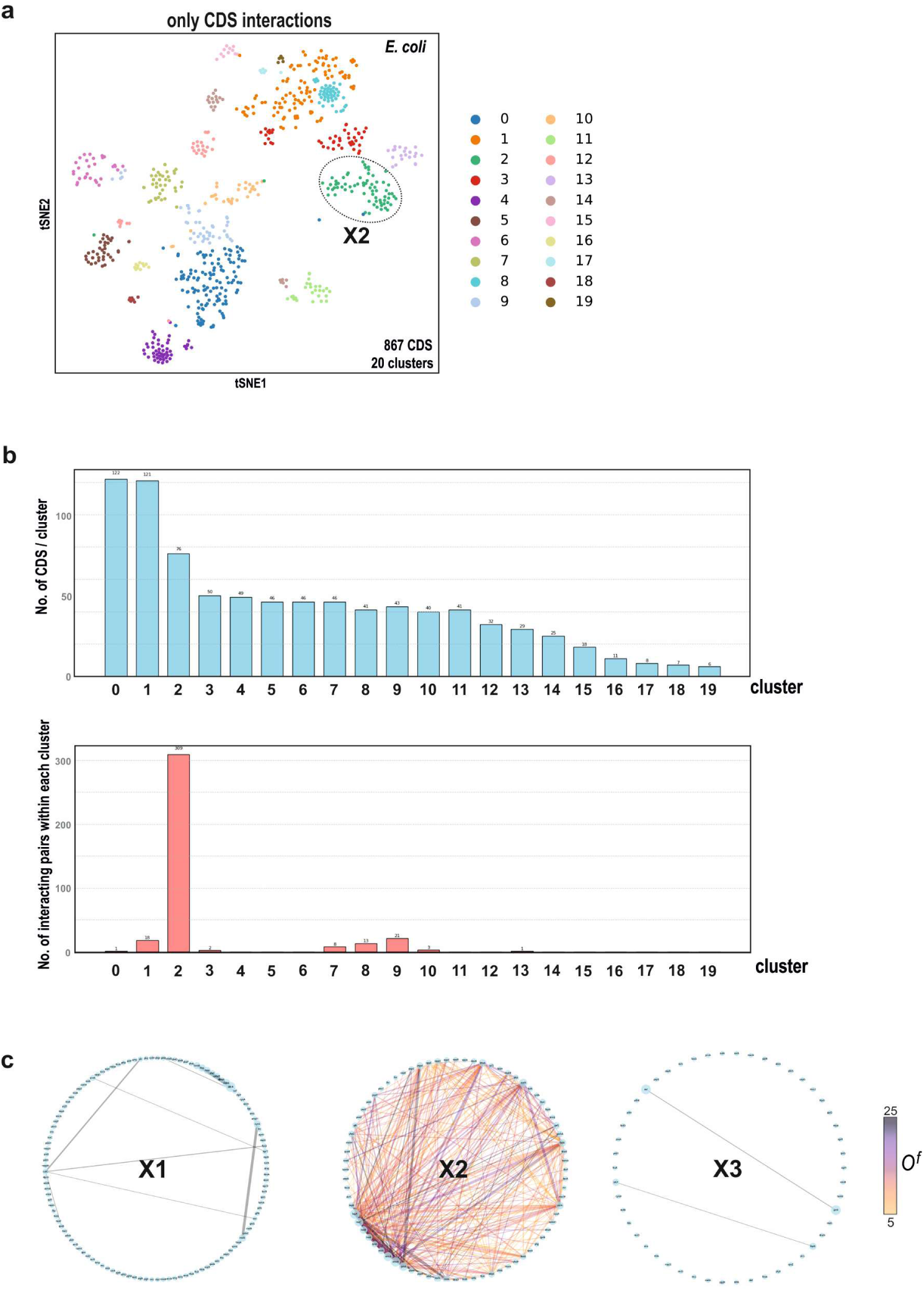
Clustering of the CDS-only interactome reveals a densely self-interacting module. **a**, t-SNE embedding constructed using only CDS–CDS long-range interactions (867 CDSs; 20 clusters). Cluster X2 (circled) corresponds to the C13 module in the full interactome (Fig. 4a), indicating that this community persists when sRNA edges are removed. **b,** Quantification of cluster properties. Top, number of CDSs per cluster. Bottom, number of within-cluster interacting pairs - Cluster X2 exhibits markedly higher internal connectivity than all other clusters. **c,** Circular network layouts for three representative clusters (X1, X2, X3). Nodes are CDSs; edges denote significant CDS–CDS contacts (edge colour encodes *O^f^*; scale at right). Cluster X2 forms a dense hairball of interactions, in stark contrast to the sparse networks of X1 and X3.

**Supplementary Figure 24.**
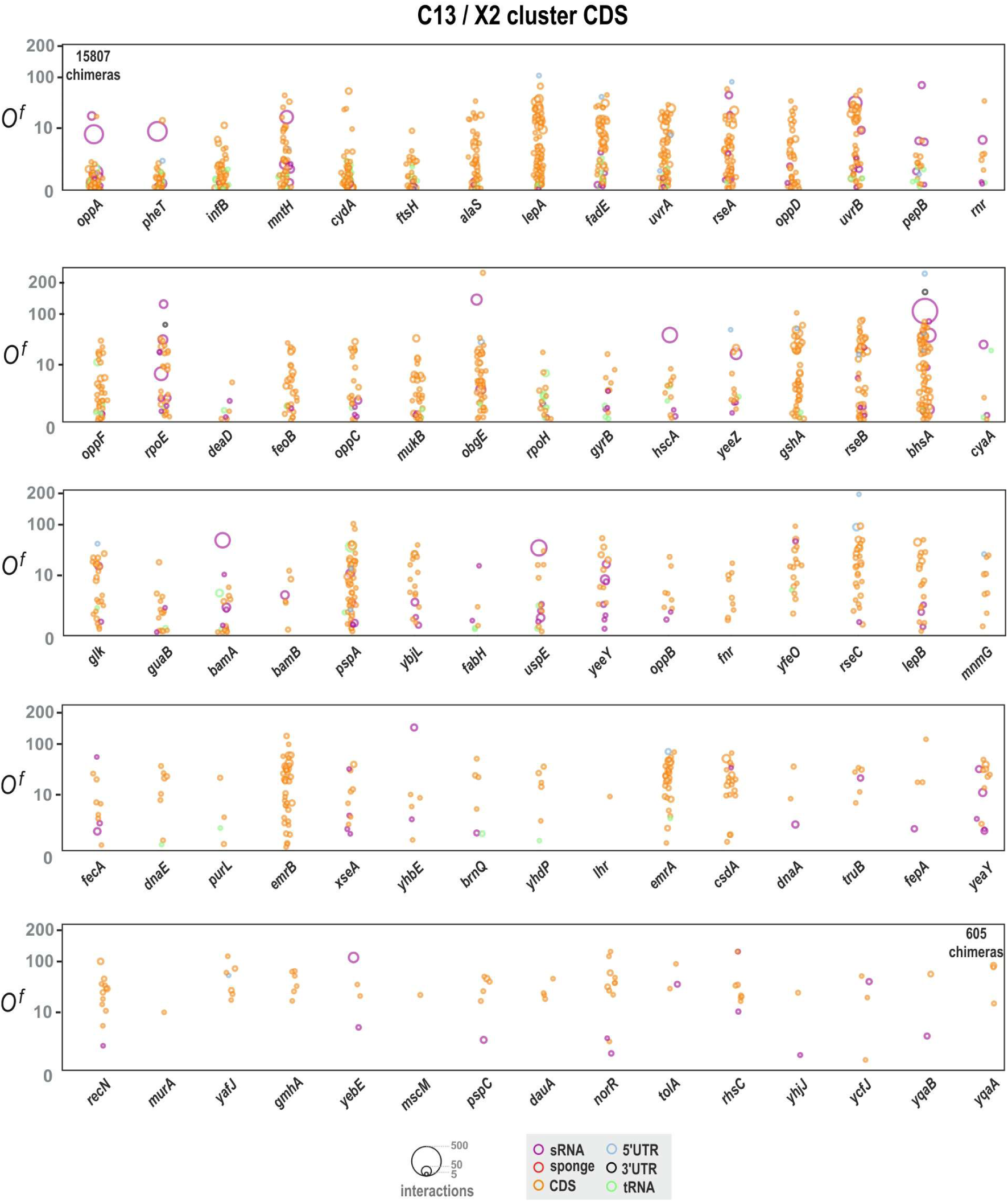
Collapsed interaction maps for all CDSs within the C13/X2 cluster. A series of collapsed interaction maps (csMaps) for CDS members of the C13/X2 cluster. The plots are arranged in descending order based on the total number of chimeric reads associated with each query CDS. The maps show that the interaction partners for these CDSs are overwhelmingly other CDSs. This contrasts with the sRNA-dominated interaction profiles seen in other clusters. A representative subset of 15 csMaps from this series is shown in Fig. 5b.

**Supplementary Figure 25.**
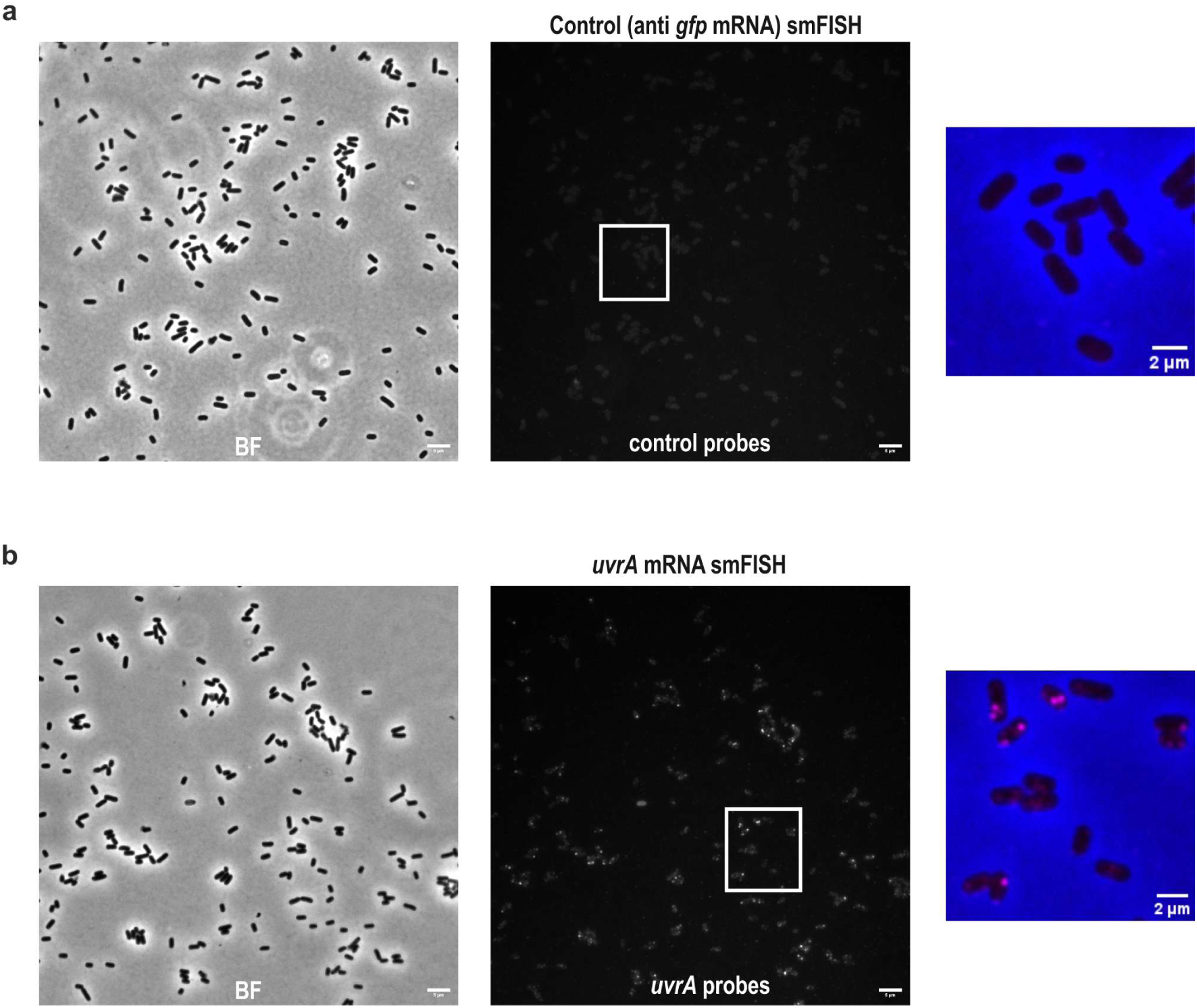
smFISH controls for Figure 5e. **a**, Negative-control smFISH with probes against *gfp* mRNA in wild-type *E. coli* (no *gfp* gene). Images were acquired and processed identically to (b). The fluorescence channel shows only diffuse background signal and a complete absence of the discrete spots seen in panel (b), confirming the high specificity of the hybridization protocol and probes. **b**, smFISH targeting *uvrA* mRNA reveals multiple discrete cytoplasmic foci of varying intensities. Dim spots are consistent with single transcripts, whereas brighter puncta indicate local multimeric aggregates. White boxes mark regions enlarged at right; in merged views the brightfield image is false-coloured blue for cellular context. Scale bars: 5 µm (wide-field), 2 µm (zoom).

**Supplementary Figure 26.**
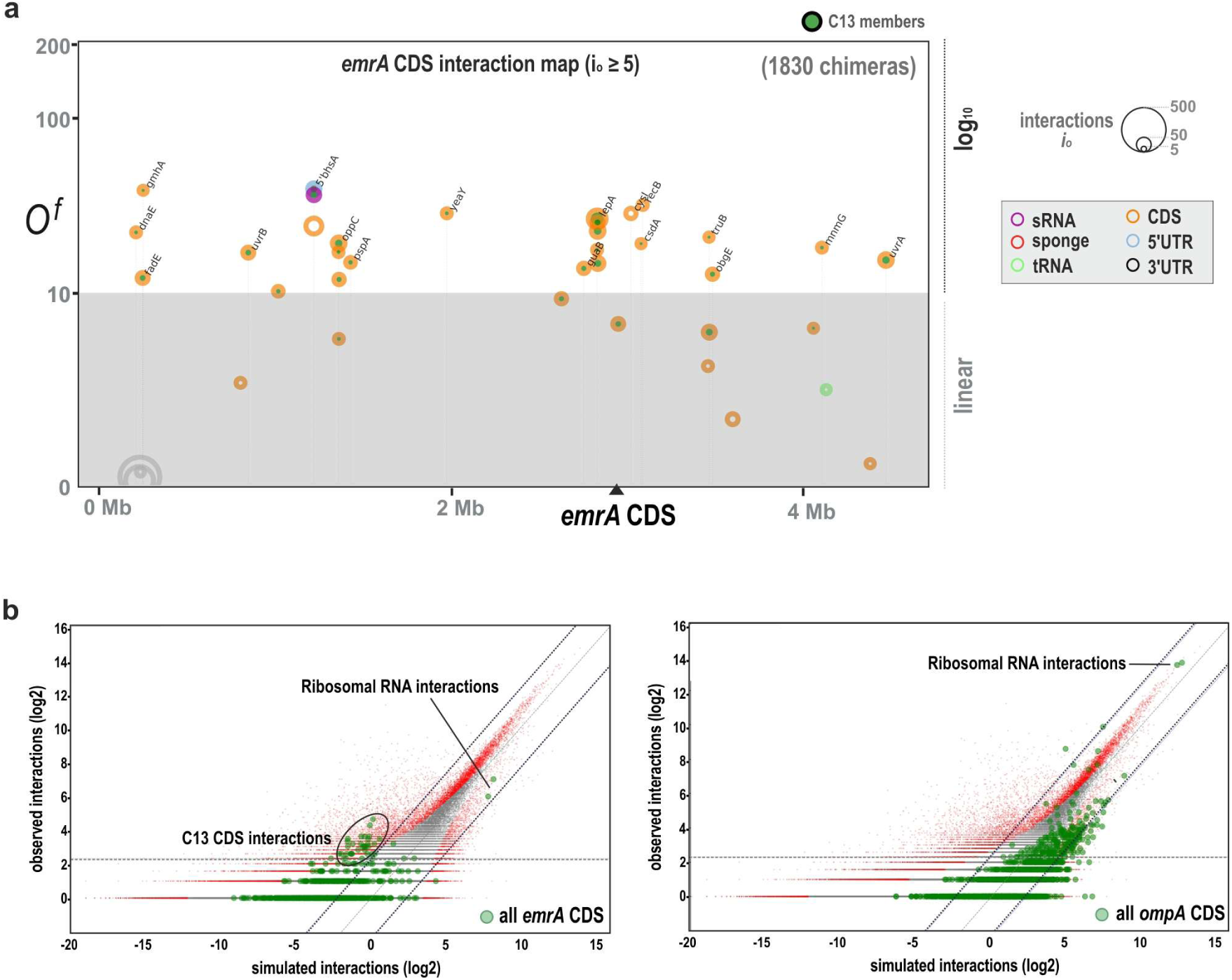
The distinct interactome of the C13 member *emrA*. **a**, Global interaction map for the *emrA* CDS. The plot confirms that its interaction partners are almost exclusively other members of the C13 stress-induced module. **b**, Observed-vs-simulated plots comparing the complete interactomes of *emrA* (left) and a control mRNA, *ompA* (right). The plot for *emrA* shows a distinct enrichment for interactions with other C13 CDSs (circled), while interactions with ribosomal RNAs are not enriched. In contrast, the interactome of the highly translated *ompA* mRNA is dominated by interactions with ribosomal RNAs. This comparison highlights the differential engagement of C13 members with the translational machinery.

**Supplementary Figure 27.**
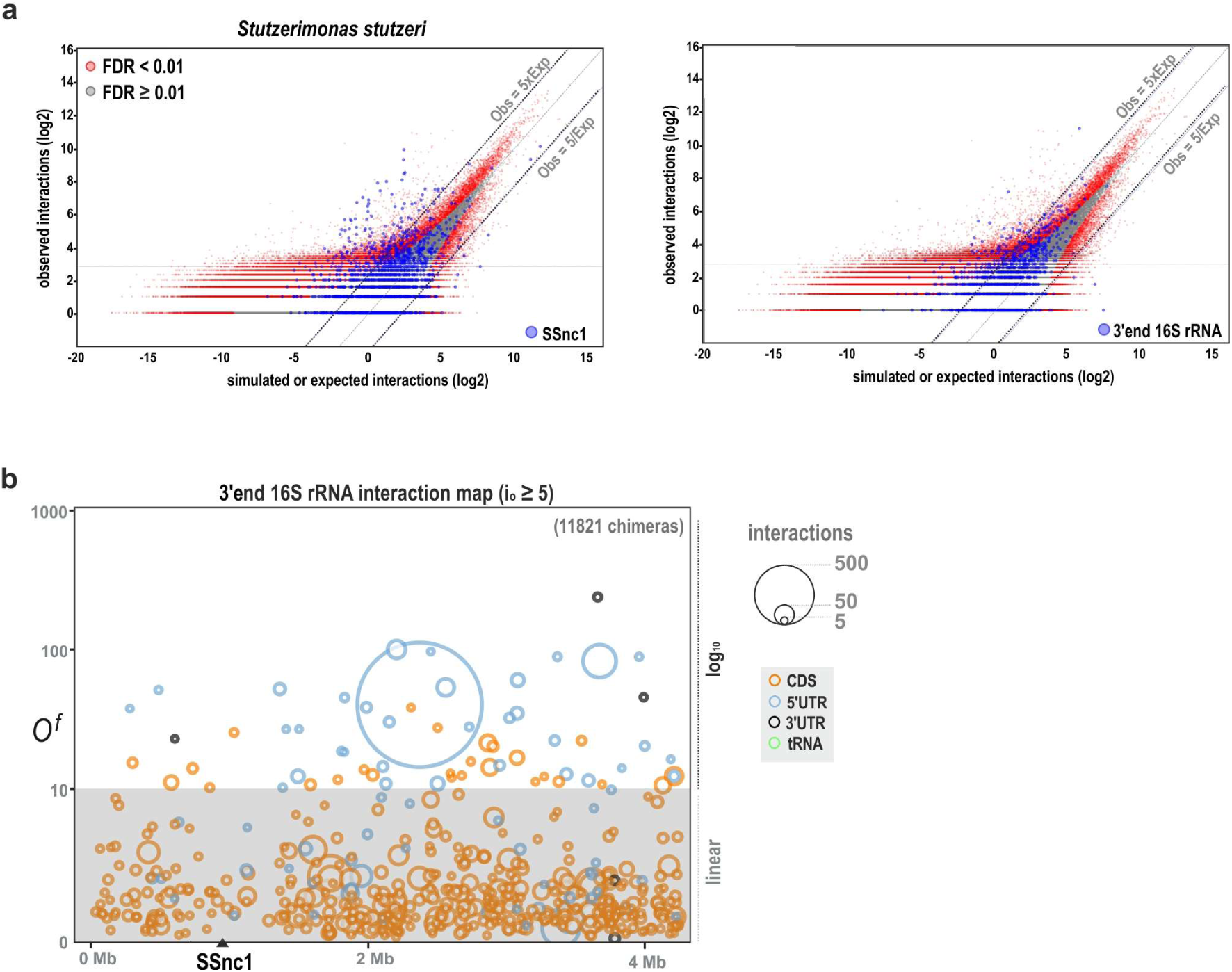
Statistical identification of significant interactions in *Stutzerimonas stutzeri*. **a**, Scatter plots of observed versus simulated interaction counts (log₂ scale) for all RNA-RNA pairs in *S. stutzeri*. Most interactions fall along the diagonal (Obs = Exp), representing the stochastic background, while statistically significant interactions (red dots, FDR < 0.01) are highly enriched. In the left panel, interactions involving the newly identified sRNA, SSnc1, are highlighted in blue, showing their strong enrichment. In the right panel, the RNA-RNA interaction pairs involving the 3’ extension of the 16S rRNA are highlighted in blue. **b**, Global interaction map for the 3′ extension of 16S rRNA in *S. stutzeri* (i_o_ ≥ 5; 11821 chimeras). Circles represent partners by genomic coordinate (x) and enrichment odds ratio *O^f^* (y); area ∝ i_o_ and colour encodes RNA feature. As in *E. coli*, the 3′ tail engages a distinct set of 5’UTRs, supporting a conserved ribosome-tethering mechanism.

**Supplementary Figure 28.**
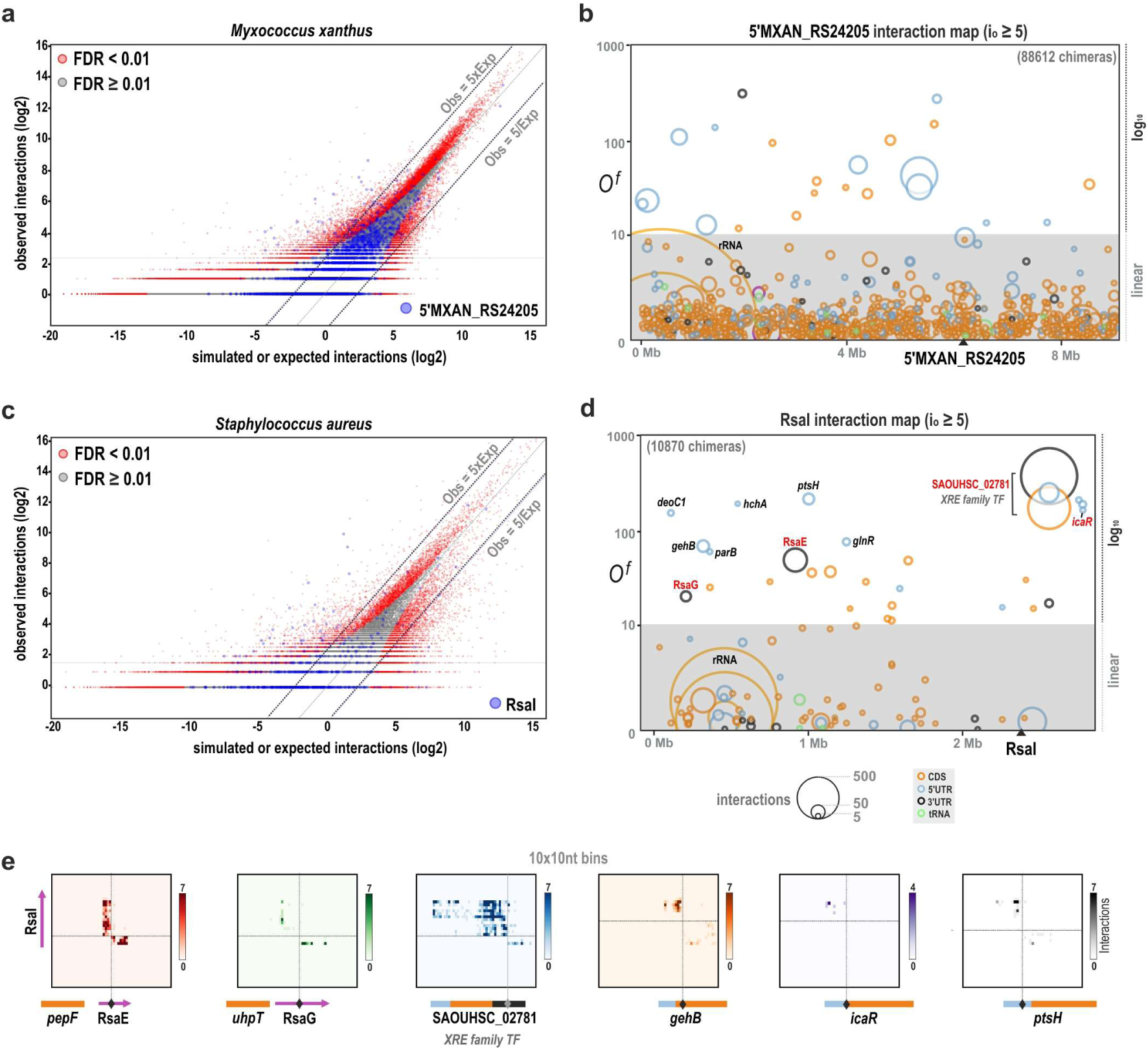
Statistical identification of significant interactions in *Myxococcus xanthus* and *Staphylococcus aureus*. **a**, Scatter plots of observed versus simulated interaction counts for all RNA-RNA pairs in *M. xanthus*. In the left panel, interactions involving the newly identified sRNA (near 5’MXAN_RS24205), are highlighted in blue. **b**, Global interaction map for the potential sRNA originating from the 5’MXAN_RS24205 locus in *M. xanthus*, showing its numerous high-confidence mRNA targets. **c**, A scatter plot of observed versus simulated interaction counts for all RNA-RNA pairs in *S. aureus*. Interactions involving the sRNA RsaI are highlighted in blue, demonstrating its significant enrichment over the stochastic background. **d**, Global interaction map for RsaI sRNA in *S. aureus*. The target RNAs previously discovered using RsaI pull-down using MAPS^60^ have been marked in red. RsaG and RsaE were identified as the 3’UTRs of *uhpT* and *pepF*, respectively as sRNAs were not included in the annotation file used for *S. aureus* TRIC-seq data analysis. e, Inter-RNA heat maps (10 × 10-nt bins) showing RsaI interactions with several targets identified in the global interaction map.

**Supplementary Table 1.** TRIC-seq Library Statistics. Summary of sequencing and mapping statistics for all libraries.

| Library Name | Total raw reads (M) | Reads after preprocessing (M) | % Mapped reads | STAR Mapper Long-range chimeras (M) | CRSSANT Short-range chimeras (M)<sup>#</sup> | Total chimeras (M) | % Chimeric reads |
| --- | --- | --- | --- | --- | --- | --- | --- |
| *E. coli* (12 libraries)<sup>†</sup> | 411.93 | 221.84 | 91.33% | 7.41 | 7.41 | 14.82 | 6.50% |
| R17S1 | 61.04 | *Early stationary (OD 2.0)* |  |  |  |  |  |
| R17S2 | 21.65 | *Stationary (OD 2.0 + 3 hrs)* |  |  |  |  |  |
| R17S4 | 23.93 | *Late stationary / overnight (OD 2.0 + 12 hrs)* |  |  |  |  |  |
| R20S1 | 39.13 | *Stationary # Replicate 1* |  |  |  |  |  |
| R20S2 | 38.13 | *Stationary # Replicate 2* |  |  |  |  |  |
| R20S3 | 43.89 | *Stationary (No MNase digestion)* |  |  |  |  |  |
| R20S4 | 41.24 | *Stationary (40 min MNase digestion)* |  |  |  |  |  |
| R20S21 | 38.46 | *Stationary (24 hrs on LB agar plate)* |  |  |  |  |  |
| R14S1 | 14.02 | *Stationary (RNase T excluded)* |  |  |  |  |  |
| R14S2 | 15.36 | *Stationary* |  |  |  |  |  |
| R14S3 | 17.18 | *Stationary* |  |  |  |  |  |
| R14S7 | 57.90 | *Stationary (rRNA not removed)* |  |  |  |  |  |
| R20S1_re*<sup>‡</sup> | 518.19 | 218.93 | 94.01% | 7.96 | - | 7.96 | 3.64% |
| R20S2_re*<sup>‡</sup> | 200.99 | 87.90 | 93.30% | 2.74 | - | 2.74 | 3.11% |
| *S. aureus* * (R22S14) | 267.35 | 124.43 | 82.93% | 12.70 | - | 12.70 | 10.21% |
| *S. aureus* grown for 30 hrs on TSA agar plate. Lysostaphin included during permeabilization |  |  |  |  |  |  |  |
| *M. xanthus* * (R22S7) | 484.93 | 229.62 | 95.83% | 11.81 | - | 11.81 | 5.14% |
| *M. xanthus* grown for 48 hrs on CTT agar plate |  |  |  |  |  |  |  |
| *S. stutzeri* * (R20S9) | 417.87 | 186.95 | 93.30% | 6.76 | - | 6.76 | 3.62% |
| *S. stutzeri* grown for 24 hrs on LB agar plate |  |  |  |  |  |  |  |

\*These libraries were sequenced on an Illumina NovaSeq X (2x150 bp). All other libraries were sequenced on an Illumina NextSeq 500 (2x75 bp).
<sup>†</sup>Statistics for the 12 *E. coli* NextSeq libraries that were combined for the analysis presented in the figures. Short-range chimeras were identified using CRSSANT<sup>75</sup>.
<sup>‡</sup>Resequenced to evaluate TRIC-seq replicates. Only used in Supplementary Fig. 8b,c and Supplementary Fig. 9a.
<sup>#</sup>For all species other than *E. coli*, only chimeras from STAR mapper, primarily (Long-range) are reported.

## Supplementary Data Legends

The externally hosted datasets comprise

**(i)** Species-specific chimeric-junction tables (Supplementary Data 1a–1e; BED-like, tab-delimited)
**(ii)** species-specific configuration-model interaction calls (Supplementary Data 2a–2d; CSV). File names and concise legends are given below. Full column definitions are included here and mirrored in the Figshare record. Genomic coordinates are 1-based and feature names follow the unified annotation used in this study (Methods).

**Supplementary Data 1** | Chimeric junction (BED-like, tab-delimited files)

Each row records a single RNA–RNA chimera detected. For *E. coli*, both junctions called by STAR along with short-range junctions called by CRSSANT are reported separately. For the other species, only long-range junctions called by STAR are reported. This minimal BED-like layout is used to report the interaction calls in Supplementary Data 2. These chimeras were used to generate all contact maps (inta, inter, composite) in this study.

*Columns:*

**A. ref_accession** - RefSeq accession of the genome build used (NC_000913.3 for *E. coli*).

**B. Pos1** - genomic coordinate of chimera end 1.

**C. Pos2** - genomic coordinate of chimera end 2.

Supplementary_Data_1a_Ecoli_STAR_chimeras.bed - *E. coli* chimeras (STAR). Supplementary_Data_1b_Ecoli_CRSSANT_chimeras.bed - *E. coli* chimeras (CRSSANT). Supplementary_Data_1c_Stutzerimonas_stutzeri_STAR_chimeras.bed - *S. stutzeri* chimeras (STAR). Supplementary_Data_1d_Myxococcus_xanthus_STAR_chimeras.bed - *M. xanthus* chimeras (STAR). Supplementary_Data_1e_Staphylococcus_aureus_STAR_chimeras.bed - *S. aureus* chimeras (STAR).

**Supplementary Data 2** | *trans* RNA–RNA interaction analysis using configuration model (CSV files)

Complete lists of long-range RNA–RNA interaction pairs and statistics from the degree-preserving configuration model (1,000,000 randomizations; FDR by Benjamini–Hochberg). These tables were used to generate global and condensed interaction maps in this study.

*Columns:*

**A. ref** - Name of the first RNA feature in the pair.

**B. target** - Name of the second RNA feature in the pair.

**C. interaction_count** - Number of unique chimeric reads supporting the pair.

**D. total_ref** - Total chimeras involving ref across the dataset.

**E. total_target** - Total chimeras involving target across the dataset.

**F. enrichment_score** - Normalized score for local (e.g., intra-operonic) interactions (distinct from odds_ratio).

**G. ref_type** - Feature type of ref (CDS, 5′UTR, 3′UTR, hkRNA, tRNA, ncRNA, sponge).

**H. target_type** - Feature type of target.

**I. expected_count** - Mean expected count for the pair under the configuration-model null.

**J. odds_ratio** - Observed/expected; specificity metric for long-range enrichment.

**K. start_ref, L. end_ref** - Genomic span of ref.

**L. start_target, N. end_target** - Genomic span of target.

**O. p_value_FDR** - FDR-corrected P-value for the long-range enrichment.

Supplementary_Data_2a_Ecoli_interactions.csv - *E. coli* interaction calls. Supplementary_Data_2b_Stutzerimonas_stutzeri_interactions.csv - *S. stutzeri* interaction calls. Supplementary_Data_2c_Myxococcus_xanthus_interactions.csv - *M. xanthus* interaction calls. Supplementary_Data_2d_Staphylococcus_aureus_interactions.csv - *S. aureus* interaction calls.

## Methods

### Bacterial strains and growth conditions

*Escherichia coli* strain BW25113 (used for all *E. coli* experiments) was grown in LB liquid media or on LB agar plates at 37°C. *Stutzerimonas stutzeri* (strain DSM10701) cultures were grown on LB agar plates at 37°C. *Staphylococcus aureus* (strain NCTC8325) was grown on tryptic soy agar plates at 37°C, and *Myxococcus xanthus* (strain DK1622) was grown on CTT agar plates at 30°C.

### TRIC-seq protocol

#### Construction of TRIC-seq libraries

TRIC-seq library preparation comprises in-situ fixation and permeabilisation, RNA fragmentation, 3′- end pCp-biotin labelling, proximity ligation, RNA purification, enrichment of chimeric junctions, and strand-specific library preparation for sequencing. Library-specific, optional step differences are listed in Supplementary Table 1.

#### Cell fixation, permeabilisation and RNA fragmentation

For each library, 2 mL of stationary-phase culture was harvested by centrifugation and resuspended in 1 mL of 1× PBS. For cells grown on solid agar, a sterile cotton swab was used to collect biomass into 1 mL of 1× PBS. Methanol fixation was performed by slowly adding 4 mL of 100% methanol (final 80%) while mixing to avoid precipitation, followed by 10 min at room temperature (RT) on an orbital shaker. Cells were collected, washed once with 1× PBS, resuspended in 5 mL of 3% formaldehyde (in 1× PBS) and incubated for 1 h at 4 °C on a rotor. Formaldehyde was removed by three washes with 2 mL of 1× PBS. Cell pellets were flash-frozen in liquid nitrogen and stored at −80 °C for up to one month.

For DSP fixation, pellets were thawed on ice and resuspended in 1 mL of 1× PBS containing 3 mM DSP, incubated 30 min at 30 °C with gentle shaking, and quenched with glycine (final 125 mM) for 10 min at RT. Cells were resuspended in 200 µL TES (10 mM Tris-HCl pH 7.5, 1 mM EDTA, 100 mM NaCl) and permeabilised using Ready-Lyse Lysozyme Solution (Biosearch Technologies, #R1804M) at 10 U/µL for 30 min at RT with gentle shaking. For *S. aureus*, lysostaphin (50 µg/mL; Sigma, #SAE0091) plus Ready-Lyse (10 U/µL) was used in TS buffer (10 mM Tris-HCl pH 7.5, 100 mM NaCl).

After permeabilisation, cells were washed three times in 1× PNK (50 mM Tris-HCl pH 7.5, 10 mM MgCl₂, 0.2% Triton X-100). An aliquot was inspected microscopically to confirm intact cell outlines. For RNA fragmentation, cells were incubated in 200 µL of 1× MNase mix (50 mM Tris-HCl pH 7.9, 5 mM CaCl₂, 0.01 gel U/µL Micrococcal Nuclease; NEB #M0247) at 37 °C. To obtain a broad fragment-size distribution, aliquots (25 µL) were removed at 5–40 min (5-min intervals) and each quenched with 25 µL of 2× quench (100 mM Tris-HCl pH 7.5, 40 mM EGTA, 0.4% Triton X-100). Quenched aliquots were pooled and washed twice with 1× PNK.

#### In situ pCp–biotin labelling and proximity ligation

These steps followed RIC-seq¹⁰ with minor adaptations. Fragmented RNAs were first treated with FastAP Thermosensitive Alkaline Phosphatase (10 U; Thermo Fisher #EF0651) in 100 µL 1× FastAP buffer for 15 min at 37 °C. Cells were washed twice with 1× quench, twice with high-salt buffer (5× PBS, 0.2% Triton X-100), and three times with 1× PNK. Cells were then resuspended in 100 µL ligation mix (10 µL 10× T4 RNA Ligase buffer, 4 µL 1 mM pCp-biotin [Jena Bioscience #NU-1706-BIO], 50 U T4

RNA Ligase 1 [Thermo Fisher #EL0021], 4 µL 40 U/µL RiboLock RNase Inhibitor [Thermo Fisher #EO0381], 50 µL 30% PEG-20000, water) and incubated overnight at 16 °C to label 3′-hydroxyl ends. FastAP treatment was repeated to remove the 3′-phosphate from the biotin moiety. Next, T4 Polynucleotide Kinase treatment (Thermo Fisher #EK0032; 100 µL: 10 µL 10× PNK buffer, 10 µL 10 mM ATP, 10 µL PNK, water) for 45 min at 37 °C phosphorylated 5′ ends. Reactions were stopped by washing twice with 1× PNK + 20 mM EGTA, then three times with 1× PNK.

For in-situ proximity ligation, permeabilised cells were resuspended in 100 µL ligation mix (10 µL 10× T4 RNA Ligase buffer, 5 µL 10 U/µL T4 RNA Ligase, 10 µL 1 mg/mL BSA, 4 µL 40 U/µL RiboLock, water) and incubated 72 h at 16 °C with gentle shaking. To maintain activity, 20 U T4 RNA Ligase, 2 µL 10 mM ATP and 1 µL RiboLock were added at 24 h and 48 h. Cell integrity was re-checked microscopically.

#### RNA purification and enrichment of chimeric junctions

After ligation, cells were pelleted and crosslinks reversed with Proteinase K at 65 °C for 3 h. Total RNA was extracted using phenol:chloroform:isoamyl alcohol (PCI) (RNA grade; Carl Roth #X985.3), collecting the aqueous phase and precipitating overnight at −20 °C with 30 volumes ethanol : 1 volume 3 M sodium acetate (pH 5.2). Pellets were collected at 20,000×g (30 min, 4 °C) and resuspended in 20 µL nuclease-free water, then DNase I treated (NEB #M0303) and re-purified by PCI/ethanol precipitation. Optionally, RNase T (NEB #M0265; 1× NEBuffer 4; 2 h, 25 °C) was used to clip residual 3′ pCp-biotin from non-ligated ends while preserving internal chimeras; RNA was re-purified. Optional rRNA depletion used the NEB bacterial rRNA Depletion Kit (#E7850S) per manufacturer’s instructions.

RNA was resuspended in 6 µL water and briefly fragmented (add 4 µL 5× NEBNext First-Strand Synthesis Reaction Buffer; 94 °C for 7 min). Biotinylated chimeras were captured on Dynabeads MyOne Streptavidin C1 (Thermo Fisher #65001) as in RIC-seq, washed stringently, eluted, and purified (PCI + ethanol).

#### Strand-specific library preparation and sequencing

Enriched, fragmented chimeras were quantified (NanoDrop). Directional libraries were prepared with NEBNext Ultra II Directional RNA Library Prep for Illumina (NEB #E7760; dUTP method, random hexamers) and sequenced as 2×75 bp (Illumina NextSeq) or 2×150 bp (Illumina NovaSeq) paired-end reads.

#### TRIC-seq data analysis

The computational analysis of TRIC-seq data involves several key stages: (i) generation of a comprehensive genome annotation; (ii) mapping of sequencing reads to identify both local and long-range chimeric interactions; (iii) statistical analysis to identify significant interactions; and (iv) visualization of both RNA structures and interaction networks.

#### Genome Annotation and Feature Definition

To enable an unbiased, transcriptome-wide analysis, a custom, comprehensive genome annotation was first generated from the standard GFF3 annotation file for each bacterium. A custom Python script was used to ensure the entire genome was covered by defined features. The script first parsed standard features (CDS, tRNA, rRNA). For *E. coli*, known ncRNAs from the EcoCyc database^76^ were also included at this stage. For the other bacteria, only the sRNA/ncRNA annotations in the GFF3 file were included.

For all species, intergenic gaps were then systematically addressed. Intergenic gaps <40 bp were closed by extending the smaller flanking gene; for larger gaps, strand-aware 5′ and 3′UTR features were defined from flanking gene boundaries, allowing intergenic sRNA discovery in non-models. To avoid mapping ambiguity from multiple rRNA operons, all but the first copy of each major rRNA (5S, 16S, 23S) were masked in the reference FASTA. Every nucleotide was thereby assigned to a feature.

#### Mapping and Filtering of Chimeric Reads

Raw FASTQ files were pre-processed using fastp^77^ (version 1.0.1) for adapter trimming, quality filtering, PCR duplicate removal, and merging of overlapping read pairs (--overlap_len_require 18). All processed reads (merged and unmerged) were then aligned to the custom genome index using STAR^78^ (version 2.7.11a) in chimeric detection mode (--chimOutType Junctions), with parameters optimized to capture RNA chimeras (--chimSegmentMin 13, --chimJunctionOverhangMin 15). This generated a Chimeric.out.junction file which primarily reported long-range (*trans*) interactions and an Aligned.out.sam file. The Aligned.out.sam file was subsequently processed with the CRSSANT pipeline to extract chimeras (primarily short-range) from gapped reads^75^.

#### Gini Coefficient Analysis

To quantify the heterogeneity of interaction distributions along a transcript, which serves as a proxy for its structural state, a Gini coefficient was calculated for each annotated CDS and ncRNA feature. The analysis was based exclusively on long-range interactions to specifically probe the overall folding state rather than local contacts. For each gene, a per-nucleotide interaction profile was generated by counting the number of chimeric reads where one end mapped to a given nucleotide within the gene, and the other end mapped outside a ±5 kb flank of that gene’s coordinates.

The Gini coefficient was then calculated from this per-nucleotide interaction profile. The calculation is based on the Lorenz curve, which plots the cumulative fraction of interactions against the cumulative fraction of nucleotides. The Gini coefficient represents the degree of inequality in the interaction distribution; a value of 0 indicates perfect equality (a uniform distribution of interactions across the transcript), while a value approaching 1 indicates maximum inequality (interactions concentrated in very few locations). This analysis was performed for all annotated CDS and ncRNA features to generate the dataset used for the scatter plot shown in Fig. 1c and the distributions in Supplementary Fig. 2. The set of 200 high-TE genes used for comparison were from Burkhardt *et al.*^24^; top 50 absolute protein-synthesis genes (Suppl. Fig. 2; highlighted in Fig. 3g) were from Li *et al*.²⁵.

#### Identification of accessible regions from long-range Interactions

To identify accessible, likely single-stranded regions on abundant RNAs such as the 16S rRNA, a per-nucleotide structural profile was generated from long-range interaction data. The underlying premise is that the frequency of long-range inter-molecular ligation at a nucleotide serves as a proxy for its *in vivo* accessibility and single-stranded nature. For each target RNA, all chimeric reads were filtered to keep only those in which one end mapped to the RNA and the other end mapped ≥ 5 kb away on the genome. A per-nucleotide interaction profile was then built by counting these filtered long-range chimeras at every position. This raw profile was smoothed with a 3-nt sliding-window average, and local maxima—representing the most frequently ligated and therefore most accessible sites—were identified with the *find_peaks* function in SciPy. Enrichment of these maxima in accessible regions (single stranded bases and bases at the base of loops/bulges) versus stem interiors was assessed with a one-tailed hypergeometric test under a uniform-random placement model.

#### Statistical framework for *trans*-Interactions

To distinguish specific, biologically relevant interactions from the background of stochastic collisions inherent in a crowded cellular environment, a statistical framework based on a configuration model was implemented. This model was chosen because it generates the most appropriate null hypothesis for this type of data. It preserves the overall interaction frequency (or “degree”) of each individual RNA, thereby accounting for confounding factors like its abundance and general chemical propensity to interact. This allows for a more accurate assessment of whether an observed interaction frequency between two specific RNAs is significantly greater than what would be expected by chance alone.

For the analysis, a null distribution was generated by performing 1 million Monte Carlo randomisations, where interaction partners were randomly shuffled while preserving the degree of each RNA. The observed number of interactions between any two RNAs was then compared to this simulated distribution to calculate an odds ratio (*O^f^*) and a p-value. The p-values were corrected for multiple hypothesis testing using the Benjamini-Hochberg method to obtain a false discovery rate. Unless stated otherwise, a fixed enrichment cut-off of *O^f^* ≥ 10 together with a minimum support of io ≥5 was applied when reporting trans-regulatory interactions, reflecting the GcvB benchmarking (Fig. 2c; Supplementary Fig. 9b).

#### Benchmarking of sRNA target prediction

To validate the performance of TRIC-seq for sRNA target identification, the GcvB regulon was used as a benchmark. The set of 56 experimentally validated GcvB targets compiled by Miyakoshi *et al.* were used as true positives; all other genes with ≥ 5 GcvB interactions formed the negative set (Fig. 2c). For the ROC curve analysis (Fig. 2c), the highest *O^f^* across any feature of a gene was used as its score. This analysis was performed on the set of all genes that formed at least one chimeric read with GcvB.

To assess the effect of sequencing depth, performance metrics (sensitivity, specificity and precision) were computed across interaction-count thresholds while holding the odds-ratio cut-off fixed at *O^f^* ≥ 10. For each integer threshold io from 1 to 50, a gene was called a predicted positive if it had at least one GcvB interaction meeting *O^f^* ≥ 10 and supported by ≥ io chimeric reads; metrics were then calculated against the 56 validated targets. Specificity rapidly approached ∼1 and precision saturated with only a few supporting reads, indicating that few chimeras suffice for high-confidence calls; higher thresholds mainly reduced sensitivity with little gain in specificity (Supplementary Fig. 9b).

#### Global and Collapsed interaction maps

To visualize the interaction partners of a specific RNA, two types of plots were used. The Global Interaction Map plots the partners of a query RNA along the genome, where each partner or target is a circle, whose position indicates its genomic coordinate and odds ratio (*O^f^*), size indicates interaction count (io), and colour indicates feature type. The RIL-seq defined targets highlighted in global maps were obtained from stationary phase Hfq RIL-seq library in *E. coli* hosted at RILseqDB^23^. For intergenic RIL-seq targets, the flanking genes were both considered target for comparison with TRIC-seq. The

Collapsed interaction map (csMap) allows for the comparison of multiple query RNAs by collapsing the genome onto a categorical x-axis. To avoid overplotting in dense regions, this script implements a local filtering, displaying only the interaction with the highest *O^f^* within any 1000-bp window.

#### RNA Structure and contact heat maps

Intramolecular contact maps were generated by binning a gene sequence (2 to 30 nt resolution) and counting interactions per bin pair, followed by ICE (iterative correction and eigenvector decomposition) normalization^79^. Inter-RNA heat maps visualized the interaction between two RNAs by binning a window (e.g., ± 200 nt) around each RNA. Intra-Inter heat maps simultaneously visualized internal structure (negative values, blue) and intermolecular contacts (positive values, red) on a concatenated axis. Predicted duplexes and hybridisation energies were computed with IntaRNA ^74^.

#### Operon metaplot analysis

To analyse average interaction patterns across operon boundaries (Fig. 1b), TRIC-seq contacts from consecutive CDS pairs were aggregated. Only boundaries with ≥ 50 inter-gene chimeras were included (n = 773). Each pair of adjacent CDSs was scaled to 100 bins per CDS (200×200 matrix total). Each matrix was normalised by its total contacts (equal weight per boundary) and averaged to produce the metaplot.

#### TRIC-seq structural comparison with Crystal Structures

To benchmark the accuracy of TRIC-seq, contact maps for the 16S rRNA, 23S rRNA, and tmRNA were compared against their known crystal structure (PDB ID: 7ACJ). Chimeric reads were binned (5 nt) and ICE-normalized. A distance matrix was computed from the 3D coordinates of the phosphate backbone atoms for each corresponding bin. For validation, an ROC analysis was performed, where a “true positive” was defined as a pair of bins with a distance ≤ 25 Å. The AUC was then calculated to quantify how well TRIC-seq contacts recapitulated known spatial proximities.

#### Minimum Free Energy (MFE) analysis

To assess the thermodynamic stability of the observed interactions between the 16S rRNA 3’ tail and 5’UTRs, a minimum free energy (MFE) analysis was performed. All annotated 5’UTR sequences were extracted from the *E. coli* genome. The MFE of duplex formation between the 19-nucleotide sequence of the 16S rRNA 3’ tail extension (5’-CACCTCCTTACCTTAAAGA-3’) and the last 20 nucleotides of each 5’UTR was calculated using the RNA.duplexfold function from the ViennaRNA package^80^. For a negative control, the same analysis was performed using a scrambled version of the 16S 3’ tail sequence (5’- CACCTACACATTCTGCAAG-3’). The distributions of MFE values for all 5’UTRs were then compared to the distributions for specific subsets using a two-sided Mann-Whitney U (MWU) test to determine statistical significance.

#### Network clustering and visualization

Unsupervised clustering of RNA features was performed using the scanpy Python library. A sparse interaction matrix was constructed from long-range interactions, filtered to include interactions with a pair-wise interaction count of at least 5 for *E. coli* dataset. The data was normalized and log-transformed, and PCA was used for dimensionality reduction. A neighborhood graph (n_neighbors=15, n_pcs=15) was computed and Leiden clustering applied. Embeddings were visualised with t-SNE.

#### Single-Molecule mRNA FISH

To validate the *in vivo* co-aggregation of transcripts identified in Cluster 13, smFISH was performed. A set of 48 unique 20-nt fluorescent oligonucleotide probes were designed for *uvrA* (labelled with Atto565) and *fadE* (labelled with Atto488) and were manufactured by SynBio. For the experiment, *E. coli* cells were fixed for 30 minutes with 4% paraformaldehyde at room temperature. The fixed cells were collected by centrifugation and washed twice with PBS. Permeabilization was achieved through a sequential ethanol treatment: first, a 1-hour incubation in 70% ethanol, followed by a 1-hour incubation in 100% ethanol at room temperature. After permeabilization, the cells were air-dried on the bench for 1 hour, washed once with 2x SSC buffer, and resuspended in a hybridization buffer (2x SSC, 25% formamide, 0.1% SDS). The fluorescently labelled DNA oligo probe sets were then added to a final concentration of 10 ng/µL each, and hybridization was carried out overnight at 37°C. The following day, the cells were washed three times for 1 hour each at 42°C with the hybridization buffer. After the final wash, cells were resuspended in 2x SSC and mounted on a slide with a 1% agarose pad (in 2xSSC) for visualization. Imaging was performed using an Olympus BX fluorescence microscope.

#### Strain construction and Western Blot Analysis

To validate the regulatory effects of the sRNA CpxQ on its newly identified targets, *psiF* and *gcd*, Western blot analysis was performed. First, the endogenous *psiF* and *gcd* genes in *E. coli* were tagged with a 3xFLAG epitope at their C-termini using the lambda red recombineering protocol. The 3xFLAG tag was introduced as part of a chloramphenicol resistance cassette with 50 bp homology arms for targeted integration. A CpxQ deletion strain (Δ*cpxQ*) was also constructed using lambda red recombineering, replacing the *cpxQ* gene with a kanamycin resistance cassette. For overexpression experiments, the *cpxQ* gene was cloned into a vector under the control of a pTAC promoter, which also contained the *lacI* repressor gene and an ampicillin resistance marker. To collect samples, the CpxQ overexpression strain was grown to late-exponential phase, and expression was induced with 100 µM IPTG for 1 hour. For Western blotting, bacterial samples corresponding to an OD₆₀₀ of 1.0 were collected and boiled in 100 µL of protein sample loading buffer containing DTT. A 5 µL aliquot of this lysate, corresponding to 0.05 OD units of cells, was loaded onto a 15% SDS-PAGE gel. The separated proteins were then transferred to a nitrocellulose membrane using a semi-wet transfer apparatus. The FLAG-tagged proteins were detected using a primary Monoclonal ANTI-FLAG M2 antibody produced in mouse (Sigma, #F3165), followed by a secondary IRDye 680RD Donkey anti-Mouse IgG antibody (LI-COR, #926-68072). As a loading control, biotinylated proteins were detected simultaneously using IRDye 800CW Streptavidin (LI-COR, #926-32230). The fluorescent signals from both channels were detected using a LI-COR Odyssey Fc Imager and quantified using ImageJ.

## Data availability

All TRIC-seq libraries used this study have been deposited in GEO repository under accession number GSE305265. Processed datasets (Supplementary Data 1: chimeric-junction tables; Supplementary Data 2: configuration-model interaction calls) for all four microbes are hosted at Figshare (https://figshare.com/s/341c66f6ebbaae99f3b0). Source data for main figures is provided separately.

## Code availability

All custom scripts used in this study are available on GitHub. Analysis pipelines and utilities are available at the TRIC-seq repository (https://github.com/gdx7/TRIC-seq), and visualisation scripts (global/condensed interaction maps and intra-/inter-RNA heat maps) are available at TRICweb repository (https://github.com/gdx7/TRICweb). A live web app is hosted at www.TRICseq.com for interactive visualisation of datasets from all four microbes generated in this study.

## Acknowledgements

I thank Tjalling J. Siersma, Dennis Rijnsburger and Gideon Meerhoff for their assistance in setting up the drna lab; Selina van Leeuwen (MAD, University of Amsterdam) and Biomarker Technologies (BMK) GmbH for high-quality sequencing services. I thank Rabea Wagner, Frans van der Kloet, Biwen Wang, Alberto Agnolin, Jurian Wijnheijmer, Fleur van de Koolwijk, Michael Lekkerkerker, Marine Kleiber, Selina Forrer, Alberto Pavan, Vera van Melis, Nils Meiresonne and Dolça Serra Sallent for helpful discussions and support. I am thankful to Leendert Hamoen for providing a bench space at the outset of this project and comments on the manuscript. Large language models (Gemini 2.5 Pro and OpenAI o1) were used to assist in developing custom analysis and visualisation scripts. This work was funded by a VENI grant (VI.VENI.192.103) and a VIDI grant (VI.VIDI.223.132) to G.D. from the Dutch Research Council (NWO).

## References

1. Storz, G., Vogel, J. & Wassarman, K. M. Regulation by Small RNAs in Bacteria: Expanding Frontiers. Molecular Cell vol. 43 Preprint at 10.1016/j.molcel.2011.08.022 (2011).

2. Protter, D. S. W. & Parker, R. Principles and Properties of Stress Granules. Trends Cell Biol 26, 668–679 (2016).

3. Jain, S. et al. ATPase-Modulated Stress Granules Contain a Diverse Proteome and Substructure. Cell 164, 487–498 (2016).

4. Muthunayake, N. S., Tomares, D. T., Childers, W. S. & Schrader, J. M. Phase-separated bacterial ribonucleoprotein bodies organize mRNA decay. Wiley Interdiscip Rev RNA 11, e1599 (2020).

5. Al-Husini, N. et al. BR-Bodies Provide Selectively Permeable Condensates that Stimulate mRNA Decay and Prevent Release of Decay Intermediates. Mol Cell 78, 670–682.e8 (2020).

6. Weng, X. et al. Spatial organization of RNA polymerase and its relationship with transcription in *Escherichia coli*. Proceedings of the National Academy of Sciences 116, 20115–20123 (2019).

7. Sarfatis, A., Wang, Y., Twumasi-Ankrah, N. & Moffitt, J. R. Highly multiplexed spatial transcriptomics in bacteria. Science 387, eadr0932 (2025).

8. Moffitt, J. R., Pandey, S., Boettiger, A. N., Wang, S. & Zhuang, X. Spatial organization shapes the turnover of a bacterial transcriptome. Elife 5, (2016).

9. Lu, Z. et al. RNA Duplex Map in Living Cells Reveals Higher-Order Transcriptome Structure. Cell 165, (2016).

10. Cao, C., et al. Global in situ profiling of RNA-RNA spatial interactions with RIC-seq. Nature Protocols vol. 16 Preprint at 10.1038/s41596-021-00524-2 (2021).

11. Cai, Z. et al. RIC-seq for global in situ profiling of RNA–RNA spatial interactions. Nature 582, (2020).

12. Wu, T. et al. KARR-seq reveals cellular higher-order RNA structures and RNA-RNA interactions. Nat Biotechnol 42, 1909–1920 (2024).

13. Bartholomäus, A. et al. Bacteria differently regulate mRNA abundance to specifically respond to various stresses. Philos Trans A Math Phys Eng Sci 374, (2016).

14. Bremer, H. & Dennis, P. P. Modulation of Chemical Composition and Other Parameters of the Cell at Different Exponential Growth Rates. EcoSal Plus 3, (2008).

15. Melamed, S. et al. Mapping the small RNA interactome in bacteria using RIL-seq. Nat Protoc 13, (2018).

16. Iosub, I. A. et al. Hfq CLASH uncovers sRNA-target interaction networks linked to nutrient availability adaptation. Elife 9, (2020).

17. Liu, F. et al. In vivo RNA interactome profiling reveals 3’UTR-processed small RNA targeting a central regulatory hub. Nat Commun 14, 8106 (2023).

18. Carrier, M.-C., Lalaouna, D. & Massé, E. A game of tag: MAPS catches up on RNA interactomes. RNA Biol 13, 473–6 (2016).

19. Mustoe, A. M. et al. Pervasive Regulatory Functions of mRNA Structure Revealed by High-Resolution SHAPE Probing. Cell 173, 181–195.e18 (2018).

20. Hsieh, T. H. S., Fudenberg, G., Goloborodko, A. & Rando, O. J. Micro-C XL: Assaying chromosome conformation from the nucleosome to the entire genome. Nat Methods 13, (2016).

21. Gavrilov, A. A. et al. Elementary 3D organization of active and silenced E. coli genome. Nature (2025) doi:10.1038/s41586-025-09396-y.

22. Kuchina, A. et al. Microbial single-cell RNA sequencing by split-pool barcoding. Science 371, (2021).

23. Melamed, S. et al. Global Mapping of Small RNA-Target Interactions in Bacteria. Mol Cell 63, 884–97 (2016).

24. Burkhardt, D. H. et al. Operon mRNAs are organized into ORF-centric structures that predict translation efficiency. Elife 6, (2017).

25. Li, G.-W., Burkhardt, D., Gross, C. & Weissman, J. S. Quantifying absolute protein synthesis rates reveals principles underlying allocation of cellular resources. Cell 157, 624–35 (2014).

26. Szabo, Q., Bantignies, F. & Cavalli, G. Principles of genome folding into topologically associating domains. Sci Adv (2019) doi:10.1126/sciadv.aaw1668.

27. Rouskin, S., Zubradt, M., Washietl, S., Kellis, M. & Weissman, J. S. Genome-wide probing of RNA structure reveals active unfolding of mRNA structures in vivo. Nature 505, 701–5 (2014).

28. Lalaouna, D., Eyraud, A., Devinck, A., Prévost, K. & Massé, E. GcvB small RNA uses two distinct seed regions to regulate an extensive targetome. Mol Microbiol 111, (2019).

29. Sharma, C. M. et al. Pervasive post-transcriptional control of genes involved in amino acid metabolism by the Hfq-dependent GcvB small RNA. Mol Microbiol 81, (2011).

30. Miyakoshi, M. et al. Mining RNA-seq data reveals the massive regulon of GcvB small RNA and its physiological significance in maintaining amino acid homeostasis in *Escherichia coli*. Mol Microbiol (2021) doi:10.1111/MMI.14814.

31. Sharma, C. M., Darfeuille, F., Plantinga, T. H. & Vogel, J. A small RNA regulates multiple ABC transporter mRNAs by targeting C/A-rich elements inside and upstream of ribosome-binding sites. Genes Dev 21, 2804–2817 (2007).

32. Weilbacher, T. et al. A novel sRNA component of the carbon storage regulatory system of Escherichia coli. Mol Microbiol 48, 657–70 (2003).

33. Göpel, Y., Papenfort, K., Reichenbach, B., Vogel, J. & Görke, B. Targeted decay of a regulatory small RNA by an adaptor protein for RNase E and counteraction by an anti-adaptor RNA. Genes Dev 27, 552–64 (2013).

34. Chao, Y. & Vogel, J. A 3’ UTR-Derived Small RNA Provides the Regulatory Noncoding Arm of the Inner Membrane Stress Response. Mol Cell 61, (2016).

35. Grabowicz, M., Koren, D. & Silhavy, T. J. The Cpxq sRNA negatively regulates skp to prevent mistargeting of β-barrel outer membrane proteins into the cytoplasmic membrane. mBio 7, (2016).

36. Babitzke, P. & Romeo, T. CsrB sRNA family: sequestration of RNA-binding regulatory proteins. Curr Opin Microbiol 10, 156–63 (2007).

37. Holmqvist, E. et al. Global RNA recognition patterns of post-transcriptional regulators Hfq and CsrA revealed by UV crosslinking in vivo. EMBO J 35, 991–1011 (2016).

38. Baussier, C., Oriol, C., Durand, S., Py, B. & Mandin, P. Small RNA OxyS induces resistance to aminoglycosides during oxidative stress by controlling Fe-S cluster biogenesis in Escherichia coli. Proc Natl Acad Sci U S A 121, e2317858121 (2024).

39. Battesti, A., Majdalani, N. & Gottesman, S. The RpoS-Mediated General Stress Response in *Escherichia coli*. Annu Rev Microbiol 65, 189–213 (2011).

40. Sedlyarova, N. et al. sRNA-Mediated Control of Transcription Termination in E. coli. Cell 167, (2016).

41. Badaoui, A. et al. Ribose lowers RpoS translation through RbsD mRNA. Preprint at 10.1101/2023.08.28.555190 (2023).

42. Denham, E. L. The Sponge RNAs of bacteria - How to find them and their role in regulating the post-transcriptional network. Biochim Biophys Acta Gene Regul Mech 1863, 194565 (2020).

43. M, M., Y, C. & J, V. Cross talk between ABC transporter mRNAs via a target mRNA-derived sponge of the GcvB small RNA. EMBO J 34, 1478–1492 (2015).

44. Adams, P. P. et al. Regulatory roles of Escherichia coli 5’ UTR and ORF-internal RNAs detected by 3’ end mapping. Elife 10, (2021).

45. Lalaouna, D. et al. A 3’ external transcribed spacer in a tRNA transcript acts as a sponge for small RNAs to prevent transcriptional noise. Mol Cell 58, (2015).

46. Melamed, S., Adams, P. P., Zhang, A., Zhang, H. & Storz, G. RNA-RNA Interactomes of ProQ and Hfq Reveal Overlapping and Competing Roles. Mol Cell 77, (2020).

47. Papenfort, K., Bouvier, M., Mika, F., Sharma, C. M. & Vogel, J. Evidence for an autonomous 5′ target recognition domain in an Hfq-associated small RNA. Proceedings of the National Academy of Sciences 107, 20435–20440 (2010).

48. Waters, L. S., Sandoval, M. & Storz, G. The Escherichia coli MntR miniregulon includes genes encoding a small protein and an efflux pump required for manganese homeostasis. J Bacteriol 193, 5887–97 (2011).

49. Wang, H. et al. Increasing intracellular magnesium levels with the 31-amino acid MgtS protein. Proc Natl Acad Sci U S A 114, 5689–5694 (2017).

50. Jensen, P. A. Ten species comprise half of the bacteriology literature, leaving most species unstudied. Preprint at 10.1101/2025.01.04.631297 (2025).

51. Lv, F., et al. Integrated Hfq-interacting RNAome and transcriptomic analysis reveals complex regulatory networks of nitrogen fixation in root-associated *Pseudomonas stutzeri* A1501. mSphere 9, (2024).

52. Goldman, B. S. et al. Evolution of sensory complexity recorded in a myxobacterial genome. Proceedings of the National Academy of Sciences 103, 15200–15205 (2006).

53. Yu, Y. T. N., Yuan, X. & Velicer, G. J. Adaptive evolution of an sRNA that controls myxococcus development. Science vol. 328 Preprint at 10.1126/science.1187200 (2010).

54. Geisinger, E., Adhikari, R. P., Jin, R., Ross, H. F. & Novick, R. P. Inhibition of *rot* translation by RNAIII, a key feature of *agr* function. Mol Microbiol 61, 1038–1048 (2006).

55. Bohn, C., Rigoulay, C. & Bouloc, P. No detectable effect of RNA-binding protein Hfq absence in Staphylococcus aureus. BMC Microbiol 7, 10 (2007).

56. Bronesky, D., et al. *Staphylococcus aureus* RNAIII and Its Regulon Link Quorum Sensing, Stress Responses, Metabolic Adaptation, and Regulation of Virulence Gene Expression. Annu Rev Microbiol 70, 299–316 (2016).

57. McKellar, S. W. et al. RNase III CLASH in MRSA uncovers sRNA regulatory networks coupling metabolism to toxin expression. Nat Commun 13, 3560 (2022).

58. Gudeta, D. D., Lei, M. G. & Lee, C. Y. Contribution of *hla* Regulation by SaeR to Staphylococcus aureus USA300 Pathogenesis. Infect Immun 87, (2019).

59. Wang, K. et al. GehB Inactivates Lipoproteins to Delay the Healing of Acute Wounds Infected with Staphylococcus aureus. Curr Microbiol 81, 36 (2024).

60. Bronesky, D. et al. A multifaceted small RNA modulates gene expression upon glucose limitation in Staphylococcus aureus. EMBO J 38, (2019).

61. Das, S. et al. Natural mutations in a *Staphylococcus aureus* virulence regulator attenuate cytotoxicity but permit bacteremia and abscess formation. Proceedings of the National Academy of Sciences 113, (2016).

62. Morrison, J. M. et al. Characterization of SSR42, a Novel Virulence Factor Regulatory RNA That Contributes to the Pathogenesis of a Staphylococcus aureus USA300 Representative. J Bacteriol 194, 2924–2938 (2012).

63. Jobson, M.-E. et al. SSR42 is a Novel Regulator of Cytolytic Activity in Staphylococcus aureus. Preprint at 10.1101/2024.07.11.603084 (2024).

64. Horn, J. et al. Long Noncoding RNA SSR42 Controls Staphylococcus aureus Alpha-Toxin Transcription in Response to Environmental Stimuli. J Bacteriol 200, (2018).

65. Homberger, C., Barquist, L. & Vogel, J. Ushering in a new era of single-cell transcriptomics in bacteria. microLife 3, (2022).

66. Pei, L. et al. Aggresomes protect mRNA under stress in Escherichia coli. Preprint at 10.1101/2024.04.27.591437 (2024).

67. Ortiz-Rodrĺguez, L. A. et al. Stress Changes the Material State of a Bacterial Biomolecular Condensate and Shifts its Function from mRNA Decay to Storage. Preprint at 10.1101/2024.11.12.623272 (2024).

68. Anderson, P. & Kedersha, N. RNA granules: post-transcriptional and epigenetic modulators of gene expression. Nat Rev Mol Cell Biol 10, 430–6 (2009).

69. Mateju, D. et al. Single-Molecule Imaging Reveals Translation of mRNAs Localized to Stress Granules. Cell 183, 1801–1812.e13 (2020).

70. Vesper, O. et al. Selective translation of leaderless mRNAs by specialized ribosomes generated by MazF in Escherichia coli. Cell 147, 147–57 (2011).

71. Kwon, D. RNA function follows form - why is it so hard to predict? Nature 639, 1106–1108 (2025).

72. Holmqvist, E. & Vogel, J. RNA-binding proteins in bacteria. Nat Rev Microbiol 16, 601–615 (2018).

73. Gerovac, M. et al. Global discovery of bacterial RNA-binding proteins by RNase-sensitive gradient profiles reports a new FinO domain protein. RNA 26, 1448–1463 (2020).

74. Mann, M., Wright, P. R. & Backofen, R. IntaRNA 2.0: enhanced and customizable prediction of RNA-RNA interactions. Nucleic Acids Res 45, W435–W439 (2017).

75. Zhang, M. et al. Classification and clustering of RNA crosslink-ligation data reveal complex structures and homodimers. Genome Res (2022) doi:10.1101/gr.275979.121.

76. Moore, L. R. et al. Revisiting the y-ome of *Escherichia coli*. Nucleic Acids Res 52, 12201–12207 (2024).

77. Chen, S., Zhou, Y., Chen, Y. & Gu, J. fastp: an ultra-fast all-in-one FASTQ preprocessor. Bioinformatics 34, i884–i890 (2018).

78. Dobin, A. et al. STAR: ultrafast universal RNA-seq aligner. Bioinformatics 29, 15– 21 (2013).

79. Imakaev, M. et al. Iterative correction of Hi-C data reveals hallmarks of chromosome organization. Nat Methods 9, 999–1003 (2012).

80. Varenyk, Y., Spicher, T., Hofacker, I. L. & Lorenz, R. Modified RNAs and predictions with the ViennaRNA Package. Bioinformatics 39, (2023).

